# Global cellular proteo-lipidomic profiling of diverse lysosomal storage disease mutants using nMOST

**DOI:** 10.1101/2024.03.26.586828

**Authors:** Felix Kraus, Yuchen He, Sharan Swarup, Katherine A. Overmyer, Yizhi Jiang, Johann Brenner, Cristina Capitanio, Anna Bieber, Annie Jen, Nicole M. Nightingale, Benton J. Anderson, Chan Lee, Joao A. Paulo, Ian R. Smith, Jürgen M. Plitzko, Steven P. Gygi, Brenda A. Schulman, Florian Wilfling, Joshua J. Coon, J. Wade Harper

## Abstract

Lysosomal storage diseases (LSDs) comprise ∼50 monogenic disorders marked by the buildup of cellular material in lysosomes, yet systematic global molecular phenotyping of proteins and lipids is lacking. We present a nanoflow-based multi-omic single-shot technology (nMOST) workflow that quantifies HeLa cell proteomes and lipidomes from over two dozen LSD mutants. Global cross-correlation analysis between lipids and proteins identified autophagy defects, notably the accumulation of ferritinophagy substrates and receptors, especially in *NPC1*^-/-^ and *NPC2*^-/-^ mutants, where lysosomes accumulate cholesterol. Autophagic and endocytic cargo delivery failures correlated with elevated lyso-phosphatidylcholine species and multi-lamellar structures visualized by cryo-electron tomography. Loss of mitochondrial cristae, MICOS- complex components, and OXPHOS components rich in iron-sulfur cluster proteins in *NPC2*^-/-^ cells was largely alleviated when iron was provided through the transferrin system. This study reveals how lysosomal dysfunction affects mitochondrial homeostasis and underscores nMOST as a valuable discovery tool for identifying molecular phenotypes across LSDs.

## INTRODUCTION

Lysosomes are the central organelle for degradative and recycling functions within eukaryotic cells, and digest cargo (e.g. macromolecules) delivered via various trafficking systems, including autophagy or endocytosis.^1,2^ Moreover, lysosomes are integral to lipid catabolism and nutrient sensing via the MTOR protein kinase. Genetic and clinical studies have linked lysosomal dysfunction to a wide variety of storage accumulation pathologies, called “lysosomal storage diseases”, or henceforth LSDs.^3^ More than 50 LSDs have been described and while individually rare, their combined prevalence is 1:5000 in live births, while some population groups carry higher incidence rates. Not surprisingly, given the central role of lysosomes in cellular health, LSDs have been linked to various human diseases, including Niemann-Pick type C1/2 (*NPC1* and *NPC2*), Gaucher (*GBA1*), Pompe (*GAA*), Danon (*LAMP2*), and Neuronal Ceroid Lipofuscinoses (*GRN*), and increased risk of Parkinson’s is observed within a subset of LSDs (*SMPD1*, *ASAH1*, *ATP13A2*, *CTSD*, *GBA1*).^4–7^ The majority of LSD genes encode catabolic enzymes (e.g. hydrolases) that function in the lysosomal lumen, although an important subset function as small molecule/ion transporters that maintain metabolic balance within the lysosome. Defects in these activities can result in a cascade of effects on downstream processes, leading to the accumulation of various types of materials: sphingolipids, mucopolysaccharides, glycoproteins, and lipofuscin.^3^ Additional storage material can also accumulate as a secondary response to the primary storage defect, including phospholipids, glycosphingolipids, and cholesterol. Several LSDs also lead to an imbalance in lipid catabolism and/or defects in sphingolipid metabolism (e.g. *CLN5* and *GRN*)^8,9^, raising the question of the extent to which dysregulation of lipid homeostasis underlies divergent LSDs.^10^

Understanding complex relationships across lipidomes and proteomes necessitates an approach for molecular profiling across the diversity of LSD loss-of-function mutations. In principle, systematic profiling should facilitate the identification of similarities and differences in molecular phenotypes, defined here as alterations in the abundance of proteins and lipids, exhibited across the various disease classes. Here, we develop a highly sensitive <u>n</u>anoflow liquid chromatography (LC) <u>M</u>ulti-<u>O</u>mic <u>S</u>ingle-shot <u>T</u>echnology (nMOST) for simultaneous analysis of lipids and proteins. nMOST represents a second generation implementation of the microflow LC MOST method (µMOST)^11^ and integrates a multi-omic sample preparation strategy^12^ with intelligent lipidomic data acquisition^8,13^ to provide significantly improved sensitivity. To demonstrate the generality of the approach and to create a resource for the community, we applied nMOST to a collection of more than two dozen LSD mutant HeLa cell lines. Global lipid-protein cross-correlation analysis reveals patterns of accumulation of individual proteins and lipids in distinct genotypes. Among the strongest correlations identified was autophagy and ferritinophagy signatures associated with *NPC1*^-/-^ and *NPC2*^-/-^ cells, consistent with previous studies,^14–16^ but unexpectedly, we also observed defects in the mitochondrial proteome, particularly in *NPC2*^-/-^ cells. *NPC1* and *2*, which are mutated in Niemann-Pick disease, play a key role in cholesterol trafficking out of the lysosome.^17^ Cholesterol esters within the lysosomal lumen are hydrolyzed by the LSD-associated protein LIPA (Lipase A), and the cholesterol product is then carried by the lumenal NPC2 protein to the membrane-embedded NPC1 transporter, facilitating cholesterol egress from the lysosome. In addition to the known accumulation of cholesterol in cells lacking *NPC1* or *NPC2*, nMOST analysis additionally revealed alterations in other lipids including Lyso-PC species, which are major membrane building blocks in cells.

Through a series of cell biological validation experiments that demonstrate the value of this approach, we demonstrate in *NPC2*^-/-^ cells that defects in autophagic turnover of ferritin – a major mechanism for cellular control of iron availability^18,19^ – lead to defects in mitochondrial cristae and oxidative phosphorylation (OXPHOS) complexes rich in iron-sulfur cluster proteins, which can be ameliorated by supplying iron through the transferrin system. Iron similarly rescued the abundance of OXPHOS machinery during differentiation of *NPC1^-/-^* or *NPC2^-/-^* mutant stem cells to induced neurons (iNeurons). Moreover, we demonstrate a defect in delivery of autophagic and endocytic cargo to the lysosomal lumen. Importantly, these defects correlate with the formation multilamellar structures within lysosomes, as visualized at nanometer resolution using cryo-electron tomography (cryo-ET), and with accumulation of Lyso-PC as a possible membrane building block. This resource of quantitative lipidomic and proteomic data across diverse classes of LSD mutations – available through an online data portal (https://coonlabdatadev.com/) – provides a rich landscape for further biological discovery.

## RESULTS

### Robust proteomic and lipidomic analysis using nMOST

We previously reported µMOST as an approach to acquire proteomics and lipidomics data simultaneously.^11^ However, the sensitivity of microflow was limited, which prompted the development of an analogous nanoflow method with substantially increased sensitivity (**Figure 1A**). nMOST takes advantage of the fact that the vast majority of lipid species elute from reverse phase columns well after the vast majority of peptides, as indicated when peptide and lipid extracts from HEK293 cells are analyzed separately with the same mobile phase gradient (**Figure 1B, and 1C**, left and middle panels). We found, however, that sequential loading of lipid and peptide extracts followed by LC-MS provided virtually identical performance (**Figure 1B and 1C**, right panels), with correlation coefficients (*r*) for both protein label free quantification (LFQ) and lipid intensity > 0.99 (**Figure 1D, Figure S1A**). Thus, the presence of peptides on the immobile phase did not affect lipid detection or quantification and vice versa. Consistent with added sensitivity of the nanoflow approach, nMOST delivered >2-fold more protein (5593 vs 2540) or >3-fold more lipid (967 vs 281) identifications as compared with µMOST when analyzing 1 μg of HEK293 cell-derived peptides or 0.03% lipid mass in a single LC-MS run (**Figure 1E**). Direct comparison between nMOST and µMOST in quantifying protein groups of sub-cellular organelles (mitochondria, lysosomes, endosomes, nucleus) across three magnitudes of sample dilution demonstrate the superiority of nMOST over previous implementations (**Figure 1E and S1B**). Moreover, the method was found to be robust, with a similar number of biomolecules quantified over an extended period of data collection, with a median relative standard deviation (RSD) of 5.0% for proteins and 13.2% for lipids (**Figure 1F**, **Figure S1C, and S1D** and see below).

**Figure 1.**
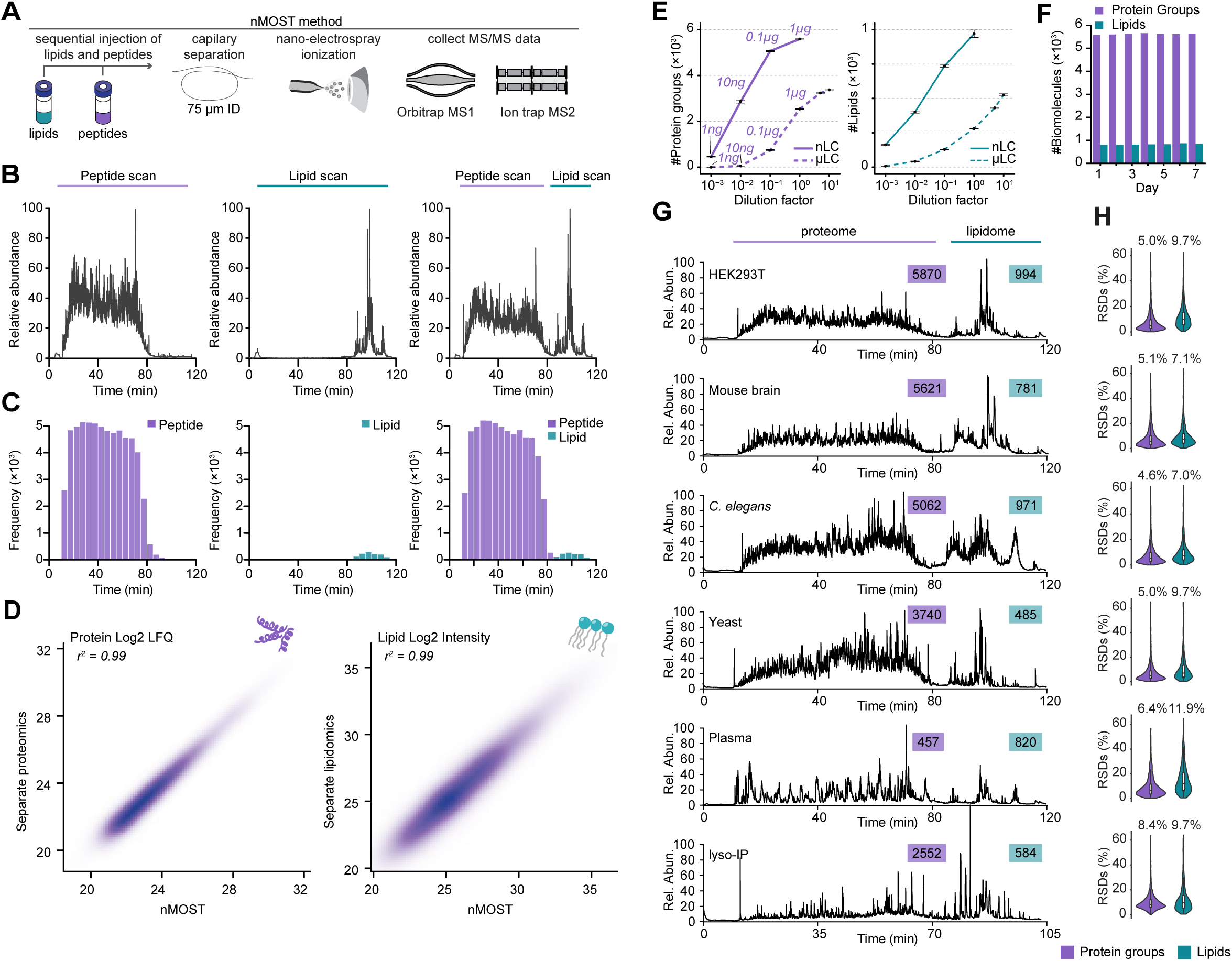
Development and benchmarking of nMOST for proteomics and lipidomics analysis. **(A)** Schematic of the nMOST method, which allows proteome and lipidome analysis by LC-MS. Lipid and protein extracts isolated from the same cell sources are sequentially injected onto LC prior to elution with an organic gradient and MS analysis (see **STAR METHODS**). **(B)** Chromatograms showing HEK293 cell peptide and lipid elution features during a 120 min gradient examining (left panel) total protein extract, (middle panel) total lipid extract, and (right panel) sequentially loaded protein and lipid extracts and nMOST analysis. The vast majority of peptides elute before 80 min while the majority of lipids elute between 80 and 120 min. **(C)** Peptide and lipid identifications from the corresponding LC-MS run in panel B. **(D)** Correlation of proteins (left panel) and lipids (right panel) identified by separate LC-MS (y-axis) versus nMOST (x-axis). r^2^ values are >0.99. **(E)** Number of protein groups and lipid groups identified by nMOST versus µMOST methods. nMOST routinely out-performed µMOST for both proteins (left panel) and lipids (right panel). Amount of peptide injections are labelled above the line for each nMOST and µMOST. **(F)** Performance was comparable for both proteins and lipids when measured daily over a 7-day acquisition period. **(G)** nMOST allows simultaneous analysis of proteins and lipids from HEK293 cells, mouse brain extracts, *C. elegans* extracts, budding yeast extracts, human plasma, and lysosomes from HeLa cells isolated by Lyso-IP. **(H)** RSD values for the data in panel G.

To demonstrate versatility, we benchmarked nMOST performance across multiple species (HEK293 cells, mouse brain, *C. elegans*, budding yeast) and sample types (cell extracts, plasma, purified lysosomes) (**Figure 1G**). We observed the expected complexity of proteomes across the various samples, and routinely detected thousands of proteins and ∼500- 1000 lipid species (**Figure 1G**), with RSDs of 4.6-8.4% for proteins and 7.0-11.9% for lipids (**Figure 1H**). The consistent identifications and stable quantifications over extended periods of analysis time highlight the robustness of the method and reinforce its potential for reproducible and high-quality data acquisition to elucidate complex relationships between proteomes and lipidomes.

### A Tool-kit for Systematic Analysis of LSD Genes

The data described so far points to the utility of the nMOST approach in quantifying alterations in lipids and proteins in a diverse spectrum of biological samples. To demonstrate the utility of this approach, we set out to generate molecular landscapes of cells that are deficient in various LSD proteins. We used CRISPR/Cas9 to attempt targeting of 52 LSD genes in HeLa cell lines containing endogenously tagged TMEM192-HA for lysosome detection (HeLa^TMEM192-HA^) (**Figure 2A, S1E, Table S1**).^20,21^ In total, we validated a total of 38 mutants across multiple functional classes of LSDs using a combination of sequencing and proteomics approaches: 23 homozygous deletions, 5 heterozygous deletions, and 10 mutants containing one or more alleles with an in-frame deletion (**Figure S1F; Table S1**). We applied nMOST to total cell extracts using quadruplicate independent cultures (**Figure 2A**). This was accomplished by running the samples across 15 sets, where each set contain multiple HeLa^TMEM192-HA^ or Control parental samples and were flanked by instrument QC runs, ensuring stable performance. In total, 318 whole cell extract samples were subjected to nMOST, representing 4 weeks of cumulative continuous data collection, with little change in method performance, as indicated by the log_2_ abundance values for proteins and lipids. Additionally, 45 QC samples spread throughout the data collection period demonstrated high performance reproducibility (**Figure S1G-I**). In total, over 7000 proteins and 2000 lipids were routinely identified and quantified in whole cell extracts (**Table S2**). In the majority of cases for heterozygous and in-frame deletion clones, the levels of target proteins detected by proteomics, when detected, indicated substantially reduced protein levels (**Figure S1F;** see **Table S1** for detailed genotyping results of each clone). We excluded heterozygous deletions in the subsequent analysis, leaving 33 LSD mutant cell lines which serve as a resource for phenotypic analysis of a broad range of LSD genes.

**Figure 2.**
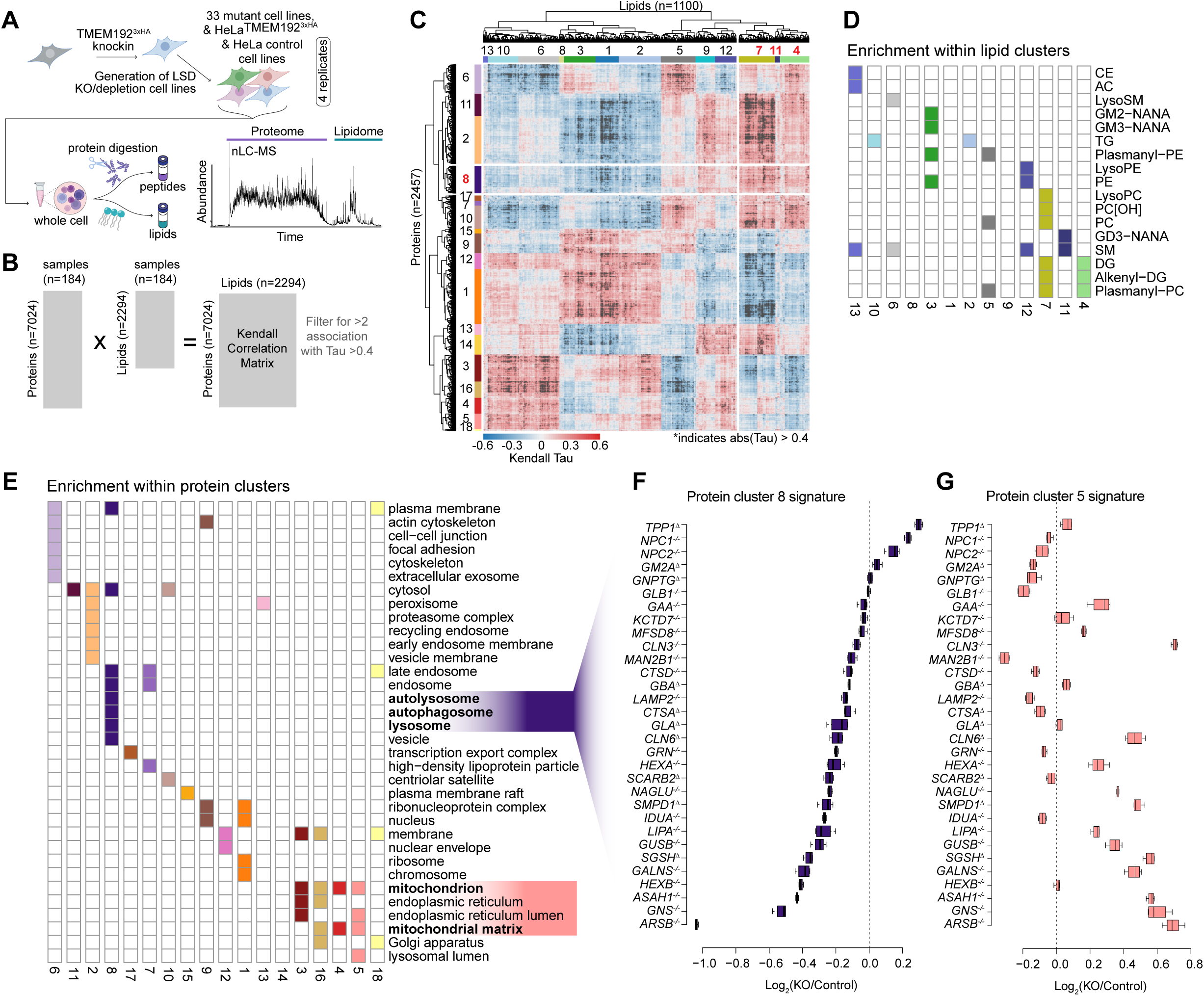
Landscape of total proteomes and lipidomes from LSD mutant cells using nMOST. **(A)** Schematic describing the method for analysis of total cell extracts across 33 LSD mutants. Protein and lipid extracts were isolated from the samples in quadruplicate, and then sequentially injected for analysis by LC-MS over a 120 min gradient. **(B,C)** Panel B is a schematic depicting the method used for lipid/protein cross-correlation analysis employing a Kendall rank correlation (filtered for >1 association with Tau >0.4). Panel C shows a heatmap for Tau values. Clusters for proteins and lipids are shown. **(D)** Schematic showing the enrichment of specific lipids within individual lipid clusters. **(E)** Schematic showing the subset of GO term Cellular Compartment enriched within individual protein clusters. **(F)** Summed protein cluster 8 signature (sum abundance of all proteins within cluster 8 (enriched for autophagy terms) across the LSD mutant cells plotted as log_2_FC (KO/WT). **(G)** Signature of protein cluster 5 (sum protein abundance relative to WT) across the LSD mutant cells.

### Molecular fingerprinting of LSDs using nMOST

To identify molecular fingerprints across the LSD mutant cells, we performed lipid-protein cross correlation analysis using a Kendall rank approach (**Figure 2B**), resulting in 1100 lipids and 2457 proteins with at least two correlations with |Tau| >0.4. Hierarchical clustering of the correlation matrix revealed 13 lipid and 18 protein clusters (**Figure 2C, Table S3**) with significant enrichment of either lipid-classes or subcellular compartments across the proteo-lipidomic landscape for 33 LSD mutant cell lines (**Figure 2D and 2E**). Given the importance of autophagy in cellular homeostasis, we focused on protein cluster 8; this cluster encompassing lysosome, autolysosome and autophagosome terms and correlated significantly with phosphocholines (PC), lysoPCs, plasmanyl-PCs, diacylglycerols (DGs), alkenyl-DGs, and gangliosides in lipid clusters 4, 7 and 11 (**Figure 2C-E**). The summed cluster 8 signature plotted as log_2_(KO/Control) indicated that *NPC1^-/-^*, *NPC2^-/-^*, and *TPP1*^-/-^ cells were among the strongest candidates for affected mutants within the lysosome cluster, cluster 8 (**Figure 2F, Table S2 and S3**). This contrasted with a variety of other LSD mutants, that were enriched in clusters 4 and 5 and displayed increased levels of mitochondrial proteins (e.g. *LIPA^-/-^*, *ARSB^-/-^*, *GNS^-/-^*) (**Figure 2G**).

To further deconvolute which organelles and processes were most affected by altered cholesterol efflux from lysosomes, we created a curated sublist of organelle proteins (1784 IDs, see **STAR METHODS**), encompassing annotated proteins for mitochondria, lysosome, endosome, Golgi, ER, proteasome and as well as autophagy and iron homeostasis and performed kmeans clustering (**Figure S1J**; contains average abundance of indicated annotation group). *NPC1^-/-^*and *NPC2^-/-^* clustered closely together (Genotype Group 2) while *LIPA^-/-^* was located in Group 5, consistent with differential effect on organelle proteomes (**Figure S1J**). We focused on three clusters of interest: While proteins belonging to k-means cluster 3 (GO: [regulation of] mitochondrial RNA catabolic process) were increased across all three genotypes, cluster 4 (GO: [macro-] autophagy & vacuole organization) was elevated in Group 2 mutants containing *NPC1*^-/-^ and *NPC2*^-/-^, but not in *LIPA^-/-^*in Group 5, while k-means cluster 5 (GO: ATP synthesis and aerobic electron transport chain) was increased in Group 5 but not Group 2 (**Figure S1J**). Thus, the signatures observed in **Figure 2C-E** for *NPC* mutants and *LIPA^-/-^*may reflect changes in organelle homeostasis, especially mitochondria and autophagy.

### A nMOST-LSD Resource for Lipid-Protein Correlation Analysis

The availability of deep proteome and lipidome data across cells lacking various LSD proteins provides an opportunity to identify common and distinct patterns, potentially identifying specific “molecular phenotypes” that may contribute to distinct underlying cellular defects. An example of how such correlations can be used is provided in **Figure 3**, in this case examining lipid-protein correlations for protein cluster 8 and the corresponding lipid clusters 4, 7, and 11 (**Figure 3A and 3B**, also see **Figure 2C**). We plotted all lipid-protein pairs with at least one correlation with Tau > 0.4, performed GO enrichment analysis and highlighted autophagy proteins of interest (red dots). Proteins in cluster 8 had overall high correlations to the select lipid species, which was especially true for a subset of autophagy receptors. We next examined the Top10 enriched proteins per lipid species and stratified them by their frequency (**Figure 3C**). Almost half of the Top10 proteins were either ubiquitin-binding autophagy receptors, NCOA4, or ATG8 proteins.

**Figure 3.**
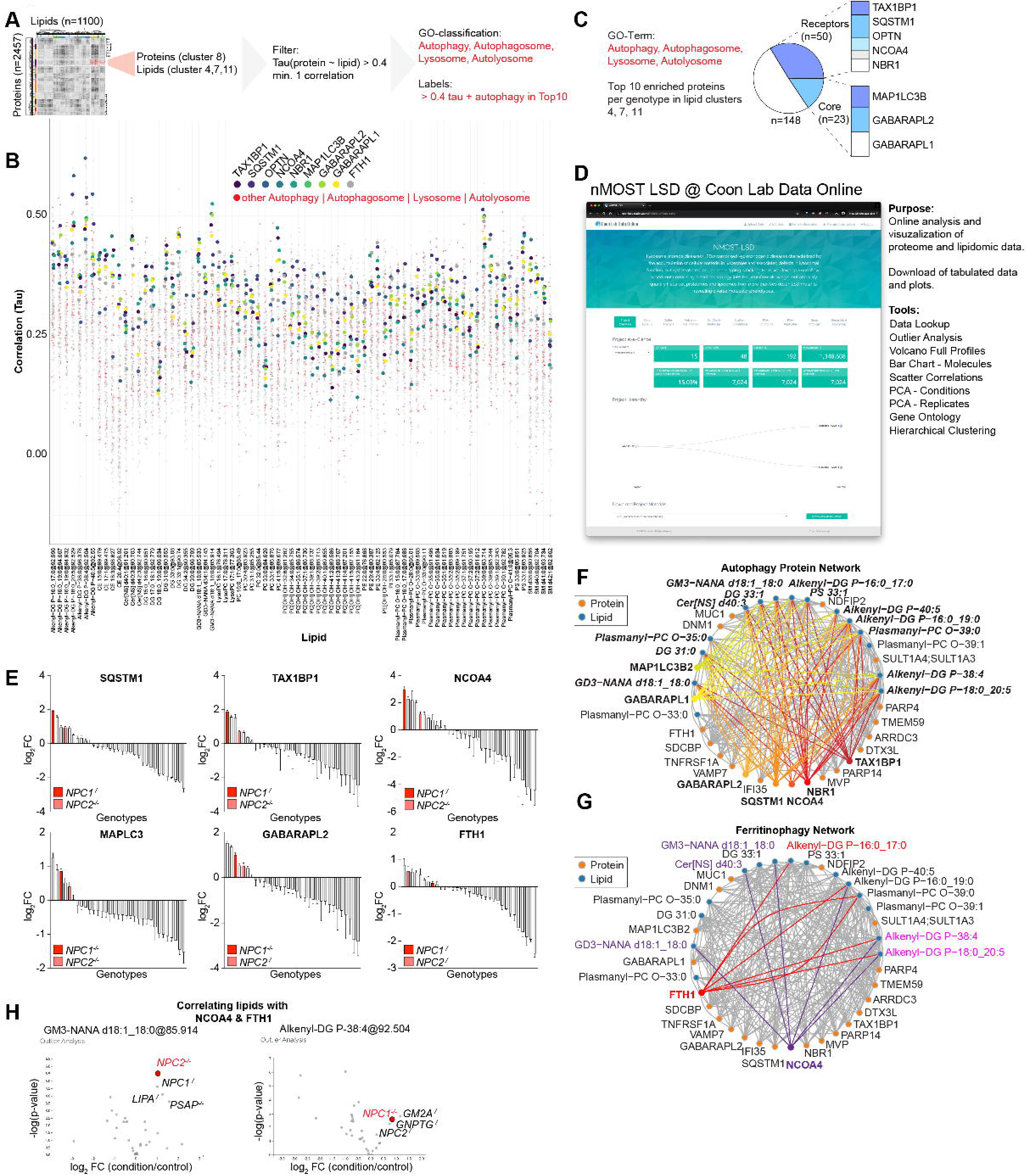
An nMOST-LSD Resource for Lipid-Protein Correlation Analysis. **(A)** Schematic of search strategy to find functional protein-lipid relationships from LSD-nMOST cross-ome dataset. **(B)** Manhattan-style plot of lipid species (x-axis, lipid clusters 4, 7, 11) vs. protein correlations from protein cluster 8. Red dots represent proteins associated with GO-terms autophagy, autophagosome, lysosome, autolysosome. Additionally, select autophagy proteins are highlighted in viridis-color scheme. **(C)** Pie-chart of Top 10 enriched proteins from panel **B**. Autophagy receptors and autophagy core components represent ∼50 of the hits and composition is shown on the right. **(D)** Screenshot of nMOST LSD on the Coon Lab Data Online portal. Tools available online are listed on the right. **(E)** Log_2_FC ranked bargraph of indicated autophagy proteins across all analysed LSD genotypes. *NPC1^-/-^* and *NPC2^-/-^*genotypes are highlighted in shades of red. Data extracted from online portal. **(F,G)** Protein- lipid network extracted from protein cluster 8 and lipid clusters 4,7,11. **F** depicts protein-lipid connections for autophagy markers. **G** depicts protein-lipid connections for ferritinophagy markers. **(H)** Outlier analysis of two lipid species highlight correlated with NCOA4 and FTH1 (see **G**). Data extracted from online portal.

To facilitate further mining of this data by the community, we established an online portal, where the LSD-nMOST data can be parsed, analysed and visualized in multiple ways (**Figure 3D**). This portal allows visualization of abundance profiles for individual proteins across all of the LSD mutant cell line proteomes, as shown for several autophagy receptors, and highlighting the finding cells lacking NPC1 or NPC2 typically exhibit among the highest levels of these receptors, especially the ferritinophagy receptor NCOA4 and one of its main cargo, FTH1 (**Figure 3E**). Network analysis for general autophagy markers (**Figure 3F**) or ferritinophagy (**Figure 3G**) revealed correlations involving specific lipids in distinct lipid classes, including GABARAPL1, and ubiquitin-binding autophagy receptors with alkenyl-DG, GM3, Cer[NS], and DG3 species (**Figure 3F**). Linkage between multiple ubiquitin-binding autophagy receptors and these specific lipids may provide a lipid fingerprint associated with accumulation of specific receptors. Similarly, the portal allows the identification of mutant cell lines that display significantly different patterns of specific biomolecules (on both protein and lipids level) compared to the rest of the LSD-nMOST dataset, and which other genotypes display a similar pattern (Outlier-Analysis). For example, *NPC1*^-/-^ and *NPC2*^-/-^ display strong correlations between the NCOA4/FTH1 pair and alkenyl-DG or GM3 species (**Figure 3F,G**), and several other genotypes showed similar lipid outlier profiles (**Figure 3H**).

### nMOST analysis of lysosomal cholesterol pathway mutants

The data described above highlighted the link between autophagy signatures and LSD proteins linked with the lysosomal cholesterol transport pathway. To examine these relationships in greater detail, we performed validation experiments in cells lacking *NPC1*, *NPC2*, or *LIPA* – as well as GAA, a glycogen storage mutant linked with Pompe’s disease (**Figure S2A, Table S1**). As expected, lysosomes of *NPC1*^-/-^ and *NPC2*^-/-^ mutants, but not *LIPA*^-/-^ or *GAA*^-/-^ mutants, dramatically accumulate cholesterol within lysosomes based on staining with the cholesterol binding probe Filipin (**Figure S2A and S2B**). Lysosomes in *NPC1*^-/-^ and *NPC2*^-/-^ cells displayed slightly elevated pH (5.9 and 6.5, respectively, compared with *LIPA*^-/-^, *GAA*^-/-^, and Control cells, with pH ∼5.4-5.7) in line with previous reports^22^ (**Figure S2C**).

We performed nMOST analysis of this “4KO” cohort in quadruplicate replicates under both full media (Fed) and nutrient stress conditions know to induce autophagy (EBSS, 6 h) (**Figure S2D; Table S4**). The absence of the deletion target was verified by label-free quantification of nMOST data (**Figure S2E**) and PCA analysis revealed high sample and treatment reproducibility (**Figure S2F and S2G**), pointing to the robustness of the nMOST method. Hierarchical clustering of the 1007 lipids quantified revealed distinct alterations in the abundance of multiple lipid classes, with *NPC1*^-/-^ and *NPC2*^-/-^ cells clustering together, as anticipated (**Figure S2H**). Interestingly, individual lipid species within specific classes demonstrates divergent patters of accumulation or loss for individual genotypes (e.g. PC species in *LIPA*^-/-^ or *NPC1*^-/-^ cells) (**Figure S2H**). As explored below, we identified a set of Lyso- PC species that were selectively increased in *NPC1*^-/-^ and *NPC2*^-/-^ cells in both Fed and starved conditions (**Figure S2H**). Similarly, hierarchical clustering of proteomic data focused on the major cellular organelle systems with links to autophagy and mitochondrial pathways (see **STAR METHODS** for description of curated organelle proteomes) revealed distinct patterns of alterations in protein abundance based on genotype, with *NPC1*^-/-^ and *NPC2*^-/-^ cells clustering together with *LIPA*^-/-^ (**Figure S2I**). By comparison, *GAA*^-/-^ cells had fewer alterations in protein abundance relative to control cells (**Figure S2I**).

The patterns of protein alterations confirmed the major findings of the large-scale LSD screen, and identified alterations in the abundance of several mitochondrial functional categories, as well as lysosomal and autophagy proteins in *LIPA^-/-^*, *NPC1*^-/-^, and *NPC2*^-/-^ mutants, but not *GAA*^-/-^ cells (**Figure S2I**). In particular, we confirmed an increase in the abundance of Ub-binding receptors SQSTM1, TAX1BP1, and NBR1, as well as LC3B (MAP1LC3B), under both Fed and EBSS-treated conditions, particularly in *NPC1*^-/-^ and *NPC2*^-/-^ cells (**Figure 4A**), a result that recapitulates previous studies examining lysosomes in *NPC1*^-/-^ cells.^14^ Importantly, this phenotype appeared to be a direct reflection of loss of NPC pathway function, as selective inhibition of the NPC1 transporter with the small molecule U18666A in Control HeLa^TMEM192-HA^ cells resulted in analogous accumulation of SQSTM1, LC3B, and GABARAP proteins, as determined by immunoblotting of cell extracts (**Figure 4B**).

**Figure 4.**
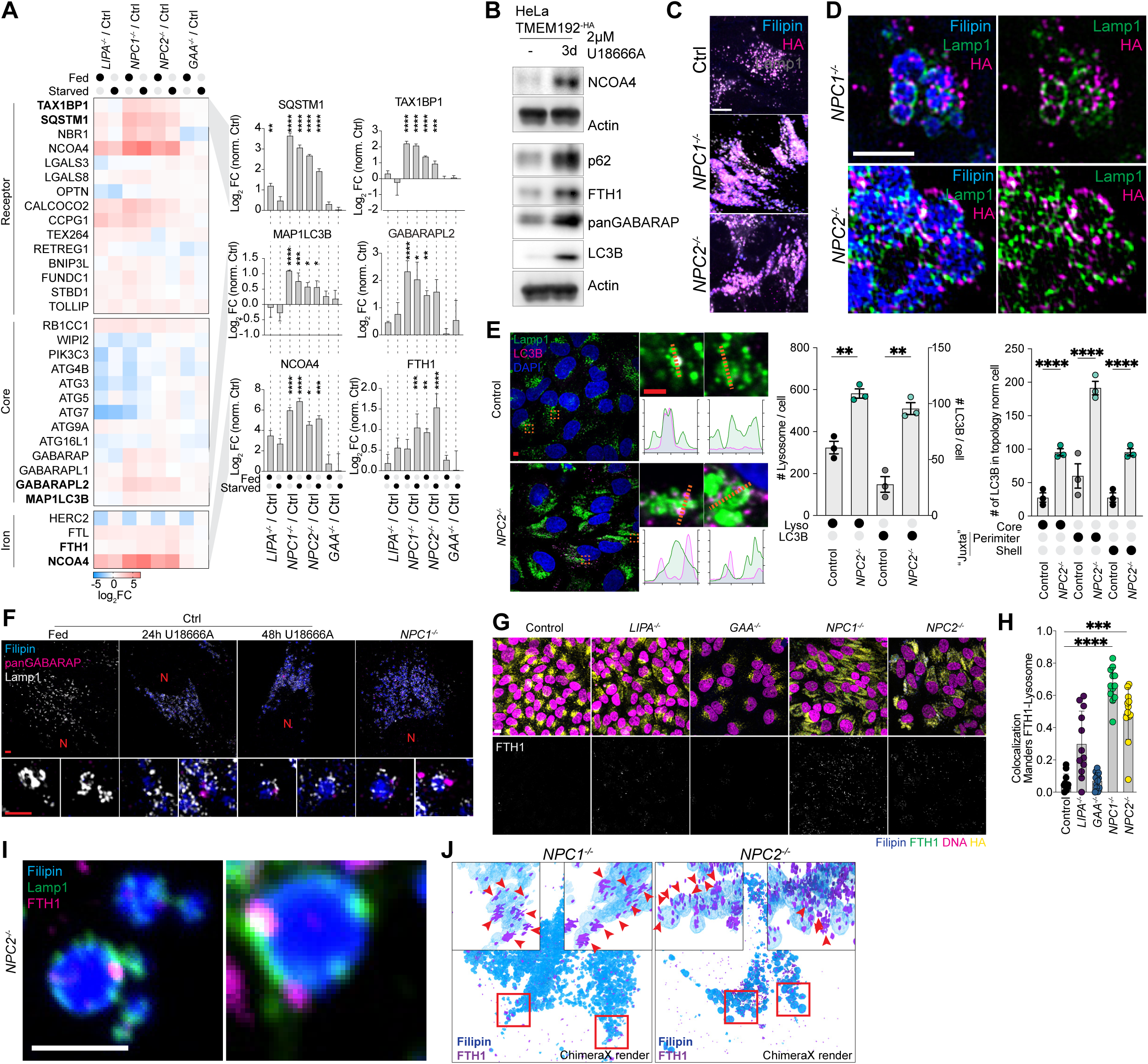
Juxta-lysosomal accumulation of autophagy receptors and ferritin in *NPC1*^-/-^ and *NPC2*^-/-^ cells. **(A)** Log_2_FC relative to Control cells for the indicated autophagy receptors for 4KO cells. MAPLC3B: p(****) <0.0001, p(***) = 0.0001, p(*)=0.0129 & 0.0157. SQSTM1: p(****) <0.0001, p(**) = 0.0047. TAX1BP1: p(****) <0.0001, p(***) = 0.0001. NBR1: p(****) <0.0001. NCOA4: p(****) <0.0001, p(*) = 0.0244; FTH1: p(****) <0.0001; p(***) = 0.0007, p(**) = 0.0075. Data based on quadruplicate replicate nMOST measurements, ordinary one-way ANOVA with multiple comparisons, alpha = 0.05. Error bars depict S.D. **(B)** Western Blot for select autophagy and ferritinophagy proteins of whole cell lysates from HeLa Control treated for 3 days w/o U18666A. **(C)** The indicated cells were stained with Filipin to stain cholesterol-rich lysosomes and immunostained with α-LAMP1 and α-HA to detect TMEM192^-HA^ in lysosomes, followed by imaging with confocal microscopy. Scale bars = 10 µm. **(D)** Cells from panel C were imaged using 3D-SIM. Scale bar = 2 µm. **(E)** Immunostaining of Control and *NPC2*^-/-^ cells with α-LAMP1, α-LC3B and nuclei stained with DAPI. Line trace plots across individual Lamp1- positive lysosomes and the corresponding LC3B intensities. Scale bars = 5 µm and 2 µm (insets). Quantification of LC3B localization relative to LAMP1 is plotted on the right. **(F)** 3D-SIM reconstructions of HeLa Control cells treated for the indicated times with U18666A and *NPC1^-/-^* immunostained for α-LAMP1 and α-panGABARAP, with lysosomes marked by Filipin staining. **(G)** Confocal fluorescent images of control and 4KO cells in Fed cells immunostained with α- FTH1 and α-HA to detect TMEM192^HA^. Cholesterol-rich lysosomes were stained with Filipin and nuclei were stained with DNA SPY555. Scale bar = 20 µm. **(H)** Images from panel G were quantified by measuring Mander’s overlap between FTH1 signal and the lysosome mask provided by α-HA staining. Data from 12 image stack per condition; genotype (Number of cells Fed): Ctrl(3307), *LIPA*^-/-^(1996), *GAA*^-/-^(1401), *NPC1*^-/-^(1211), *NPC2*^-/-^(1629). Error bars depict S.D. **(I)** Confocal images of *NPC2*^-/-^ cells immunostained for α-LAMP1 and α-FTH1, with lysosomes marked by Filipin staining. A single z-slice is shown. Scale bar = 2 µm. **(J)** 3D-SIM reconstructions of *NPC1*^-/-^ or *NPC2*^-/-^ cells immunostained with α-FTH1 and the surface volume of cholesterol-rich lysosomes marked by Filipin.

### Juxta-lysosomal autophagy receptors and cargo in NPC mutants

Lysosomes of *NPC1*^-/-^ and *NPC2*^-/-^ HeLa^TMEM192-HA^ cells exhibited a swollen morphology, with Filipin-positive lumen and puncta corresponding to LAMP1 and TMEM192^HA^, known to localize in the limiting lysosomal membrane, typically decorating the outer layers of the Filipin positive core (**Figure 4C and 4D; S3A**). Previous work^14^ concluded that LC3B accumulated within the lysosomal lumen in *NPC1*^-/-^ cells, and proposed defects in lysosomal degradation as being responsible for receptor accumulation. To examine this possibility further, we measured the abundance and localization of LC3B in Control and *NPC* mutants. Loss of either NPC protein resulted in significantly increased LC3B and SQSTM1 puncta (**Figure S3B and S3C**). However, in *NPC2^-/-^*cells, the majority of LC3B puncta were detected at juxta-lysosomal locations, as opposed to the core region of lysosomes (**Figure 4E**). We did not observe significant accumulation of LC3B in Control cells, as expected (**Figure 4E and Figure S3D**). This ATG8-accumulation phenotype was replicated by treatment of Control cells with increasing durations of U18666A treatment, resulting in the accumulation of analogous juxta-lysosomal GABARAP puncta around aberrant swollen lysosomes (**Figure 4F**), analogous to observations in *NPC1*^-/-^ cells.

In addition to general autophagy receptors, *NPC1*^-/-^ and *NPC2*^-/-^ cells accumulated NCOA4, a receptor for ferritinophagy,^18,19^ and the cargo proteins FTH1 and FTL (**Figure 4A**). Ferritin, a cage-like protein complex composed FTH1 and/or FTL proteins, binds ∼4500 Fe^+3^ atoms and also promotes the conversion of Fe^+2^ species to Fe^+3^ to reduce reactivity.^23^ FTH1 complexes are delivered to lysosomes as well as late endosomes via NCOA4, which directly binds a conserved motif in FTH1 and promotes encapsulation within an autophagosome.^18,24–26^ As with general autophagy receptors, NCOA4 and FTH1 accumulated in response to NPC1 inhibition with U18666A (**Figure 4B**), indicating a direct effect of NPC pathway inhibition. Consistent with a defect in ferritinophagy, FTH1-positive puncta significantly accumulated in *NPC1^-/-^*and *NPC2^-/-^* (**Figure 4G and 4H**), and at closer inspection, such puncta were found in the proximity of lysosomes (**Figure 4I**). The localization of juxta-lysosomal FHT1 puncta was verified using three-dimensional structured illumination microscopy (3D-SIM), with the majority of FTH1 signal external to or only partially embedded in Filipin-positive structures (**Figure 4J**).

Previous work has established a pathway for <u>C</u>onjugation of <u>A</u>TG8 proteins to <u>S</u>ingle <u>M</u>embranes (CASM) such as lysosomes in response to various signals through an ATG16L1- dependent mechanism.^27^ As such, we examined whether increased juxta-lysosomal ATG8 proteins in *NPC* mutant cells reflected CASM. Although Control cells display increased GABARAP-II species when treated with the CASM activator MLSA5,^28^ an agonist of the lysosomal TRPML1 cation channel,^29^ levels of GABARAP-II were already very high in *NPC1*^-/-^ cells and were not further increased with MLSA5 (**Figure S3E**). Moreover, levels of GABARAP-II remained elevated in the presence of VPS34 inhibitor, which blocks canonical autophagy (**Figure S3E**), suggesting that elevated GABARAP-II and LC3B-II levels may not be reversible within the time frame of this experiment. Therefore, as an orthogonal approach to examine a possible role for CASM in ATG8 accumulation, we used SopF, a *Samonella* effector protein known to inhibit CASM by blocking interaction between ATG16L1 and V-ATPase.^30,31^ We blocked NPC1 activity in Control cells that were transiently expressing GFP-SopF using U18666A (24 h) and examined ATG8 proteins and lysosomal co-localization by imaging in cells with or without GFP-SopF (**Figure S3F**). SopF expression led to a ∼50% reduction in the fraction of LAMP1-positive lysosomes with co-incident GABARAP, but this effect was independent of NPC1 inhibition (**Figure S3F**). Total LC3B puncta and increased lysosomal size verified NPC1 inhibition in this experiment (**Figure S3F**). Thus, we conclude that autophagy receptors such as ATG8 proteins accumulate in a juxta-lysosomal localization, with no obvious role for CASM under the conditions tested. Together, these data indicate an inability of *NPC1* and *NPC2* mutants to successfully deliver multiple types of autophagic receptors and FTH1 cargo to the lysosomal lumen is the driving force behind the autophagic deficiency, as opposed to a defect in the process of lysosomal degradation, *per se*.^14^

### Defective endocytic cargo delivery to lysosomes in *NPC2*^-/-^ cells

The finding that autophagic receptors and FTH1 cargo accumulate at juxta-lysosomal locations in *NPC1*^-/-^ and *NPC2*^-/-^ cells suggested an inability of lysosomes to efficiently fuse with autophagosomes. Endocytosis represents a distinct pathway for lysosomal trafficking, where late endosomes fuse with lysosomes to facilitate degradation of endocytic cargo. We hypothesized that if lysosomes from *NPC2*^-/-^ cells are defective in this process, cargo containing vesicles would accumulate in juxta-lysosomal locations analogous to that observed with autophagy receptors (**Figure 4A**). To examine this hypothesis, Control and *NPC2*^-/-^ cells under Fed conditions were incubated with the extracellular endocytic cargo Dextran647 and LysoTrackerRed (to visualize lysosomes). Live-cell 3D-SIM revealed that while Dextran647 puncta in Control cells were largely co-incident with the lysosomal lumen, the Dextran647 signal in *NPC2*^-/-^ cells was largely excluded, and appeared to form partial “halo”-type structures surrounding lysosomes (**Figure 5B**). The simplest explanation for these results is that attempted fusion of Dextran647-loaded endosomes with cholesterol-laden lysosomes places Dextran647 signal co-incident with the limiting membrane without allowing full access to the lysosomal lumen. As with LC3, SQSTM1, and FTH1, much of the Dextran647 signal in either cells grown on Glucose or Galactose remained juxta-lysosomal (**Figure 5B and 5C**), consistent with defective fusion. Importantly, juxta-lysosomal Dextran647 signal was also observed in Control cells treated for 3 days with U18666A to an extent similar to that observed in *NPC1*^-/-^ cells (**Figure 5D**), indicating that defects in cholesterol efflux from lysosomes can rapidly lead to defects in lysosomal function independent of prolonged efflux defects in the context of cells constitutively lacking *NPC1*.

**Figure 5:**
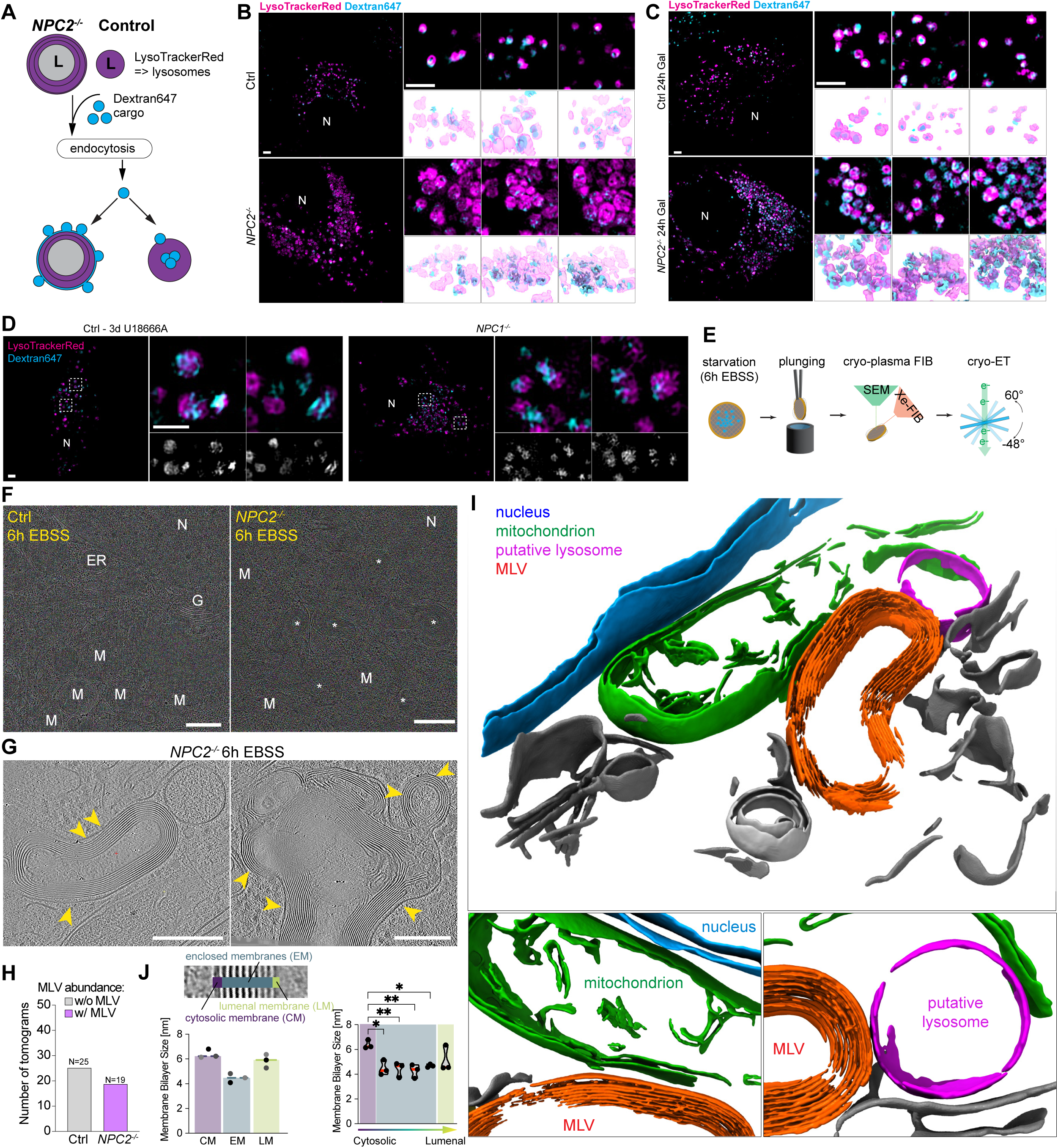
Visualization of multi-lamellar membranes in *NPC2*^-/-^ lysosomes by cryo-ET. **(A)** Schematic showing endocytosis of dextran and its ultimate incorporation into the lysosome in Control and *NPC2*^-/-^ cells. In Control cells, dextran endocytosis successfully delivers dextran to the lysosomal lumen via vesicle fusion. In *NPC2*^-/-^ cells with multi-lamellar membranes, successful fusion and delivery of dextran is reduced and successful fusion events result in dextran present in the limited lumenal space between the limiting lysosomal membrane and the first internal membrane. **(B)** Control or *NPC2*^-/-^ cells were treated with dextran conjugated with Alexa647 dye and imaged by live-cell 3D-SIM. Renderings derived from 3D-SIM reconstructions are shown below. Scale bar = 2 µm. **(C)** As in panel B, but cells were cultured on Galactose growth media for 24h. Scale bar = 2 µm. **(D)** Control cells were treated for 3 days with the NPC1 inhibitor U18666A incubated alongside *NPC1^-/-^* cells with dextran conjugated with Alexa647 dye and imaged by live-cell 3D-SIM. Renderings derived from 3D-SIM reconstructions are shown below. Scale bar = 2 µm. **(E)** Schematic of the plasma-FIB and cryo- ET workflow. **(F)** Example lamella overviews of Control and *NPC2*^-/-^ cells under 6 h EBSS nutrient starvation conditions. Scale bar = 500 nm. **(G)** Example tomogram slice of multi- lamellar vesicles in *NPC2*^-/-^ cells. Scale bar = 200 nm. **(H)** Quantification of MLV-containing tomograms from Control and *NPC2*^-/-^ cells. Total number of tomograms analyzed is stated above the bar charts. (**I**) 3D-renderings of a segmented *NPC2*^-/-^ tomogram. Zoom-ins highlighting close proximity between MLV (orange) with mitochondria (green) and a putative lysosome (pink) are shown beneath. **(J)** Quantification of membrane bilayer size (left) and distance between membrane leaflets (right) across three tomograms for the cytosolic membrane (CM), the enclosed membranes (EM), and the luminal membrane (LM). Quantification of the spacing between individual membranes: CM to first EM (left), between EMs (middle), and EM to LM (right). p(*) = 0.011, 0.21; p(**) = 0.0086, 0.0052. Data based on triplicate experiments (lamellae), ordinary one-way ANOVA with multiple comparisons, alpha = 0.05. Error bars depict S.D. Abbreviations: ER = Endoplasmic reticulum; MLV = Multi-lamellar vesicle.

As an addition measure of lysosomal function, we examined whether *NPC1*^-/-^ cells maintained the ability to release cations upon TRPML1 activation. We loaded Hela Control and *NPC1^-/-^* cells with the cell impermeable calcium indicator Oregon Green 488 BAPTA-5N (OGB- 5N) and measured its lysosomal signal with or without MLSA5 treatment (**Figure S3G**). Control cells displayed a reduction in lysosomal calcium indicator, consistent with cation efflux upon TRPML1 activation. In contrast, lysosomes in *NPC1*^-/-^ cells retained calcium indicator, suggesting that efflux was inhibited in the absence of NPC1 (**Figure S3G**). These data indicate that multiple aspects of lysosomal function are compromised in cells lacking NPC pathway function.

### Multi-lamellar vesicles in *NPC2*^-/-^ cells correlate with Lyso-PC abundance by nMOST

The observations described above led us to examine lysosomal ultrastructure at higher resolution than is possible by light microscopy. Consistent with previous studies^32^, electron microscopy (EM) revealed numerous enlarged vesicular structures in *NPC1*^-/-^ and *NPC2*^-/-^ cells, containing densely stained membrane structures which were rare in Control cells (**Figure S3H**). To examine the morphology of these structures *in situ* without fixation artefacts, we made use of semi-automated cryo-plasma FIB (cryo-PFIB) milling paired with cryo-ET (electron tomography). (**Figure 5E, Figure S4A**). Consistent with classical EM, *NPC2^-/-^* cells treated with EBSS harboured numerous multi-lamellar vesicles (MLVs). These structures were frequently seen in low-magnification EM-overviews but rarely in Control cells (**Figure 5F, white stars**). Based on the size and cellular localization of these MLVs, we hypothesized that these structures correspond to the aberrant, cholesterol-filled lysosomes characterized by fluorescent microscopy (**Figure 5C, 5D, 5F and Figure S2B**). Analysis of the corresponding tomograms (25 for Control, 19 for *NPC2^-/-^* cells) revealed that the majority of MLVs in *NPC2*^-/-^ cells were structurally aberrant, containing as many as 17 highly organized membranes surrounding a lumen (**Figure 5F-H**). Segmentation of tomograms highlight the highly organized membrane structures with *NPC2*^-/-^ MLVs (**Figure 5I**), as well as their close proximity to mitochondria. Interestingly, the intermembrane distance inside MLVs was highly regular (2.6 ± 0.2 nm between membranes at half maximum), whereas the distance of the outer limiting membrane to the first enclosed membrane was highly variable across MLVs (**Figure 5G** yellow arrows, **Figure S4B-D**). For the limiting membrane, both leaflets were clearly distinguishable with full width at half maximum (FWHM) of 6.4 ± 0.4 nm. (**Figure 5J**). However, the interleaflet space of the enclosed membranes and the luminal membrane could not be resolved at the magnification used and showed an FWHM of 4.5 ± 0.4 nm and 5.8 ± 0.5 nm, respectively. (**Figure 5J; S4B- D**).

Accumulation of MLVs suggested potential alterations in lipids abundance, possibly as a consequence of defective cholesterol efflux. We therefore employed nMOST data to globally examine lipid alterations in *NPC* mutant cells with or without starvation, identifying alterations in two major lipid classes. First, cholesterol ester (CE) abundance was generally elevated in whole-cell lipidomics of *NPC2^-/-^* cells, as indicated by a skew of CE species towards the upper end of ranked abundance relative to Control cells (**Figure S2H and S4E**). This was particularly evident for CE species with chain lengths shorter than 20 carbons (**Figure S4E**). Retrieval of lipid-protein cross-ome correlation networks of select high-abundance CE species in *NPC2^-/-^* cells from the LSD-nMOST dataset revealed correlation with autophagy and ferritinophagy markers (**Figure S4F**). Second, we observed strong accumulation of Lyso-PC species, major building blocks of cellular membranes, but not Lyso-PE species and other types of phospholipids, in *NPC2*^-/-^ cells under both fed and starvation conditions (**Figure S4G**). LysoPC species likewise carry autophagy/ferritinophagy signatures (**Figure S4H**). Third, increased LysoPC appeared selective for *NPC* mutants, as LysoPC levels in *LIPA^-/-^* or *GAA^-/-^* cells were less strongly affected (**Figure S4I**). Interestingly, Lyso-PC species that were enriched in *NPC2^-/-^* cells had, on average, chain lengths of <20 carbons (**Figure S4J**). We speculate that characteristic and tightly packed spacing of MLVs in *NPC2*^-/-^ cells (**Figure 5I**) could reflect the generation of multi-lamellar membranes enriched in shorter chain Lyso-PC species.

### OXPHOS complex and cristae defects in *NPC2*^-/-^ cells

The finding that *NPC1^-/-^*and *NPC2^-/-^* cells exhibited accumulation of FTH1 and NCOA4 led us to explore whether iron dependent processes downstream of lysosomal function and ferritinophagy are affected. We analyzed nMOST data from *NPC1*^-/-^, *NPC2*^-/-^, *LIPA*^-/-^, *GAA*^-/-^, and Control cells in either Fed or EBSS-treated (6 h) states for alterations in two systems that are heavily reliant on iron availability – cytosolic Fe-S cluster assembly machinery^33^ and components of the mitochondrial OXPHOS system, which contain several subunits with Fe-S clusters^34^. We were particularly interested in the OXPHOS system as GO terms related to this were found enriched in *NPC2*^-/-^ cells by nMOST (**Figure S1J and S2I**).^35^ The inner-membrane space (IMS) compartment in *NPC2^-/-^* cells was especially depleted of organelle-annotated proteins (**Figure 6A, Table S4**). When normalized to Control, we observed a reduction in a cohort of Complex I (CI) subunits in *NPC2*^-/-^ cells under starvation conditions, which was most pronounced for components of the N-module (log_2_FC ∼-0.41) (**Figure 6A-D and Figure S5A and S5B**). Five of the 8 subunits within the N-module of CI contain Fe-S clusters, consistent with a reliance on iron for stability and/or assembly.^34,36^ The abundance of the Q-module, which contains 4 subunits with Fe-S clusters, as well as Complex IV (CIV), was also slightly reduced in *NPC2*^-/-^ mutants in the presence of EBSS (log_2_FC ∼-0.1) (**Figure 6A-D and Figure S5A and S5B**). These alterations were in contrast with the abundance of mitochondrial Fe-S cluster assembly machinery in *NPC1*^-/-^ and *NPC2*^-/-^ cells, which was largely unchanged or slightly increased when compared with Control cells (**Figure S5C and S5D**).

**Figure 6:**
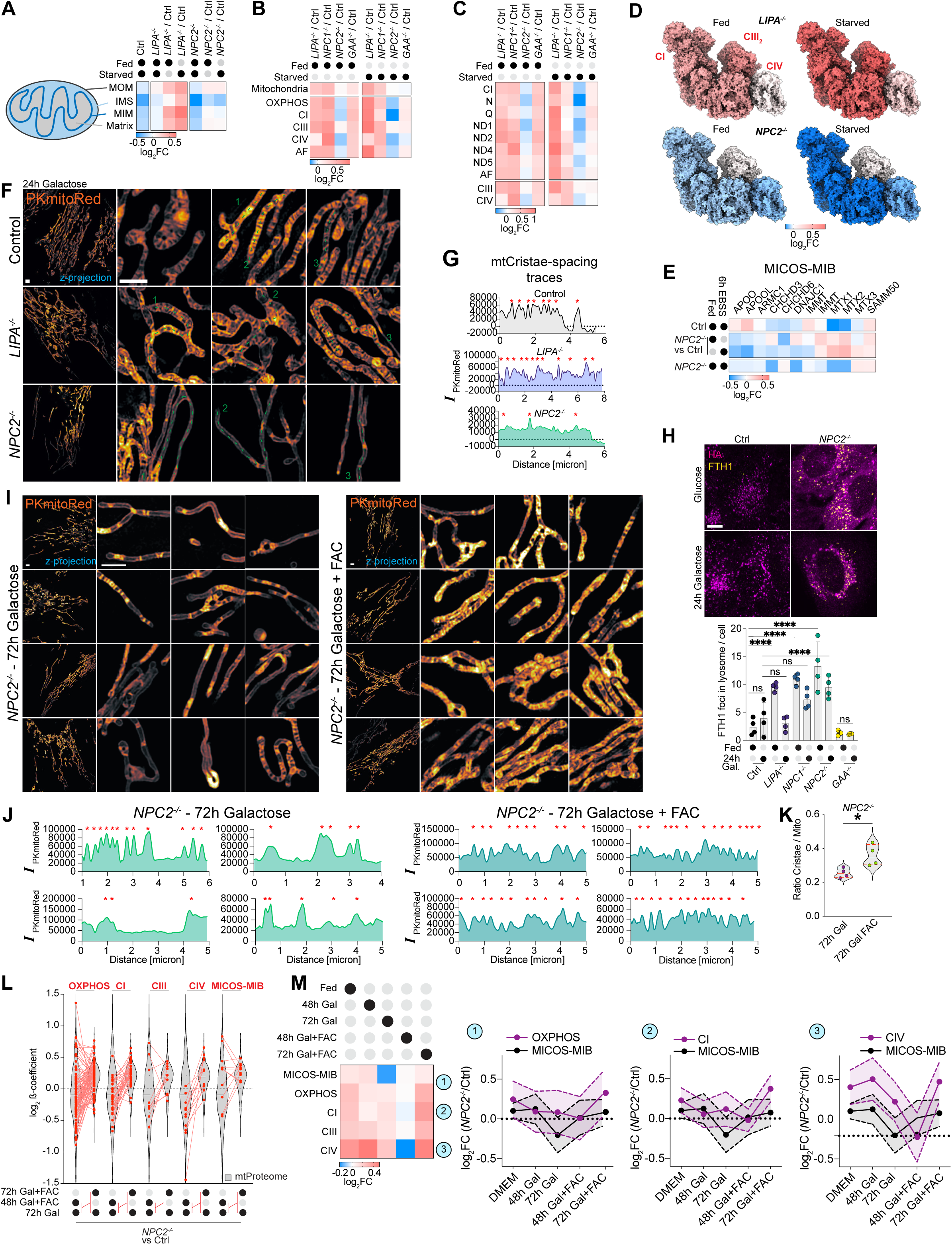
Defects in mitochondrial cristae/OXPHOS systems in *NPC2*^-/-^ cells and amelioration by extracellular iron. **(A)** Heatmap depicting log_2_FC of components of different mitochondrial compartments in either Fed and EBSS-treated Control and 4KO cells. Data based on quadruplicate biological replicate nMOST measurements. **(B,C)** Panel B: Log_2_FC values for modules within OXPHOS for 4KO cells in Fed and EBSS-treated cells based on quadruplicate replicate nMOST data. Panel C: Log_2_FC of replisome sub-module abundance comparing Fed versus EBSS treated Control, *LIPA*^-/-^, *GAA*^-/-^, *NPC1*^-/-^, and *NPC2*^-/-^ cells based on quadruplicate biological replicate nMOST data. **(D)** Abundance of heatmaps in **B** mapped onto structure of mitochondrial respirasome (CI, CIII_2_, CIV from PDB: 5XTH). Legend shows colour panel for log_2_FC values. Based on quadruplicate replicate nMOST data. **(E)** Heatmap for components of the MICOS-MIB complex in for 4KO cells [normalized with Control]. Data based on quadruplicate biological replicate nMOST measurements. **(F)** Z-projections of live-cell 3D-SIM images from Control, *LIPA^-/-^* and *NPC2^-/-^* cells after culturing on galactose (24 h) stained with the IMS dye PKmitoRed. Scale bar = 2 µm. **(G)** Line-plots of dashed lines from panel **F** of Control, *LIPA^-/-^* and *NPC2^-/-^* cells on 24 h galactose growth conditions. **(H)** Confocal images of Control and *NPC2*^-/-^ cells grown in glucose (Fed) or Galactose (24h) followed by immunostaining with α-HA to detect TMEM192^HA^ and α- FTH1 to detect Ferritin. Scale bar = 10 µm. Right panel displayed quantification of FTH1 signal overlapping with lysosome staining/cell. Data from biological quadruplicates per sample (each replicate containing 5-9 stacks); p(****) <0.0001, ordinary two-way ANOVA with multiple comparisons, alpha = 0.05. Error bars depict S.D.. **(I)** Z-projections of live-cell 3D-SIM images from Control and *NPC2*^-/-^ cells after culturing on Galactose (72 h) with or without FAC and stained with the IMS dye PKmitoRed. Scale bar = 2 µm. **(J)** Line-plots of individual mitochondria from panel **I**. Red asterisks indicate positions of cristae. **(K)** Violin plot depicting the ratio of cristae to mitochondria with and without FAC addition. Data based on 132 (72 h Gal) or 148 (72 h Gal + FAC) segmented planes of ROI-stacks from data shown in panel **I**; p(*) = 0.0242, unpaired t.test. **(L)** Log_2_ β-coefficient for the indicated treatments shown for all OXPHOS subunits and individual sub-complexes. **(M)** Log_2_FC [*NPC2*^-/-^/Control] for the indicate protein complexes under the indicated conditions (time in Galactose with or without FAC addback). Data based on triplicate biological replicate TMTpro measurements.

The intimate connectivity between OXPHOS complex assembly and mitochondrial cristae structure^37^, together with the finding that several MICOS-MIB complex components are reduced in *NPC2*^-/-^ cells relative to Control cells (particularly in EBSS conditions, **Figure 6E**), led us to examine mitochondrial ultrastructure. Utilizing a photo-stable and cell-permeable IMS dye PKmitoRed, combined with live-cell 3D-SIM, we observed alterations in cristae morphology particularly in *NPC2*^-/-^ cells in the presence of galactose to enforce OXPHOS utilization (**Figure 6F and 6G**). Unlike Control cells, which displayed regularly spaced cristae, *NPC2*^-/-^ cells displayed an unexpected morphology reflective of altered cristae structure, including extensive regions of mitochondria, often near the cell periphery, that lacked obvious cristae as indicated by lineplots (**Figure 6F and 6G**; red stars indicate cristae bridge). In contrast with *NPC2*^-/-^ cells, *LIPA*^-/-^ cells displayed an increased number of cristae, albeit with less regular intervals (**Figure 6F, G**), in line with the overall increase in OXPHOS and mitochondrial proteome compared to Control cells (**Figure 6B and 6C, and Figure S5B**). These data indicate multiple defects in mitochondrial morphology and proteome abundance in the absence of *NPC2*.

### Alleviation of cristae morphology defects in *NPC2*^-/-^ cells by extracellular iron

The apparent defect in ferritinophagy in *NPC2*^-/-^ cells led us to examine whether mitochondrial defects could be mechanistically linked with iron availability. As an alternative to iron mobilization by ferritinophagy, we tested whether extracellular iron delivered to the cytoplasm via Transferrin-dependent endocytosis could rescue cristae morphology. Transferrin-associated iron undergoes endocytosis, where the reduced pH of the endosome allows iron release from Transferrin and transport to the cytosol via the DMT1/SLC11A2 proton-coupled metal ion transporter.^38^

We first verified that FTH1 abundance, juxta-lysosomal FTH1 localization, and endolysosomal system fusion phenotypes are retained in *NPC2*^-/-^ cells grown in Galactose (**Figure 6H**). We then examined cristae morphology in *NPC2*^-/-^ cells grown in Galactose in the presence or absence of FAC (ferric ammonium citrate, 72 h) as an extracellular iron source. Both the frequency of cristae and their spacing were significantly rescued by FAC addition (**Figure 6I-K**). Change of growth medium slightly reduced the average lysosomal size and percentage of Filipin-positive lysosomes, but FAC addition did not further influence these parameters (**Figure S5E and S5F**), indicating that FAC does not rescue lysosomal defects associated with *NPC* deficiency. We examined mitochondrial membrane potential (ΔΨm) in *NPC2*^-/-^ and Control cells by determining the ratio of Tetramethylrhodamine Methyl Ester (TMRE) to MitotrackerDeepRed (mtDR, as mask). Control cells grown on galactose (72 h) displayed high membrane potential, as indicated by the prominent TMRM signal (**Figure S5H**). In contrast, *NPC2*^-/-^ cells displayed reduced ΔΨm, a phenotype which was partially ameliorated with the addition of FAC (**Figure S5H**). Consistent with the partial rescue of cristae morphology and ΔΨm after FAC addback, *NPC2*^-/-^ cell growth of in Galactose media conditions was increase by ∼ 28% (**Figure S5I**). Taken together, these data indicate that these aspects of mitochondrial dysfunction in *NPC2*^-/-^ cells are partially alleviated by providing a ferritinophagy- independent route for iron delivery to the cytosol and mitochondria.

### MICOS-MIB complex proteome remodelling in *NPC2*^-/-^ cells by extracellular iron

To examine the effect of FAC addition on the mitochondrial proteome in an unbiased manner, we performed two 18-plex Tandem Mass Tagging (TMT) proteomics experiments examining total proteomes from Control or *NPC2^-/-^*cells under 5 conditions: DMEM, 48 h Galactose ± FAC, 72 h Galactose ± FAC (**Figure S5J, Table S5**). Mirroring the observations from the 4KO-nMOST dataset, we observed an overall increase in the abundance of autophagy receptors in *NPC2*^-/-^ (including LC3B, SQSTM1, TAX1BP1, NCOA4, FTH1 and FTL) however, other proteins related to iron homeostasis and FeS cluster assembly were largely unaltered (**Figure S5K-M**).

We next examined the mitochondrial proteome. *NPC2*^-/-^ cells cultured for 72 h in Galactose with FAC displayed a slight increase in the mitochondrial proteome compared to Galactose conditions alone; however, focusing on proteins associated with mitochondrial OXPHOS complexes revealed differential changes in response to Galactose and FAC addback (**Figure S5N and S5O**). Mapping of protein-group abundance on mitochondrial sub-compartments highlighted a spatio-temporal element in mitochondrial proteome remodelling during growth media switch and iron add-back (**Figure S5P**). Indeed, *NPC2*^-/-^ cells grown in Galactose (72 h) displayed a reduction in the abundance of MICOS-MIB complex components when compared with Control cells (**Figure S5Q and S5R**), in accordance with the observed reduction in cristae (**Figure 6F, 6G, and 6I**). In contrast, the abundance of MICOS-MIB complex subunits was largely rescued by FAC for either 48 or 72 h, consistent with imaging data described above (**Figure S5Q and S5R**). Alterations in the abundance of individual MICOS subunits are displayed schematically in **Figure S5S**. Taken together, these results indicate that imbalances in iron homeostasis can lead to reversible changes in mitochondrial ultrastructure.

### OXPHOS complex proteome remodelling in *NPC2*^-/-^ cells by extracellular iron

We next systematically examined the effect of FAC on electron transport chain components in *NPC2*^-/-^ cells. The average abundance of OXPHOS subunits as a cohort increasing over the FAC addback time-course (**Figure S5O**). Focusing on CI, changes in FeS cluster-containing N- and Q-modules revealed the largest abundance shifts in the presence of FAC. Interestingly, at early time-points these modules appear de-stabilized, followed by stabilization of the membrane-arm modules (ND1,2,4,5) and a subsequent increase in N-module proteins between 48 and 72 h of FAC addback (**Figure S5T**). Distinct patterns were observed for individual respirasome (CI-CIII_2_-CIV) modules and assembly factors, as shown for *NPC2*^-/-^ versus Control cells (**Figure S6A and S6B**). A linear increase in CI abundance was observed upon FAC addback when comparing [*NPC2*^-/-^/Ctrl], which was co-incident with increased levels of respective assembly factors. However, CIV displayed a bi-phasic pattern with an initial decrease in abundance at the 48 h time-point (**Figure S6A and S6B**).

Next, we evaluated the effect the growth media conditions and FAC addback has on the proteome in the context of *NPC2*^-/-^ vs Control (β-coefficient, **Figure 6L and Figure S6C**). Log_2_ β-coefficients for the transition from 48 to 72 h in the presence of FAC and Galactose were not significant for the proteome at large (β-coefficient = 0.008); however, although individual subunits displayed differential alterations in abundance, generally detectable increases in mitochondria (β-coefficient = 0.16), and especially OXPHOS components (β-coefficient = 0.21) were observed. This included all but one subunit of N- and Q-modules, as well as all the nuclear genome-encoded CIV subunits (**Figure S6C**). Consistent with the differences observed on the abundance between 48 and 72 h, β-coefficients across OXPHOS components at these time-points revealed differential changes that were not see with MICOS-MIB subunits (**Figure 6L**). Interestingly, recovery of CI and CIV subunits primarily occurred during the 48 to

72 h interval and was preceded temporally by MICOS-MIB rescue (**Figure 6L**). We also examined alterations the abundance of proteins known to function in assembly of Fe-S clusters in either the cytoplasm or the mitochondria (**Figure S5M**). The log_2_FC *NPC2*^-/-^/Control values of most cytosolic Fe-S cluster assembly (CIA) components^39^ were slightly decreased in response to 72 h of FAC, while, in contrast, the abundance of mitochondrial Fe-S cluster (ISC) assembly proteins were either increased or remained constant, with the exception of ABCB7, which functions to transport [2Fe-2S]-(Glutathione)_4_ from the mitochondria to the cytosol.^39^ These data are consistent with an elevated Fe-S cluster biogenesis pathway in mitochondria of *NPC2*^-/-^ cells treated with FAC when compared with Control cells. The effect of FAC addback on the abundance of OXPHOS components is summarized schematically in **Figure S6D**.

### Effect of FAC on OXPHOS abundance during *in vitro* neurogenesis

Mouse models for *NPC*-deficiency displayed disturbances in iron-homeostasis^40^, potentially impacting iron availability in brain tissue. We set out to investigate if the ferritinophagy defect observed in HeLa cells would likewise affect mitochondrial OXPHOS complexes in neuronal models. Stem cells undergo a metabolic switch from glycolysis to OXPHOS during early stages of neuronal differentiation *in vitro*.^41^ We therefore asked whether *NPC* deficiency and/or FAC addback in this system would affect electron transport chain abundance during NGN2-driven *in vitro* neurogenesis. Control, *NPC1*^-/-^, and *NPC2*^-/-^ stem cells containing an inducible AAVS1-NGN2 cassette (see **STAR METHODS**) were differentiated over a 3-week period with or without FAC and subjected to proteomic and phenotypic analysis (**Figure 7A**). As expected, *NPC* mutants displayed an increase in the number of LAMP1- positive lysosomes at day 14 of differentiation (**Figure S7A**), consistent with results in HeLa cells. For proteomic analysis, cells from triplicate cultures at 5 time points during differentiation were subjected to label-free nDIA proteomics^42^ (**Figure 7A, Table S6**). Despite the number and complexity of the samples, the number of proteins quantified remained stable across runs and randomly inserted stem cell quality control (QC) lysates (n=22) demonstrated high reproducibility, with samples distinguished by PCA analysis (**Figure 7B, Figure S7B and S7C**). Neither loss of *NPC1^-/-^* or *NPC2^-/-^*, confirmed by proteomics (**Figure S7D**), or FAC addition, altered the known timing of alterations in pluripotency and differentiation markers (**Figure S7E**),^41^ and as expected, FAC addition promoted the accumulation of FTH1 by immunoblotting and proteomics (**Figure S7F and S7G**).

**Figure 7:**
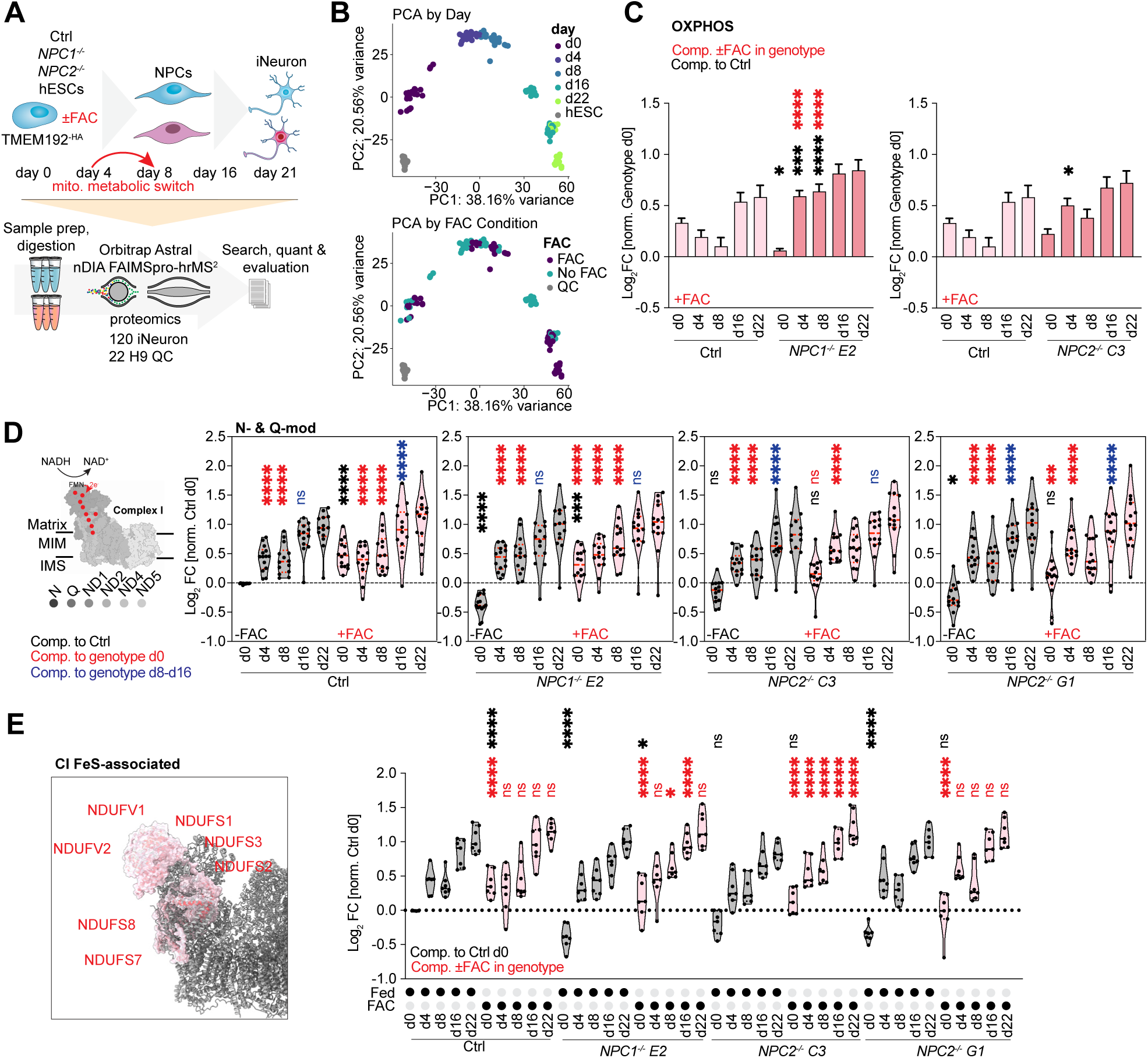
Proteomic analysis of NPC mutants during neurogenesis with iron addback. **(A)** Schematic of experimental approach to study effect of iron supplementation during NGN_2_- driven neurogenesis and downstream label-free proteomic analysis. For each time-point samples were analysed in triplicates. **(B)** PCA plot of LFQ data, color-coded according to day of differentiation or ±FAC. **(C)** Bargraph of mean log_2_FC (normalized within genotype day 0) mitochondrial OXPHOS-components in presence of FAC. Black asterisks indicate statistical comparison to Control cells; red asterisks indicate comparison ± FAC within genotype. Data from triplicate replicates; *NPC1^-/-^* E2 d0: p(*) = 0.0286; d4: p(***) = 0.0003; d8, d4+FAC, d8+FAC: p(****) <0.0001. *NPC2^-/-^* C3 d0: p(*) = 0.0420. Ordinary two-way ANOVA with multiple comparisons, alpha = 0.05. Error bars depict S.E.M.. **(D)** Violin plots of log2FC (normalized to Control at day 0) N- and Q-module of Complex I. A schematic of Complex I and FeS clusters is shown in the left (based on PDB: 5XTH). Abundance comparisons over a three-week differentiation time-course ± FAC treatment are shown for each genotype (grey for -FAC, red for +FAC). Black asterisks indicate statistical comparison to Control cells; red asterisks indicate comparison within genotype. Blue asterisks indicate comparison of day 8 to 16 within genotype. Data from triplicate replicates; Control d0,4,8: p(****) <0.0001; d4,8+FAC: p(****) <0.0001; d8- 16: (****) <0.0001. *NPC1^-/-^* E2 d0: (****) <0.0001; d0+FAC: p(***) = 0.0002; all other: p(****) <0.0001. *NPC2^-/-^* C3: p(****) <0.0001. *NPC2^-/-^* G1 d0: p(*) = 0.0396; d0+FAC: p(**) = 0.0041; all other: p(****) <0.0001. Ordinary two-way ANOVA with multiple comparisons, alpha = 0.05. Error bars depict S.E.M.. **(E)** Surface renderings of FeS-associated Complex I N- and Q-modules (red) overlayed with rest of the protein complex rendered in grey (PDB: 5XTH). Violinplots of log_2_FC (normalized to Control at day 0) FeS-associated proteins. Abundance comparisons over a three-week differentiation time-course ± FAC treatment are shown for each genotype (grey for -FAC, red for +FAC). Black asterisks indicate statistical comparison to Control cells; red asterisks indicate comparison within genotype ±FAC. Data from triplicate replicates; Control: p(****) <0.0001. *NPC1^-/-^* E2 d0: (*) = 0.0489; d8+FAC: p(*) = 0.0146; all other: p(****) <0.0001. *NPC2^-/-^* C3: p(****) <0.0001. *NPC2^-/-^* G1: p(****) <0.0001. Ordinary two-way ANOVA with multiple comparisons, alpha = 0.05. Error bars depict S.E.M..

Analysis of the abundance of OXPHOS components during the differentiation time course revealed three core findings: First, in stem cells, the global abundance of OXPHOS components was reduced in *NPC1^-/-^* and *NPC2^-/-^* mutants, when normalized with Control cells, and this was rescued in the presence of FAC (**Figure S7H**). Thus, stem cells recapitulate the central finding in HeLa cells. Second, during differentiation, total OXPHOS abundance increases across all genotypes, consistent with conversion from glycolysis to oxidative phosphorylation, but FAC had a more profound effect on *NPC1^-/-^*and *NPC2^-/-^* mutants especially on day 4 or day 8 than in Control cells (**Figure 7C and Figure S7I**). Third, the effect of FAC addition on the abundance of N and Q modules as a whole, as well as iron-sulfur cluster containing components within these modules, was also evident in stem cells (**Figure 7D and 7E**). Taken together, these data indicate features of the effect on *NPC1^-/-^* or *NPC2^-/-^* deficiency and iron regulation are present in the stem cell *in vitro* neurogenesis paradigm.

## DISCUSSION

Here we report the nMOST workflow for simultaneous analysis of lipids and proteins from the same sample and its application to a collection of more than two dozen cell lines lacking individual LSD genes. Cross-correlation analysis between lipids and proteins across various genotypes reveals numerous molecular fingerprints associated with specific LSD alleles, providing a resource for further mechanistic discovery. We performed accompanying proteomics experiments using DDA-TMTpro and nDIA-LFQ to validate and follow up the biological insights we gained from nMOST, and to our knowledge, this work represents one of the most in-depth attempts in profiling LSDs, spanning multiple cell types and growth conditions.

We observed a prominent and selective phenotype with NPC1, NPC2 and TPP1 mutants involving accumulation of autophagy regulators, which correlated with accumulation of LysoPC. Through 3D-SIM imaging, we provided evidence for a block in autophagic clearance wherein autophagic receptors (e.g. LC3B) or cargo (e.g. FTH1) accumulate in juxta-lysosomal locations with evidence of defective delivery of cargo to the lysosomal lumen. This phenotype in *NPC2*^-/-^ cells correlated with the formation of multilamellar lysosomes, which we speculate may be reflective of the increased abundance of LysoPC in these cells (**Figure 2C**). Previous studies^15,43^ have implicated decreased lysosomal cleavage of cargo as well as defects in autophagosome-lysosome fusion, but the underlying mechanisms were unclear. We suggest that multilamellar membranes within lysosomes observed by cryo-ET reduce the ability of lysosomes to efficiently fuse with either autophagosomes or endosomes, thereby limiting delivery of cargo to the lysosomal lumen.

Among the autophagic cargo that accumulated in *NPC1*^-/-^ and *NPC2*^-/-^ cells was the ferritin cage protein FTH1, which was juxta-lysosomal based on SR imaging. Given that a block to ferritin degradation in the lysosome would be expected to reduce iron availability, we examined complexes known to rely on Fe-S clusters for their production, leading to the identification of mitochondrial electron transport chain complexes as being reduced in cells lacking *NPC2* (**Figure S6D**, left panel). Loss of OXPHOS complexes correlated with reduced cristae number and MICOS-MIB complexes in *NPC2*^-/-^ cells (**Figure S6D**, left panel). Importantly, delivery of iron to cells through endocytosis results in initial accumulation of MICOS-MIB subunits at 48 h (**Figure S6D**, middle panel), which supports further assembly of OXPHOS complexes at 72 h, with near full restoration of OXPHOS complexes and cristae number (**Figure S6D**, right panel). The behaviour of MICOS-MIB and OXPHOS complexes and the effects on cristae number are consistent with the self-reinforcing role that these components play in formation and stabilization of cristae.^37^ We note that while cells lacking *NPC1* also accumulate FTH1, the corresponding phenotypes and in particular the mitochondrial alterations seen in *NPC2*^-/-^ cells appear more pronounced, despite only 5% of NPC patients carry mutations in the *NPC2* gene.^44^ This phenotype held also true in stem-cell derived iNeurons. Previous studies have described an imbalance of iron metabolism and haematological abnormalities in *NPC1* mouse models and in patients with Niemann-Pick disease type C1.^40^ Further studies are required to understand the extent to which an inability to promote iron mobilization by autophagy and concomitant effects on mitochondrial function are linked with defects observed in patients. We note that while disruption of iron homeostasis in budding yeast has been linked with mitochondrial defects, the underlying mechanisms appear to be distinct.^45^ The discovery platform we have described here and its application to relevant cell lineages linked with LSDs may facilitate identification of molecular defects or pathways with relevance to disease.

## LIMITATIONS OF THE STUDY

First, our studies do not address how loss of *NPC1* or *NPC2* leads to accumulation of plasmalogen and lyso-PC lipid species, although one possibility is that these alterations are a result of defects in cholesterol distribution in various membranes. Second, our collection of HeLa cells lacking LSD genes is incomplete, and further studies are required to obtain and characterize the full set LSD mutants. In addition, while cancer cell lines such as HeLa are known to display phenotypes in common with more physiologically relevant cell system, including for example loss of *GRN* and its effect on lipids,^4^ analysis of a broader array of cell types is required to understand the generality of lipidomic and proteomic phenotypes reporter here. Our analysis of ES cell derived iNeurons revealed related phenotypes particularly in the ES cell state, but these cell types also do not fully represent many facets of cell-cell communication and interactions that exist under physiological conditions. Finally, the basis for the distinct mitochondrial phenotypes for *NPC1^-/-^*and *NPC2^-/-^* mutants in HeLa cells is unknown, although the apparent bias for accumulation of FTH1 in *NPC2^-/-^* when compared with *NPC1^-/-^*cells could be relevant. In this regard, previous studies have suggested that NPC2 can deliver cholesterol directly to the endosomal membrane, which may have a thinner glycocalyx, possibly allowing cholesterol to reach other cellular membranes in the absence of NPC1.^46,47^ As such, cells lacking NPC2 may generate a stronger mitochondrial phenotype, a topic that requires further study.

## Supporting information

Key Resources Table

## ACKNOWLEDGEMENTS

This study used the infrastructure of the Department of Cell and Virus Structure at the MPI of Biochemistry. We thank the Core for Imaging Technology & Education and the Cell Biology Microscopy Facility at Harvard Medical School for microscopy support. We thank Dr. Simon Nørrelykke and the Image Analysis Collaboratory for discussions about quantitative image analysis. Electron Microscopy Imaging, consultation and /or services were performed in the HMS Electron Microscopy Facility. This work was supported by Warren Alpert Foundation (to J.W.H.), the Bluefield Project (to J.W.H.), Aligning Science Across Parkinson’s (ASAP, to J.W.H., B.A.S.), NIH (R01NS083524, R01NS110395 to J.W.H., P41GM108538 to J.J.C., R01GM132129 to J.A.P.), the Max Planck Society (B.A.S., F.W.). F.K. was supported by a fellowship from the Alfred and Joan Goldberg Education and Fellowship Fund (Department of Cell Biology, Harvard Medical School). Michael J Fox Foundation administers the grants ASAP- 000282/ASAP-024268 on behalf of ASAP and itself. For the purpose of open access, the author has applied a CC-BY public copyright license to the Author Accepted Manuscript (AAM) version arising from this submission.

## AUTHOR CONTRIBUTIONS

Conceptualization: F.K., S.S., Y.H., J.J.C., J.W.H.; nMOST development and lipidomics: Y.H., K.A.O., A.J., N.M.N., B.J.A. under supervision of J.J.C.; Cell line creation/characterization: .S., Y.J., F.K.; Cell biology and light microscopy: F.K.; TMT/nDIA proteomics: F.K., Y.H., J.A.P., S.P.G; Cryo-ET: J.B., F.K., C.C., A.B. under supervision of B.A.S., J.M.P. and F.W.; Lysosomal pH: C.L.; Bioinformatics: F.K., I.R.S., Y.H, K.A.O. The manuscript was written by J.W.H. and F.K. with input from all authors.

## STAR METHODS

All details and catalogue numbers can be found in the **Key Resource Table**. Protocols can be found on protocols.io (dx.doi.org/10.17504/protocols.io.5qpvokmmzl4o/v1).

### Cell culture

HeLa TMEM192-3xHA cells (referred to as HeLa^TMEM192-HA^)^20^ were maintained in Dulbecco’s modified Eagle’s medium (DMEM), supplemented with 10% vol/vol fetal bovine serum (FBS), 5% vol/vol penicillin-streptomycin (P/S), 5% vol/vol GlutaMAX and 5% vol/vol non-essential amino acids (NEAA) at 37°C, 5% O_2_. Unless otherwise noted, we refer to independently grown and handled cultures as biological replicates to distinguish from assays performed on identical samples (i.e. technical replicates). For galactose growth conditions, galactose-containing DMEM was prepared from Glucose-free DMEM supplemented with 10% vol/vol dialysed FBS, 25 mM D-galactose, 5% vol/vol penicillin-streptomycin (P/S), 5% vol/vol GlutaMAX), 5% vol/vol Sodium Pyruvate and 50 µg/mL Uridine.

### Gene-Editing

Generation of LSD mutants in the HeLa^TMEM192-HA^ or H9 AAVS-NGN2^TMEM192-HA^ background^20,48^ was facilitated using CRISPR/Cas9 with target sites determined using CHOPCHOP.^49^ Guide RNAs were ligated into the px459 plasmid (Addgene plasmid # 62988) and cells transfected using Lipofectaime LTX reagent (Thermo Fisher Scientific, 15338100), according to manufacturer’s instructions. Two days post-transfection, single, puromycin-resistant cells were sorted into 96-well dishes containing 300 µL full growth medium. To generate *NPC1^-/-^* or *NPC2^- /-^* in H9 ESCs, 0.6 μg sgRNA was incubated with 3 μg SpCas9 protein for 10 minutes at room temperature and electroporated into 2x10^5^ WT H9 cells using Neon transfection system (Thermo Fisher Scientific) and sorted into 96-well dishes containing 300 µL full growth medium (composition as described above). Single cells were allowed to grow into colonies and duplicated for multiplex sequencing. Genomic DNA samples were obtained by incubating cells in 30 µL PBND (50 mM KCl, 10 mM Tris-HCl, pH 8.3, 2.5 mM MgCl_2_-6H_2_O, 0.45% NP-40 and 0.45% Tween-20) with protease K (40 µg/ml) at 37°C for 5 min and heated to 55°C and 95°C for 30min and 15 min, respectively. The first round of PCR was performed to amplify the target region using gene-specific primers that contain partial Illumina adaptor sequences (i.e., Forward primer: 5’- ACACTCTTTCCCTACACGACGCTCTTCCGATCT[n]_18-22_ -3’, Reverse primer: 5’- GTGACTGGAGTTCAGACGTGTGCTCTTCCGATCT[n]_18-22_ -3’, [n]_18-22_ represent gene specific sequences). The resulting PCR products with adapter-modified ends can be further amplified in the second round of PCR by universal primers containing attachment sites for the flow cell and index sequences (i.e., Forward primer: 5’- AATGATACGGCGACCACCGAGATCTACACTCTTTCCCTACACGACGCTCTTC CGATCT -3’, Reverse primer: 5’- CAAGCAGAAGACGGCATACGAGAT-[n]_8_-GTGACTGGAGTTCAG ACGTGTGCT -3’, [n]_8_ represents index sequences). The final PCR products were purified using QIAquick PCR purification kit (Qiagen, 28106). Sequencing was performed using Miseq Reagent kits v2 on Illumina Miseq following the denature and dilute libraries guide of Miseq system, and sequencing data was analysed by Outknocker program (www.OutKnocker.org).^50^ Knockout candidates were confirmed by Western blot on whole cell lysates or by proteomics. The sgRNAs were generated using GeneArt Precision gRNA Synthesis Kit (Thermo Fisher Scientific) according to the manufacturer’s instruction and purified using RNeasy Mini Kit (Qiagen). The sgRNA target sequences and sequencing results can be found in **Table S1**. HeLa^TMEM192-HA^ Control, *GRN*^-/-^ and *HEXA*^-/-^ as well as the H9 AAVS-NGN2^TMEM192-HA^ Control cells have been previously reported.^8,20,48^

### iNeuron differentiation

Human ES cells (H9, WiCell Institute) were cultured in E8 medium ^51,41^ on Geltrex-coated tissue culture plates with daily medium change. Cells were passaged every 4-5 days with 0.5 mM EDTA in 1× DPBS (Thermo Fisher Scientific). SpCas9 and AsCas12a/AsCpf1 expression plasmids pET-NLS-Cas9-6xHis (Addgene plasmid # 62934) and modified pDEST-his-AsCpf1- EC, generated by deleting the MBP sequence from plasmid pDEST-hisMBP-AsCpf1-EC (Addgene plasmid # 79007), were transformed into *Rosetta™(DE3)pLysS* Competent Cells (Novagen), respectively, for expression. SpCas9 and AsCas12a/AsCpf1 proteins were purified as described elsewhere.^52,53^ Briefly, cells expressing SpCas9 [0.5 mM isopropylthio-β-galactoside, 14 h induction] were lysed in FastBreak buffer (Promega, Inc) and the NaCl concentration adjusted to 500 mM. Extracts were centrifuged at 38,000 g for 10 min at 4°C and the supernatant incubated with Ni-NTA resin for 1 h. The resin was washed extensively with 50 mM Tris pH8.0, 500 mM NaCl, 10% glycerol, 20 mM imidazole, and 2 mM TCEP prior to elution with this buffer supplemented with 400 mM imidazole. Proteins were diluted two volumes/volume in PBS and fractionated on a Heparin-Sepharose column using a 0.1 to 1.0 M NaCl gradient. Cas9-containing fractions were stored in PBS, 20% glycerol, 2 mM TCEP at - 80°C. AsCpf1 expression was induced similarly, and cells pelleted by centrifugation. Cells were lysed by sonication in 50 mM HEPES pH7, 200 mM NaCl, 5 mM MgCl_2_, 1 mM DTT, and 10 mM imidazole supplemented with lysozyme (1 mg/ml) and protease inhibitors (Roche complete, EDTA-free). After centrifugation (16,000 g for 30 min), the supernatant was incubated with Ni-NTA resin, the resin washed with 2M NaCl, and bound proteins eluted with 250 mM imidazole, and buffer exchanged into lysis buffer lacking MgCl_2_ and imidazole prior to storage at -80°C.

For human ES cell conversion to iNeurons, cells were expanded and plated at 2x10^4^/cm^2^ on Geltrex-coated tissue plates in DMEM/F12 supplemented with 1x N2, 1x NEAA (Thermo Fisher Scientific), human Brain-derived neurotrophic factor (BDNF, 10 ng/ml, PeproTech), human Neurotrophin-3 (NT-3, 10 ng/ml, PeproTech), mouse laminin (0.2 μg/ml, Cultrex), Y-27632 (10 μM, PeproTech) and Doxycycline (2 μg/ml, Alfa Aesar) on Day 0. On Day 1, Y-27632 was withdrawn. On Day 2, medium was replaced with Neurobasal medium supplemented with 1x B27 and 1x Glutamax (Thermo Fisher Scientific) containing BDNF, NT-3 and 1 μg/ml Doxycycline. Starting on Day 4, half of the medium was replaced every other day thereafter. On Day 7, the cells were treated with Accutase (Thermo Fisher Scientific) and plated at 3-4x10^4^/cm^2^ on Geltrex-coated tissue plates. Doxycycline was withdrawn on Day 10. Following day 12 of differentiation 50% ND2 media was changed every other day. FAC was added if necessary.

### NPC1 inhibitor treatments

Indicated cells were differentiated (if necessary), seeded in 6-well plates and treated with the NPC1 inhibitor U18666A for 1 – 3 days at the indicated concentrations in regular growth medium at 37°C. Before harvesting, cells were washed twice with 1xPBS and harvested for further experimental procedures.

### MLSA5 and VPS34 inhibitor treatments

Indicated cells were differentiated (if necessary), seeded in 6-well plates. Cells were treated for 2 h with the VPS34 inhibitor SAR405 (1 µM) or 3 h with MLSA5 (10 µM) in regular growth medium at 37°C. A double treated sample was also included. Before harvesting, cells were washed twice with 1xPBS and harvested for further experimental procedures.

### Western Blotting

At the indicated times, ES cells, iNeurons or HeLa cells were washed on ice in 1x PBS, harvested and pellet wash with 1x PBS and resuspended in 8 M urea buffer (8 M urea, 150 mM TRIS pH 7.4, 50 mM NaCl, PhosSTOP phosphatase inhibitor cocktail). Resuspended cell lysates were sonicated for 10 seconds and debris pelleted at 13,000 rpm for 10 min. Protein concentration was determined by BCA assay according to manufacturer’s instructions (Thermo Fisher Scientific, 23227). Indicated amounts of proteins were resuspended in 1xLDS + 100 mM DTT and boiled for 10 minutes at 85°C. Equal amount of protein and volume were loaded run on 4%-20% Bis-Tris, 8% Tris NuPAGE gels for 5 minutes at 100 V, 5 min at 150 V and then run at 200 V for the required time. Gels were transferred via wet transfer system onto PDVF membranes for Western Blotting. Chemiluminescence and colorimetric images were acquired using a BioRad ChemiDoc MP imaging system. Images from Western Blots were exported and analysed using Image Lab and ImageJ/FiJi.^54^

### PROTEOMICS

#### Sample Preparation for nMOST

For samples used for technical evaluation of MOST, Bead-enabled, Accelerated, Monophasic Multi-omics (BAMM) method was used.^12^ Silica coated superparamagnetic beads (700 nm, SeraSil-Mag) were washed and resuspended in water for a concentration of 75 μg/μL, while frozen cell pellets were being thawed on ice. 200 μL acetonitrile (ACN), 600 μL n-butanol, and 200 μL beads containing water were added to samples. After vortex, samples were sonicated for 5 minutes at 14°C. Beads were immobilized by magnet and 100 μL supernatant was aliquoted, dried down, and reconstituted in 300 μL n-butanol:isoproponal (IPA):water (8:23:69, v/v/v) in an amber autosampler vial for lipids.^55,56^ The remaining supernatant was removed. The beads were reconstituted in Rapid Digestion Buffer (Promega) diluted to 75% by water with 2 mM TCEP and 40 mM CAA. After incubation for 10 minutes at room temperature, trypsin (Promega) was added in a 20:1 ratio (protein-to-trypsin). The samples were incubated in thermomixer for 40 minutes at 60°C and 1000 RPM. Formic acid was added to terminate digestion. Peptides were desalted by Sep-Pak (Waters) C18 column, dried down in SpeedVac (Thermo Fisher Scientific), and reconstituted in 0.2% formic acid.

For HeLa whole cell extracts or Lyso-IP samples (generated as described^8,20^; dx.doi.org/10.17504/protocols.io.ewov14pjyvr2/v2.) samples, 300 μL mixture of n-butanol:ACN:water (3:1:1, v/v/v) was added. Samples were bath sonicated for 5 min at 14°C. After centrifugation at 14000 g for 5 minutes, 50 μL of the lipid containing supernatant was transferred to autosampler vials with glass insert, dried down in SpeedVac, and resuspend in 50 μL n-butanol:IPA:water (8:23:69, v/v/v).^55,56^ The remaining samples were maintained at - 80°C until protein digestion. For protein digestion of whole cell extracts, the samples were thawed on ice and centrifuged at 14000 g for 5 minutes. The remaining supernatant was removed from samples. 100 μL lysis buffer (8 M Urea, 100 mM Tris pH 8.0, 10 mM TCEP, 40 mM CAA) was added. The samples were bath sonicated for 5 minutes at 14°C and vortexed for 15min. Protein concentration was determined by Thermo protein BCA assay (reducing agent compatible). LysC (FUJIFILM Wako) was added to samples in a 50:1 ratio (protein-to-LysC) and incubate on a rocker for 4 h at room temperature. The urea was diluted to 2 M by 300 μL 100 mM Tris pH 8.0. Trypsin was added to samples in a 50:1 ratio (protein-to-trypsin) and incubate on a rocker overnight at room temperature. For protein digestion of Lyso-IP samples, 60 μL 6 M GnHCl, 100 mM Tris was added to the sample to solubilize proteins from being aggregated on beads. The samples were bath sonicated 5 minutes at 14°C; incubated in thermomixer for 5 minutes at 100 °C and 600 RPM, and then incubated for 2 h at 80°C and 600 RPM. Beads were immobilized by magnet and supernatant was transferred to a 96-well plate. GnHCl was diluted to 2 M by adding 120 μL 100 mM Tris, 10 mM TCEP, 40 mM CAA. LysC was added to samples in a 50:1 ratio (protein-to-LysC) and incubate on a rocker for 4h at room temperature. GnHCl was diluted to 0.4 M by adding 420 μL 100 mM Tris pH8.0. Trypsin was added to samples in a 50:1 ratio (protein-to-trypsin) and incubate on a rocker overnight at room temperature. 10% TFA was added to terminate digestion. After centrifugation at 12000 g for 5 minutes, digested peptides were desalted by StrataX 10 mg 96-well Plate (Phenomenex), dried down in SpeedVac, and reconstituted in 0.2% formic acid. The peptide concentration was determined by Thermo peptide BCA assay.

#### nMOST LC-MS

Separation was performed on an in-house packed BEH C18 capillary column (28 cm length × 75 μm inner diameter × 1.7 μm particle size) at 60 °C and an Ultimate3000 system (Thermo Scientific). Column packing was described previously.^57^ Mobile phase A consisted of 0.2% formic acid in water. Mobile phase B consisted of 0.2% formic acid and 5 mM ammonium formate in IPA/ACN (90:10, v/v). Lipids were loaded onto column first and then peptides at 0% mobile phase B. Mobile phase B increased to 70% over 80 minutes for scanning MS/MS spectra of peptides, increased to 100% over 26 minutes for scanning MS/MS spectra of lipids. The column was washed at 100% mobile phase B for 3 minutes and re-equilibrated at 0% mobile phase B for 10 minutes. Eluting analytes were analyzed by an Orbitrap Eclipse Tribrid mass spectrometer (Thermo Scientific). Spray voltage was 2 kV. Ion transfer tube temperature was 275°C. MS^1^ scan range was 200-1,600 *m/z*. MS^1^ resolution was 240,000 (at 200 *m/z*). Source RF was 35. MS^1^ AGC target was 300%. MS^1^ injection time was 50 ms. Duty cycle was 1 second. Polarity was positive. For proteomics data acquisition from 0 to 80 minutes, precursor selection range was 300-1,350 *m/z*. Charge states were 2-5. Dynamic exclusion was 10 seconds. Isolation width was 0.5 *m/z*. Precursors were fragmented by higher-energy collisional dissociation (HCD) with a normalized collision energy (NCE) of 25%. MS^2^ mass spectra were acquired in data-dependent mode using ion trap turbo speed. MS^2^ scan range was 150-1,350 *m/z*. MS^2^ AGC target was 300%. MS^2^ injection time was 14 ms. For lipidomics data acquisition from 80 to 120 min, precursor selection range was 300-1,600 *m/z*. Charge states were 1-2. Dynamic exclusion was 10 seconds. Isolation width was 0.7 *m/z*. Precursors were fragmented by higher-energy collisional dissociation (HCD) with a stepped NCE of 27% ± 5%. MS^2^ mass spectra were acquired in data-dependent mode using ion trap rapid speed. MS^2^ scan range was auto. MS^2^ AGC target was 300%. MS^2^ injection time was 17 ms. Real-time library search (RTLS) and complementary collision-induced dissociation (CID) were used for glycerophospholipids and sphingomyelins as described previously.^13^ For large scale LSD samples, to improve the throughput of the analysis, the total LC time was set down to 105 minutes. Peptides eluted and were analyzed from 0 to 70 minutes while lipids eluted and were analyzed from 70 to 105 minutes.

#### nMOST MS Data Process

For proteomics, raw data files were processed by MaxQuant (Version 2.0.3.0). The database was canonical plus isoforms downloaded from Uniprot in December 2021. The match between runs was on. MS/MS spectra were not required for LFQ comparisons. For lipidomics, raw data files were processed using Compound Discoverer 3.1 (Thermo Scientific) and Lipidex.^58^ Peak detection required a signal-to-noise ratio of 1.5, a minimum peak intensity of 5 × 10^5^, and a maximum peak width of 0.75 minutes. The chromatographic peaks were grouped into compound groups by a retention time tolerance of 0.5 minutes and a mass tolerance of 10 ppm. Peaks were removed if the peak areas of sample over blank were < 3-fold. An in silico generated lipid spectral library (LipiDex_HCD_Formic) was used for MS/MS spectra searching.

The threshold of dot product score was 500 and the threshold of reverse dot product score was 700. MS^2^ spectra were annotated at the molecular species level if the minimum spectral purity was at least 75%; otherwise, sum compositions were reported. The lipid identification was further filtered for adducts, dimers, in-source fragments, misidentified isotopes, and mismatched retention time by LipiDex and the degreaser module of LipiDex 2 (https://github.com/coongroup/LipiDex-2).^59^ Cross-ome correlation analysis between lipids and proteins analysed by nMOST. Proteins and lipids were correlated using Kendall rank correlation approach (R function corr(); the resulting matrix was filtered for lipids or proteins with at least 2 correlations |>0.4| Tau. The filtered matrix was further clustered using hierarchical clustering and subsetted into 18 protein clusters and 14 lipid cluster (kmeans). Members of each cluster were evaluated for enrichment in GO terms (Cellular components) or lipids class using a fisher’s exact test.

### Neurogenesis with iron supplementation and label-free nDIA proteomics

Ctrl, *NPC1^-/-^* and *NPC2-* were seeded on Geltrex-coated tissue culture plates and differentiated according to the methods stated above. For d0 and d4 timepoints, cells were seeded in triplicates in coated 12-well plates. For all other timepoints, cells were replated at day 4 in triplicates into coated 12-well plates. FAC was added during regular media changes. Cells were dissociated using 0.5 mM EDTA-PBS and washed in PBS, pelleted at 2000g for 5 minutes, the supernatant aspirated and pellet snap frozen and stored at -80°C.

Cell pellets were lysed in 140 μL 8M Urea and 100 mM Tris pH 8.0 by syringe with a 21G needle. Protein concentration was measured by protein BCA assay. 10 mM TCEP and 40 mM chloroacetamide were added for reduction and alkylation. LysC (FUJIFILM Wako) was added to samples in a 50:1 ratio (protein-to-LysC) and the samples were incubated on a rocker for 4 h at room temperature. The urea was diluted to 2 M with 100 mM Tris pH 8.0. Trypsin was added to samples in a 50:1 ratio (protein-to-trypsin) and the samples were incubated on a rocker overnight at room temperature. Digested peptides were desalted using Waters 25 mg Sep-Pak tC18 96-well Plates (186002319). The concentration of desalted peptides was measured by peptide BCA assay.

Separation was performed on a C18 capillary column (30 cm length × 100 μm inner diameter) packed with Accucore C18 resin (2.6 μm, 150 Å, Thermo Fisher Scientific) at 55 °C and an EASY-nLC 1200 system (Thermo Scientific). Flow rate was set to 450 nL/min. Mobile phase A consisted of 0.125% formic acid in ACN/water (5:95, v/v). Mobile phase B consisted of 0.125% formic acid in ACN/water (95:5, v/v). When peptides were loaded on the column, mobile phase B increased from 5% to 23% over 22 minutes, increased to 100% over 2 minutes, and stayed at 100% for 6 minutes. Eluting peptides were analyzed by an Orbitrap Astral mass spectrometer (Thermo Scientific). Spray voltage was 2.2 kV. Ion transfer tube temperature was 290°C. The FAIMS compensation voltages (CV) was -55 V. MS^1^ scan range was 380-980 *m/z*. MS^1^ resolution was 240,000 (at 200 *m/z*). Source RF was 50%. MS^1^ AGC target was 500%.

MS^1^ injection time was 50 ms. Precursors were isolated in the quadrupole by an isolation width of 2 *m/z* and fragmented by higher-energy collisional dissociation (HCD) with a normalized collision energy (NCE) of 27%. MS^2^ mass spectra were acquired in data-independent mode in the Astral analyzer. MS^2^ scan range was 110-2,000 *m/z*. MS^2^ AGC target was 500%. MS^2^ max injection time was 3 ms.

### Astral LFQ nDIA MS Data Process

Raw files were converted to mzML format using MSConvert^60^ and analyzed using FragPipe (v22.0) with DIA-NN (v1.8.2).^61–64^ All settings were kept default.

### TMTpro 18plex proteomics

#### Proteomic sample preparation

Sample preparation of proteomic analysis of whole-cell extract from HeLa control and mutant lysates performed according to previously published studies.^41,65^ Replicate cell cultures were grown and treated independently and are considered biological replicates in the context of TMT experiments. Cells were washed twice with 1xPBS and harvested on ice using a cell scraper in 1xPBS. Cells were pelleted via centrifugation for 5 minutes (5000*g*, 4°C), and washed with 1xPBS before resuspending in lysis buffer (Urea, 150 mM TRIS pH 7.4, 150mM NaCl, protease and phosphatase inhibitors added). After a 10 second sonication, and optional French-pressing through a G25 needle, lysed cells were pelleted and protein concentration of clarified sample determined using BCA kit (Thermo Fisher Scientific, 23227). 100 µg protein extract of each sample were incubated for 30 minutes @ 37°C with 5 mM TCEP for disulfide bond reduction with subsequent alkylation with 25 mM chloroacetamide for 10 minutes at RT with gentle shaking. Methanol-chloroform precipitation of samples was performed as follows: To each sample, 4 parts MeOH was added, vortexed, one part chloroform added, vortexed, and finally 3 parts water added. After vortexing, suspension was centrifugated for 2 minutes at 14000*g* and the aqueous phase around the protein precipitate removed using a loading tip. Peptides were washed twice with MeOH and resuspended in 200 mM EPPS, pH 8, and digested for 2 h with LysC (1:100) at 37°C, followed by Trypsin digestion (1:100) at 37°C overnight with gentle shaking.

#### Tandem mass tag (TMT) labeling

50 µL of digested samples were labeled by adding 10 µL of TMT reagent (stock: 20 mg/ml in acetonitrile, ACN) together with 10 µL acetonitrile (final acetonitrile concentration of approximately 30% (v/v)) for 2 h at room temperature before quenching the reaction with hydroxylamine to a final concentration of 0.5% (v/v) for 15 minutes. The TMTpro-labeled samples were pooled together at a 1:1 ratio, resulting in consistent peptide amount across all channels. Pooled samples were vacuum centrifuged for 1 h at room temperature to remove ACN, followed by reconstitution in 1% FA, samples were desalted using C18 solid-phase extraction (SPE) (200 mg, Sep-Pak, Waters) and vacuum centrifuged until near dryness.

##### Basic pH reverse phase HPLC

Dried peptides were resuspended in 10 mM NH_4_HCO_3_ pH 8.0 and fractionated using basic pH reverse phase HPLC.^66^ Samples were offline fractionated into 96 fractions over a 90 minutes run by using an Agilent LC1260 with an Agilent 300 Extend C18 column (3.5 μm particles, 2.1 mm ID, and 250 mm in length) with mobile phase A containing 5% acetonitrile and 10 mM NH_4_HCO_3_in LC-MS grade H_2_O, and mobile phase B containing 90% acetonitrile and 10 mM NH_4_HCO_3_ in LC-MS grade H_2_O (both pH 8.0). The 96 resulting fractions were then pooled in a non-continuous manner into 24 fractions.^67^ This set of 24 fraction was divided into 2x12 sets (even or odd numbers), acidified by addition of 1% Formic Acid (FA) and vacuum centrifuged until near dryness. One set (12 samples) was desalted via StageTip, dried and reconstituted in 10 µL 5% ACN, 5% FA before LC-MS/MS processing.

##### Mass spectrometry acquisition

For HeLa whole-cell proteomics, data collection was performed on a Orbitrap Fusion Lumos Tribrid mass spectrometer (Thermo Fisher Scientific, San Jose, CA), coupled with a FAIMS Pro device and a Proxeon EASY-nLC1200 liquid chromatography (Thermo Scientific). 10% of resuspended samples were loaded on a 35 cm analytical column (100 mm inner diameter) packed in-house with Accurcore150 resin (150 Å, 2.6 mm, Thermo Fisher Scientific, San Jose, CA) for LC-MS analysis. Peptide separation was performed with a gradient of acetonitrile (ACN, 0.1 % FA) from 3-13% (0-83 minutes) and 13- 28% (83-90 minutes) during a 90 min run. LC-MS/MS was combined with 3 optimized compensation voltages (CV) parameters on the FAIMS Pro Interface to reduce precursor ion interference.^68^ Data-dependent acquisition (DDA) was performed by selecting the most abundant precursors from each CV’s (-40/-60/-80) MS1 scans for MS/MS over a 1.25 second duty cycle. The parameters for MS1 scans in the Orbitrap include a 400-1,600 m/z mass range at 60,000 resolution (at 200 Th) with 4 x 10^5^ automatic gain control (AGC) (100%), and a maximum injection time (max IT) of 50 ms. Most abundant precursors (with 120 s dynamic exclusion +/- 10 ppm) were selected from MS1 scans, isolated using the quadrupole (0.6 Th isolation), fragmented with higher-energy collisional dissociation (HCD, 36% normalized collision energy), and subjected to MS/MS (MS2) in the Orbitrap detector at 50,000 resolution, 5x AGC, 110 – 200 m/z mass range, IT 86 ms and with 120 s dynamic exclusion +/- 10 ppm.

##### Data processing

Raw mass spectra were converted to mzXML, monoisotopic peaks reassigned using Monocle^69^ and searched using Comet^70^ against all canonical isoforms found in the Human reference proteome database (UniProt Swiss-Prot 2019- 01; https://ftp.uniprot.org/pub/databases/uniprot/previous_major_releases/release-2019_01/)) as well as against sequences from commonly found contaminant proteins and reverse sequences of proteins as decoys, for target-decoy competition.^71^ For searches, a 50-ppm precursor ion tolerance and 0.03 Da product ion tolerance for ion trap MS/MS as well as trypsin endopeptidase specificity on C-terminal with 2 max. missed cleavages was set. Static modifications were set for carbamidomethylation of cysteine residues (+57.021 Da) and TMTpro labels on lysine residues and N-termini of peptides (+304.207 Da); variable modification was set for oxidization of methionine residues (+15.995 Da). Peptide-spectrum matches were filtered at 2% false discovery rate (FDR) using linear discriminant analysis (Picked FDR method, based on XCorr, DeltaCn, missed cleavages, peptide length, precursor mass accuracy, fraction of matched product ions, charge state, and number of modifications per peptide (additionally restricting PSM Xcorr >1 and peptide length>6,^72^ and after a 2% protein FDR target filtering^73^ PSM reporter ion intensities were quantified. Quantification was performed using a 0.003-Da window around the theoretical TMT-reporter m/z, and filtered on precursor isolation specificity of > 0.5 in the MS1 isolation window and the output was filtered using summed SNR across all TMT channels > 100. MS statsTMT^74^ was performed on peptides with >200 summed SNR across TMT channels. For each protein, the filtered peptide– spectrum match TMTpro raw intensities were summed and log_2_ normalized to create protein quantification values (weighted average) and normalized to total TMT channel intensity across all quantified PSMs (adjusted to median total TMT intensity for the TMT channels).^75^ Log_2_ normalized summed protein reporter intensities were compared using a Student’s t-test and p-values were corrected for multiple hypotheses using the Benjamini-Hochberg adjustment.^76^ Linear model analysis was performed as described. ^77^ Subcellular and functional annotations were based on previous published list of high confidence annotations (^78^, “high” & “very high” confidence, additional manual entries from^41,48^, AmiGO Pathway online tool and mitochondrial annotation was based on MitoCarta 3.0^79^). Part of heatmaps were created using Morpheus (https://software.broadinstitute.org/morpheus). GO-enrichment was performed with ShinyGO.^80^

### MICROSCOPY

#### Live-cell spinning disk microscopy – general acquisition parameters

For analysis of organelles using live-cell spinning disk microscopy, cells were seeded into either 24-well 1.5 high performance glass bottom plates (Cellvis, P24-1.5H-N) or µ-Slide 8-well, glass bottom plates (ibidi, #80807) and further cultured in the vessel until reaching appropriate confluency for microscopy. Before microscopy, cells were washed in 1x PBS and imaged in FluoroBrite DMEM media. Cells were imaged using a Yokogawa CSU-X1 spinning disk confocal on a Nikon Eclipse Ti2-E motorized microscope. The system is equipped with a Tokai Hit stage top incubator and imaging was performed at 37°C, 5% CO_2_ and 95% humidity under a Nikon Plan Apo 60×/1.40 N.A immersion oil objective lens. Fluorophores were excited in sequential manner with a Nikon LUN-F XL solid state laser combiner ([laser line – laser power]: 405 nm -80mW, 488 nm - 80 mW, 561 nm - 65 mW, 640 nm - 60 mW using a Semrock Di01- T405/488/568/647 dichroic mirror. Fluorescence emissions were collected through a Chroma ET455/50m [405 nm], Chroma ET525/36m [488 nm], Chroma ET 605/52m [561nm] and a Chroma ET700/75m [for 640 nm] filters, respectively (Chroma Technologies). Images were acquired with a Hamamatsu ORCA-Fusion BT CMOS camera (6.5 µm^2^ photodiode, 16- bit) camera and NIS-Elements image acquisition software. Consistent laser intensity and exposure time were applied to all the samples, and brightness and contrast were adjusted equally by applying the same minimum and maximum display values in ImageJ/FiJi.^54^ Image quantification was performed in ImageJ/FiJi using custom-written batch-macros.

#### Live-cell microscopy for mitochondrial membrane potential measurements

For measuring mitochondrial membrane potential in live-cells, HeLa^TMEM192-HA^ control and mutant cell lines were seeded in µ-Slide 8 well chambers and treated according to the experimental plan. Before imaging, cells were incubated with TMRE (1:5000) and MitoTrackerDeepRed (1:10000) for 1 h at 37°C, washed twice with PBS and growth media replaced before imaging. 5 % laser power and 100 ms (568 nm) or 50 ms (640 nm) exposure time was used to image 6 µm z-stacks of cells. Mitochondrial masks were created based on the MitoTracker-DeepRed signal and TMRE intensities measured within these masks for evaluation.

Measurement of lysosomal pH using live-cell spinning disk microscopy

Day before measurements, 100,000 cells were seeded in 24-well glass bottom plate (Cellvis). On the day of measurement, cells were loaded with SiR-Lysosome (1:1000, Cytoskeleton Inc.) and pHLys Red (1:1000, Dojindo) for 1 h in DMEM + 10% FBS. Stains was then washed out and chased with phenol-red free DMEM+10% FBS for 3 h before imaging. For BafA1 treatment,

1 µM BafA1 was treated 2.5 h into the chase for 30 minutes prior to live-cell imaging on confocal microscope with 20x objective. To establish the pH calibration curve, wildtype cells were bathed in calibration buffers with pH adjusted to 3, 4, 5, 6, and 7, supplemented with 10 µM monensin.^81^ For both experimental and pH calibration conditions, 5 to 6 field of views were imaged and analyzed in its entirety. This process was repeated for each independent experiment. Image analysis was performed using Fiji (ImageJ). For each field of view, background subtraction was processed using the rolling ball background subtraction method for each channel. Subsequently, Otsu’s method was used to threshold the SiR-Lysosome signal to select region of interest (ROI) corresponding to lysosomes. The selected ROI was applied to the pHLys Red channel and then measured fluorescence intensity. The fluorescence intensity of the pHLys Red channel was then fitted to the calibration curve to calculate pH value.

#### Measurement of FAC addback on lysosomes and mitochondria in iNeuron using spinning disk microscopy

For measurement of lysosomal morphology ± FAC in iNeurons, hESC were seeded on day 4 of differentiation µ-Slide 8-well, glass bottom plates (ibidi, #80807) and further differentiated in the vessel for the indicated times until reaching appropriate confluency and imaged. Iron was supplemented every second day. iNeurons were imaged using a spinning disk confocal microscope (see details above) using a Nikon Plan Apo 60×/1.40 N.A immersion oil objective lens. 5% laser power and 100 ms (568 nm) exposure time was used to image 6 µm z-stacks of cells. Image quantification was performed in CellProfiler. LysoTracker Signal was segmented using global Otsu thresholding and the object shape / number measured. Statistical analysis and plotting of microscopy data was performed in Prism.

Measurement of MLSA5 long-term treatment on lysosomes in iNeuron using spinning disk microscopy

For measurement of lysosomal number after long-term MLSA5 treatment in iNeurons, hESC were seeded on day 4 of differentiation µ-Slide 8-well glass bottom plates (ibidi, #80807) and further differentiated in the vessel for the indicated times until reaching appropriate confluency and imaged. 10 µM MLSA5 was supplemented every other day. Cells were treated with 10µM Oregon Green 488 BAPTA-1, AM, for 16 h (overnight) as well as incubated for 1 h at 37°C with LysoTrackerRed (1:10000). Cells were carefully washed once, and fresh growth media added to the wells. iNeurons were imaged using a spinning disk confocal microscope (see details above) using a Nikon Plan Apo 60×/1.40 N.A immersion oil objective lens. 200 ms exposure time @ 50% laser power (488nm) and 100 ms exposure time @ 10% laser power (568nm) and 1 min time-lapses with 60 timepoints were used to image the cells. Image quantification was performed in CellProfiler. Statistical analysis and plotting of microscopy data was performed in Prism.

#### Ca^2+^ release from lysosomes in HeLa cells using spinning disk microscopy

Measurement of lysosomal Ca^2+^ content after acute CASM initiation using MLSA5, Hela Control and *NPC1^-/-^* were into µ-Slide 8-well glass bottom plates (ibidi, #80807). After reaching appropriate confluency cells were treated with 10 µM Oregon Green 488 BAPTA-5N, for 16 h (overnight) as well as incubated for 1 h at 37°C with LysoTrackerRed (1:10000). Cells were incubated for 1 h with 10 µM MLSA5 before being carefully washed twice with PBS and fresh growth media added to the wells. HeLa cells were imaged using a spinning disk confocal microscope (see details above) using a Nikon Plan Apo 60×/1.40 N.A immersion oil objective lens. 200 ms exposure time @ 20% laser power (488nm) and 200 ms exposure time @ 10% laser power (568nm) was used to image 2 µm z-stacks of cells. Image quantification was performed in CellProfiler. Statistical analysis and plotting of microscopy data was performed in Prism.

#### Immunocytochemical analysis

HeLa cells were fixed with warm 4% paraformaldehyde (Electron Microscopy Science, #15710, purified, EM grade) in PBS at 37°C for 30 minutes and permeabilized with 0.5 % Triton X-100 in PBS for 15 minutes at room temperature. After three washes with 0.02% Tween20 in PBS (PBST), cells were blocked for 10 minutes in 3% BSA-1xPBS at room temperature and washed again three times in PBST. Cells were incubated for 3 h in primary antibodies in 3% BSA- 1xPBS and washed three times with PBST. Secondary antibodies (Thermo Scientific, 1:400 in 3% BSA-1xPBS) where applied for 1 h at room temperature. To stain nuclei, Hoechst33342 (1:10000) was added for 5 minutes to cells in PBST and finally washed three times. Filipin staining was performed after fixation for 2 h at room temperature in PBS (0.05 mg/ml). Primary and secondary antibodies used in this study can be found in the **Key Resource Table**.

#### Fixed-cell microscopy – general acquisition parameters

Immunofluorescently labelled Hela or iNeurons (antibodies indicated in figures and figure legends and details in **Key Resource Table**.) were imaged at room temperature using a Yokogawa CSU-W1 spinning disk confocal on a Nikon Eclipse Ti2-E motorized microscope equipped with a Nikon Plan Apochromat 40×/0.40 N.A air-objective lens, Nikon Plan Apochromat 60×/1.42 N.A oil-objective lens and a Plan Apochromat 100×/1.45 N.A oil-objective lens. Signals of 405/488/568/647 fluorophores were excited in sequential manner with a Nikon LUN-F XL solid state laser combiner ([laser line – laser power]: 405 nm - 80 mW, 488 nm - 80 mW, 561nm - 65 mW, 640 nm - 60 mW using a Semrock Di01-T405/488/568/647 dichroic mirror. Fluorescence emissions were collected with Chroma ET455/50m [405 nm], 488 Chroma ET525/50m [488 nm], 568 Chroma ET605/52m [561 nm], 633 Chroma ET705/72m [640 nm] filters, respectively (Chroma Technologies). Confocal images were acquired with a Hamamatsu ORCA-Fusion BT CMOS camera (6.5 µm^2^ photodiode, 16-bit) camera and NIS-Elements image acquisition software. Consistent laser intensity and exposure time were applied to all the samples, and brightness and contrast were adjusted equally by applying the same minimum and maximum display values in ImageJ/FiJi.^54^

#### Evaluation of Ferritin accumulation in lysosomes

The quantitative measurement of FTH1 accumulation inside the lysosomal mask was performed by seeding HeLa^TMEM192-HA^ control and mutant cell lines into 24-well 1.5 high performance glass bottom plates (Cellvis, 24-1.5H-N) and treated according to the experimental plan. Fed and treated cells were fixed according to the procedure stated above, stained for ferritin (FTH1), lysosomes (HA) and DNA (SPY-DNA555) and imaged using a Nikon Plan Apochromat 40×/0.40 N.A air-objective lens. 12 randomly selected positions (8 µm z-stacks) were acquired were acquired using the HCA-module in NIS-Elements. For image analysis, CellProfiler^82^ was used for the quantitative analysis of FTH1 colocalization with the HA-derived lysosomal mask. Plotting of microscopy data was performed in Prism. Primary and secondary antibodies used in this study can be found in the **Key Resource Table**.

#### Effect of GFP-SopF expression (CASM) on ATGlyation using spinning disk microscopy

HeLa^TMEM192-HA^ cells were seeded into 24-well 1.5 high performance glass bottom plates (Cellvis, 24-1.5H-N, 20k per well). The next day, GFP-SopF was transfected using FuGene HD transfection reagent (Promega) according to manufacturer protocol and a full media exchange was performed the day after. Two days after the transfection, cells were treated for 24 h with 2 µM U18666A to block NPC1 function. Cells were fixed using ice-cold methanol (10 minutes, for LC3B) or paraformaldehyde (30 minutes, for panGABARAP) and immunofluorescently labelled for GFP, Lamp1 and LC3B / panGABARAP. Images were acquired using a Yokogawa CSU-W1 spinning disk confocal on a Nikon Eclipse Ti2-E motorized microscope using a Plan Apochromat 60×/1.42 N.A oil-objective lens. Images were analysed using CellProfiler. Statistical analysis and plotting of microscopy data was performed in Prism. Primary and secondary antibodies used in this study can be found in the **Key Resource Table**.

### 3D STRUCTURED-ILLUMINATION-MICROSCOPY

#### Fixed cell 3D-SIM sample preparations

Fixed cell 3D-SIM samples were prepared as described.^83^ Briefly, HeLa^TMEM192-HA^ control and mutant cell lines were seeded on 18x18 mm Marienfeld Precision cover glasses thickness No.1.5H (tol. ± 5 μm) and cultured at the indicated conditions / treatments. To mimic NPC1-loss Control cells were treated with 2 µM U18666A for the indicated timepoints in growth medium. Cells were fixed at 37°C in 4% paraformaldehyde (Electron Microscopy Science) for 30 minutes and permeabilized for 15 minutes with 0.5% Triton X-100 in PBS at room temperature. After three washes with 0.02% Tween20 in PBS (PBST), cells were blocked for 10 minutes in 3% BSA-1xPBS at room temperature and washed again three times in PBST. If required, cholesterol molecules were labeled with Filipin for 2 h room temperature in 1x PBS (0.1 mg/ml), before washing the sample with PBST twice to remove excess label. Primary antibody incubation was performed over night at 4°C with gentle rocking in 3% BSA – 1x PBS, followed by three 5 minutes washes with PBST. Secondary antibody incubation (1:400 in 3% BSA-1x PBS) was performed at room temperature for 1 h with gentle rocking. Samples were washed three times for 5 minutes in 1xPBST. Before mounting on glass slides, coverslips were washed once in 1xPBS and mounted in Vectashield (Vector Laboratories, H-1000-10). Primary and secondary antibodies used in this study can be found in the **Key Resource Table**.

#### 3D-SIM microscopy – acquisition parameters

3D-SIM microscopy was performed on a DeltaVision OMX v4 using an Olympus 60x / 1.42 Plan Apo oil objective (Olympus, Japan). The instrument is equipped with 405 nm, 445 nm, 488 nm, 514 nm, 568 nm and 642 nm laser lines (all >= 100 mW) and images were recorded on a front-illuminated sCMOS (PCO Photonics, USA) in 95Mhz, 512x512px image size mode, 1x binning, 125 nm z-stepping and with 15 raw images taken per z-plane (5 phase-shifts, 3 angles). Raw image data was computationally reconstructed using CUDA-accelerated 3D-SIM reconstruction code (https://github.com/scopetools/cudasirecon) based on.^84^ Optimal optical transfer function (OTF) was determined via an in-house build software, developed by Talley Lambert from the NIC / CBMF (GitHub: https://github.com/tlambert03/otfsearch, all channels were registered to the 528 nm output channel, Wiener filter: 0.002, background: 90). ChimeraX was used for 3D renderings if imaging data.

#### Live-cell 3D-SIM sample preparations

MatTek 35 mm Dish, High Precision 1.5 Coverslip were coated for 2 h with poly-L-lysine a 37°C before washing excess solution off with three 1xPBS washes. Cells were seeded in dishes and cultured / treated as indicated. On day of experiment, cells were incubated with PKmitoRed (1:1000) for 1 h at 37°C washed with warm medium to remove excess dye. For assessing fusion-competency of lysosomes in NPC2^-/-^ mutants, cells were seeded in MatTek 35 mm dishes (see above) and loaded with Alex647-conjugated Dextran o/n (1:200 dilution) at 37°C. The next day, cells were stained with LysoTracker Red DND-99 for 1 h at 37°C (1:5000), washed twice with PBS and medium replaced with fresh, warm growth medium. If necessary, 2 µM U18666A was added to Control cells for the indicated times to block NPC1 function.

#### Live-cell 3D-SIM microscopy – acquisition parameters

3D-SIM microscopy was performed on a DeltaVision OMX v4 using an Olympus 60x / 1.42 Plan Apo oil objective (Olympus, Japan). The instrument is equipped with 405 nm, 445 nm, 488 nm, 514 nm, 568 nm and 642 nm laser lines (all >= 100 mW) and images were recorded on a front-illuminated sCMOS (PCO Photonics, USA) in 286Mhz, 512x512px image size mode, 1x binning, 125 nm z-stepping and with 15 raw images taken per z-plane (5 phase-shifts, 3 angles). ∼10 %T laser was used at 5-20 ms exposure times. 0.750 - 1 µm (for mitochondria) or 2 µm (for lysosomes) thick z-stacks were recorded for each timepoint / field of view. Raw image data was computationally reconstructed using CUDA-accelerated 3D-SIM reconstruction code stated above.

### ELECTRON MICROSCOPY

HeLa^TMEM192-HA^ Control and mutant cells were grown on Aclar plastic coverslips in above stated growth conditions until 70-80% confluency was reached, washed twice in 1x PBS and fixed with a fixation mixture of 2% formaldehyde and 2.5% glutaraldehyde in 0.1 M Sodium Cacodylate buffer, pH 7.4 for 1 h at room temperature. Sample preparation and microscopy was performed by the Harvard Medical School Electron microscopy facility (https://electron-microscopy.hms.harvard.edu/methods).

### CRYO-PLASMA FIB - CRYO-ELECTRON TOMOGRAPHY (CRYO-ET)

#### Cryo-ET sample preparation and freezing

HeLa^TMEM192-HA^ Control and *NPC2^-/-^* cells were cultured on EM grids as follows: 200-mesh gold grids with Silicon Dioxide R1/4 film (Quantifoil) were plasma cleaned, coated by incubation with 1 mg/mL Poly-L-Lysine (Sigma, P2636) solution in 0.1 M Borate Buffer (pH 8.5 in distilled water, autoclaved) for 2 h and washed twice with PBS. One day before plunging, a 150 μL drop of ∼150 cells/µL was added on top of each grid and placed in a well of a 4-well 35 mm cell culture dish (Greiner bio-one, 627170); after 2 h of settling time, DMEM medium was added to a final volume of 2 mL per dish. The next day, cells were starved for 6 h in phenol red-free EBSS and 10% glycerol was added to the medium few minutes before plunging. Samples were plunged into ethane/propane with a Vitrobot Mark IV (Thermo Fisher Scientific), with application of 4 μL of EBSS medium and with the following settings: room temperature, humidifier 70 %, blot force 8, blot time 9 seconds. After plunging, the grids were clipped into autogrids with cutout for FIB-milling in a custom clipping station.^85^

#### Focused ion beam (FIB) milling

TEM-transparent lamellae were produced in a commercially available Arctis cryo-Plasma Focused Ion Beam (cryo-PFIB) instrument (Thermo Fisher Scientific, Eindhoven, The Netherlands) equipped with a robotic sample delivery device (termed “Autoloader”), compustage, NiCol-scanning electron microscope (SEM) column and Tomahawk-focussed ion beam (FIB) column. Pre-clipped grids were assembled in the standard multispecimen cassette holder so that the cutouts later face the ion beam on the compustage. A Xenon ion beam was used for all described steps. After powering on and aligning the beams in the XT user interface, all consecutive steps were carried out using the proprietary Arctis WebUI software (Version 1.0).^85,86^ The grid template sets the parameters for the initial mapping of the grid from the SEM or FIB as well as for the initial protective coating, and the final sputter fiducials on the polished lamella. We used here a modified version of the “Electron tileset with auto deposition”. First, a tiled overview perpendicular to the grid was acquired with the SEM with a dwell time of 3 μs and a horizontal field width of 256 μm. Then, points of interest (POIs) were placed at suitable cell positions manually. To protect the leading edge of these positions while milling, a three-step protocol of sputter-, chemical vapor-, and sputter deposition was carried out. The sputtering process was executed by milling a calibrated regular pattern into an in-built platinum target with a 12 kV Xenon beam at a current of 70 nA for 120 seconds in order to deposit a thin film of atomic platinum. The platinum fiducials on the end of lamella preparation were induced by the same process but milling only for 5 seconds. Chemical vapor deposition was executed by heating the attached gas injection system to 28°C and opening the shutter for 50 seconds in order to deposit an organo-metallic layer of trimethyl(methylcyclopentadienyl)-platinum(IV) with a thickness of several µm on the sample.

The lamella template sets the imaging and milling parameters for the retrieval of the POIs in the ion beam, the milling angle search, the ion beam milling, and the final image acquisition. The set final lamella thickness was refined to 120 ± 10 nm depending on the ice thickness of the respective grid. Briefly, the 30 keV Xenon beam milling procedure was as follows: eucentric height and the maximum milling angle of -18° were refined before milling 0.5 µm wide stress relief cuts at a distance of 10 µm to each side of the intended lamella using a 1.0LJnA ion beam. Three milling steps were then used to remove material above and below the intended lamella position: (i) rough milling at 3.0LJnA to 1 μm thickness, (ii) 0.3LJnA to 500 nm, and (iii) 0.1LJnA to the 300 nm. The respective Silicon depth correction factors were (i) 0.4, (ii) 0.7, and (iii) 0.88. Afterwards, the lamella was polished at a current of 30 pA to 110-130 nm. In some cases, remnants of the cell top surface with its organometallic layer had to be removed in addition to make the full tilt range in cryo-ET accessible.

#### Cryo-ET data acquisition and Processing

TEM data acquisition was performed on a Krios G4 at 300 kV with Selectris X energy filter and Falcon 4i camera (Thermo Fisher Scientific, Eindhoven, The Netherlands) using Tomo5 (Version 5.12.0, Thermo Fisher Scientific). Tilt series were acquired at a nominal magnification of 42,000X (pixel size 2.93 Å) using a dose-symmetric tilt scheme with an angular increment of 2°, a dose of 2 e^-^/Å^2^ per tilt and a target defocus between -3 and -6 μm. Tilt series were collected ranging from -48° to +60° relative to the lamella pretilt., and frames were saved in the EER file format. The positions for tilt series acquisition were determined by visual inspection of 11500X magnification “search” montage maps acquired in thin areas of the sample. For publication display, search maps were cleaned and destriped using the Fiji LisC macro and the contrast was enhanced using Contrast Limited Adaptive Histogram Equalization^87^ (https://github.com/FJBauerlein/LisC_Algorithm, **Figure 4C**). Tilt series were acquired of lysosome-like structures. Tilt series frames were motion-corrected with Relion’s implementation of Motioncorr2^88^ for EER files.^59^ Alignment and CTF-correction was performed in IMOD^89^ (v.4.10.49, RRID:SCR_003297, https://bio3d.colorado.edu/imod/) and reconstruction by AreTomo^90^ (v.1.3.3) by using an adjusted version of the TomoMAN wrapper scripts (https://doi.org/10.5281/ZENODO.4110737). Tomograms at 2×binning (IMOD bin 4) with a nominal pixel size of 11.72 Å were denoised using cryo-CARE (https://github.com/juglab/cryoCARE_T2T).^91^

#### Cryo-ET dataset annotation and analysis

Membrane thickness was measured in Gwyddion^92^ (v.2.63, http://gwyddion.net/) from unbinned, ctf-corrected tomograms after preprocessing with IMOD and EMAN.^93^ To achieve the necessary contrast, tomograms were oriented in the 3Dmod slicer so that the interleaflet space was optimally visible. Tomograms were then rotated, low-pass filtered to the Nyquist frequency (EMAN2, v.2.99.47, https://blake.bcm.edu/emanwiki/EMAN2), and trimmed to the multilamellar bodies’ volumes. Then, tomograms were averaged in z along the slices containing visible membranes and converted to a single 16-bit TIF image. In Gwyddion, the tomogram was inverted and its minimum intensity value shifted to zero. Afterwards, profiles of similar length were extracted along arbitrary lines perpendicular to the membrane. At least four distinct profiles were placed across the whole membrane, each averaging 64 pixels perpendicular to the drawn line. Afterwards, the individual profiles were averaged in OriginLab 2023. Subsequently, peaks were detected and their full width at half maximum (FWHM) and peak-to-peak distance was analyzed automatically. For attenuated peaks, the peak- as well as the FWHM-positions were refined manually. Plots were created in Prism.

#### Tomogram segmentation

All membranes in the tomogram were segmented with Membrain-Seg^94^ (https://github.com/teamtomo/membrain-seg) using the publicly available pretrained model (v9), with the exception of the membranes of the multilamellar vesicles (MLV), which were segmented with TomosegmemTV^94,68^ (v.1.0, https://sites.google.com/site/3demimageprocessing/tomosegmemtv) due to their aberrant membrane spacing. All segmentations were then merged and manually refined in Amira.3D (Thermo Fisher Scientific), and final renderings were generated in ChimeraX.^70^

### DATA AVAILABILITY

The data, code, protocols, and key lab materials used and generated in this study are listed in the Key Resource Table alongside their identifiers / catalog numbers on Zenodo (10.5281/zenodo.13905094). Scripts and tabulated data related to this work are deposited on Zenodo (10.5281/zenodo.13905094). Proteomic data and analysis files part of this study are deposited at ProteomeXchange Consortium by the PRIDE partner.^95^ The PRIDE project identification number is PXD049336 (TMTpro experiments). Raw files collected by nMOST are available on MassIVE, assession number: MSV000094201. Raw files collected by nDIA are available on MassIVE, assession number: MSV000095985. Tomograms are deposited on EMDB under the accession numbers EMD-51701 and EMD-51702. The lipidomic and proteomic data viewer is available at: https://coonlabdatadev.com/ (Username: most_lsd; Password: viewer).

## SUPPLEMENTAL FIGURE LEGENDS

**Figure S1:**
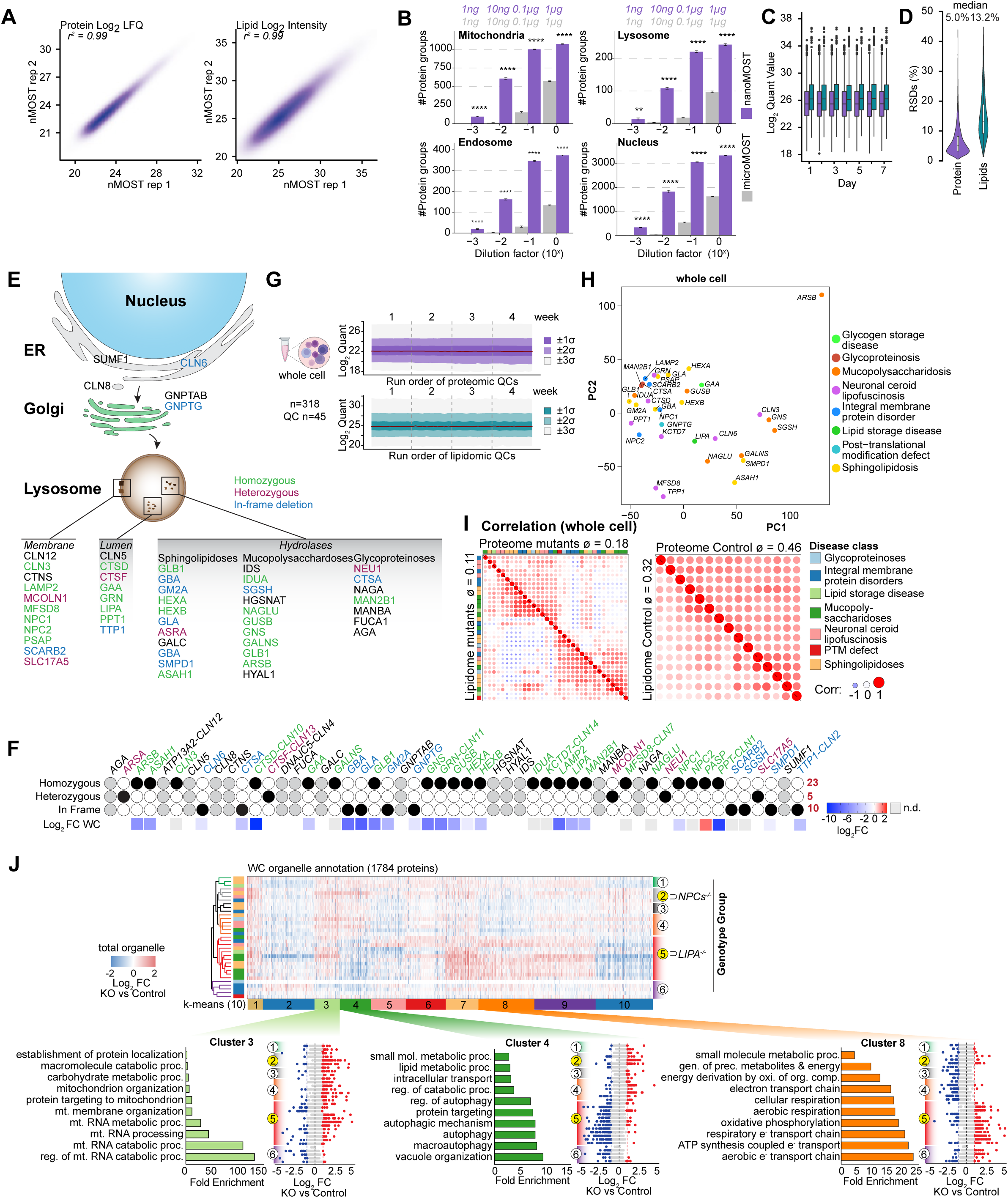
Benchmarking of nMOST and application to cells lacking LSD genes. **(A)** Correlation of log_2_ label-free quantification (LFQ) protein (left panel) and lipid biomolecules (right) of two nMOST runs. **(B)** Direct performance comparison of nMOST (purple) with µMOST (grey) over 4 magnitudes of sample dilution. Injection amounts for nMOST and µMOST are listed above. Number of protein groups identified by selected organelles are plotted. **(C)** Quantification of number of log_2_ quant value (protein and lipid) over a 7-day acquisition period using nMOST. **(D)** Violin plot depicting % relative standard deviations (RSDs) for both quantified protein and lipid identifications over a 7-day acquisition period using nMOST. **(E)** Schematic summarizing 52 LSD proteins and their localization properties when known **(F)** Summary of gene editing champaign with the goal of creating mutants across LSD genes in HeLa^TMEM192-HA^ cells. Black circles indicate the status of mutants obtained. Gray circles indicate no clones for the indicated genes. Lower panel shows log_2_FC for all detected LSD proteins in either whole cell extracts from the indicated mutant cell line based on nMOST. **(G)** Log_2_ LFQ for total proteomes/lipidomes for 363 total samples analyzed over a 4-week data collection session (318 whole cell extracts with each LSD mutant, untagged Control HeLa and HeLa^TMEM192-HA^ control cells all in quadruplicate biological replicates, and 45 MS QC samples). **(H)** PCA blots for combined proteome and lipidome across the LSD mutants in this study. **(I)** Heatmap depicting correlation of proteome and lipidomes for LSD mutants (top) and controls (bottom). **(J)** Heatmap (log_2_FC [Mutant / Control]) of average proteome abundance across the indicated organelle of LSD mutants.

**Figure S2:**
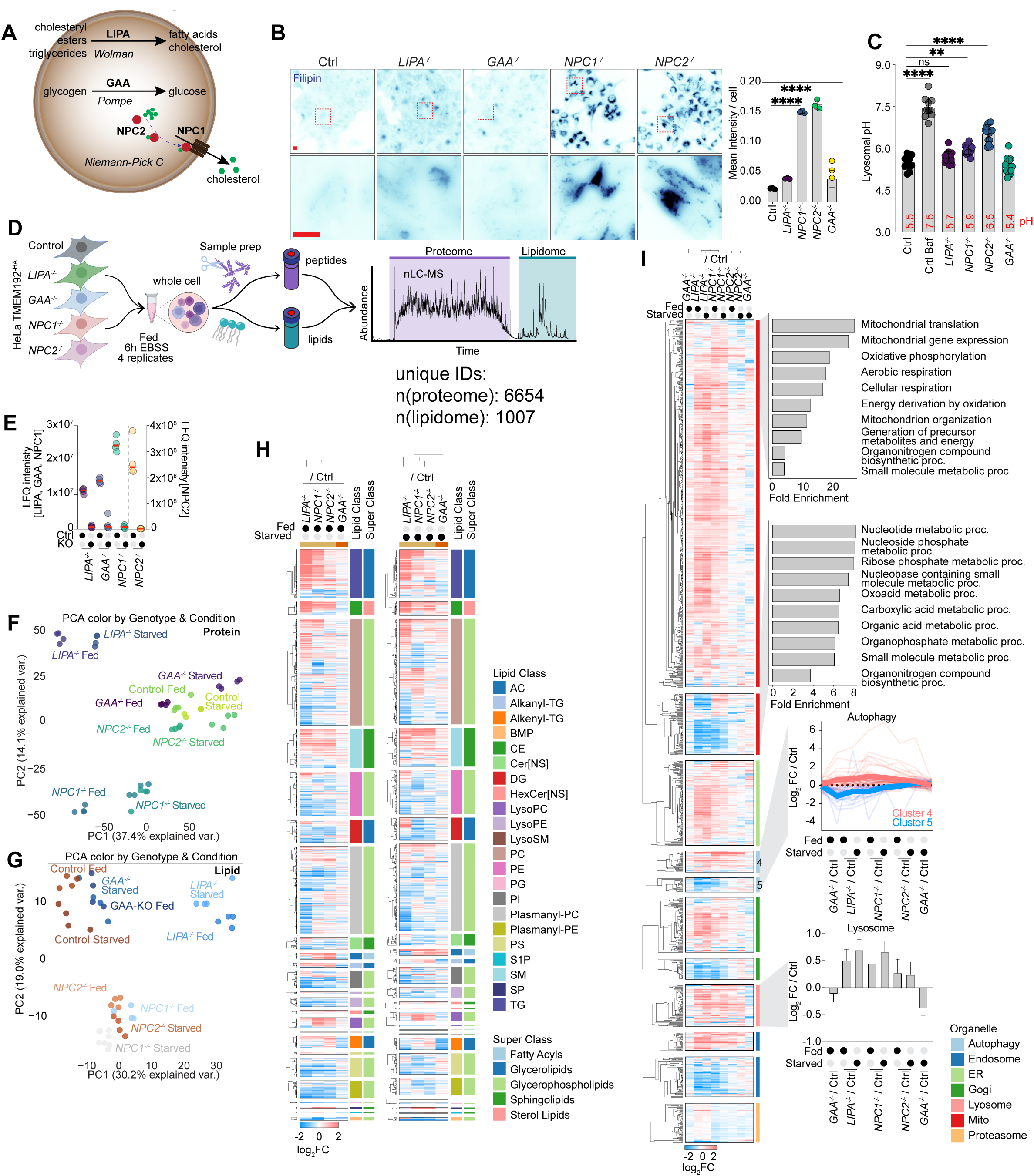
4KO-nMOST for profiling autophagy defects in LSDs involved in cholesterol metabolism. **(A)** The general functions of the four proteins selected for 4KO study (*LIPA*^-/-^, *GAA*^-/-^, *NPC1*^-/-^ and *NPC2*^-/-^) within the lysosome is shown in the schematic. **(B)** Wide-field fluorescent images of Hela Control and 4KO cell lines stained for cholesterol with Filipin. Quantification of mean Filipin intensity per cell is plotted below (data from three biological replicates, 20 image stacks per repeat; genotype(N)): Ctrl(1424), *LIPA*^-/-^(2123), *GAA*^-/-^ (1791), *NPC1*^-/-^ (1773), *NPC2*^-/-^ (3270)). p(****) <0.0001, ordinary two-way ANOVA with multiple comparisons, alpha = 0.05; error bars depict S.D. Scale bar = 20 µm. **(C)** pH measurements for 4KO cells using ratiometic confocal imaging. Each data point represents one field of view; Repeats per genotype (N): Ctrl(12), Ctrl + BafA(10), *LIPA*^-/-^, *GAA*^-/-^, *NPC1*^-/-^, *NPC2*^-/-^(15). p(****) <0.0001; p(**) = 0.0022; ordinary one-way ANOVA with multiple comparisons, alpha = 0.05; error bars depict S.D. **(D)** Application of nMOST for analysis of Control, *LIPA*^-/-^, *GAA*^-/-^, *NPC1*^-/-^ and *NPC2*^-/-^ cells (4KO cells). The 5 indicated HeLa^TMEM192-HA^ cell lines were analysed in quadupliates for both fed and starvation conditions. Number of unique IDs for proteins and lipids are shown under the chromatograph. **(E)** LFQ of LIPA, GAA, NPC1 and NPC2 in Control and mutant cells based on nMOST data. Data based on quadruplicate replicate nMOST measurements. **(F,G)** PCA analysis of 4KO proteomic (panel A) and lipidomic (panel B) data from nMOST analysis of the indicated cell lines under Fed or EBSS (6 h) conditions. Data based on quadruplicate biological replicate nMOST measurements. **(H)** Heatmap of log_2_ abundance of lipids under indicated treatment conditions. Lipid classes / super classes are highlighted on the right of the heatmap. Data based on quadruplicate biological replicate nMOST measurements. **(I)** Heatmap of log_2_ abundance of organelle-annotated proteins under indicated treatment conditions. Top two graphs show results of GO-term enrichment analysis associated with the two mitochondrial clusters. Abundance for autophagy clusters 4 and 5 are plotted on the right. Abundance of lysosome is plotted at the bottom right bar-graph. Data based on quadruplicate biological replicate nMOST measurements.

**Figure S3:**
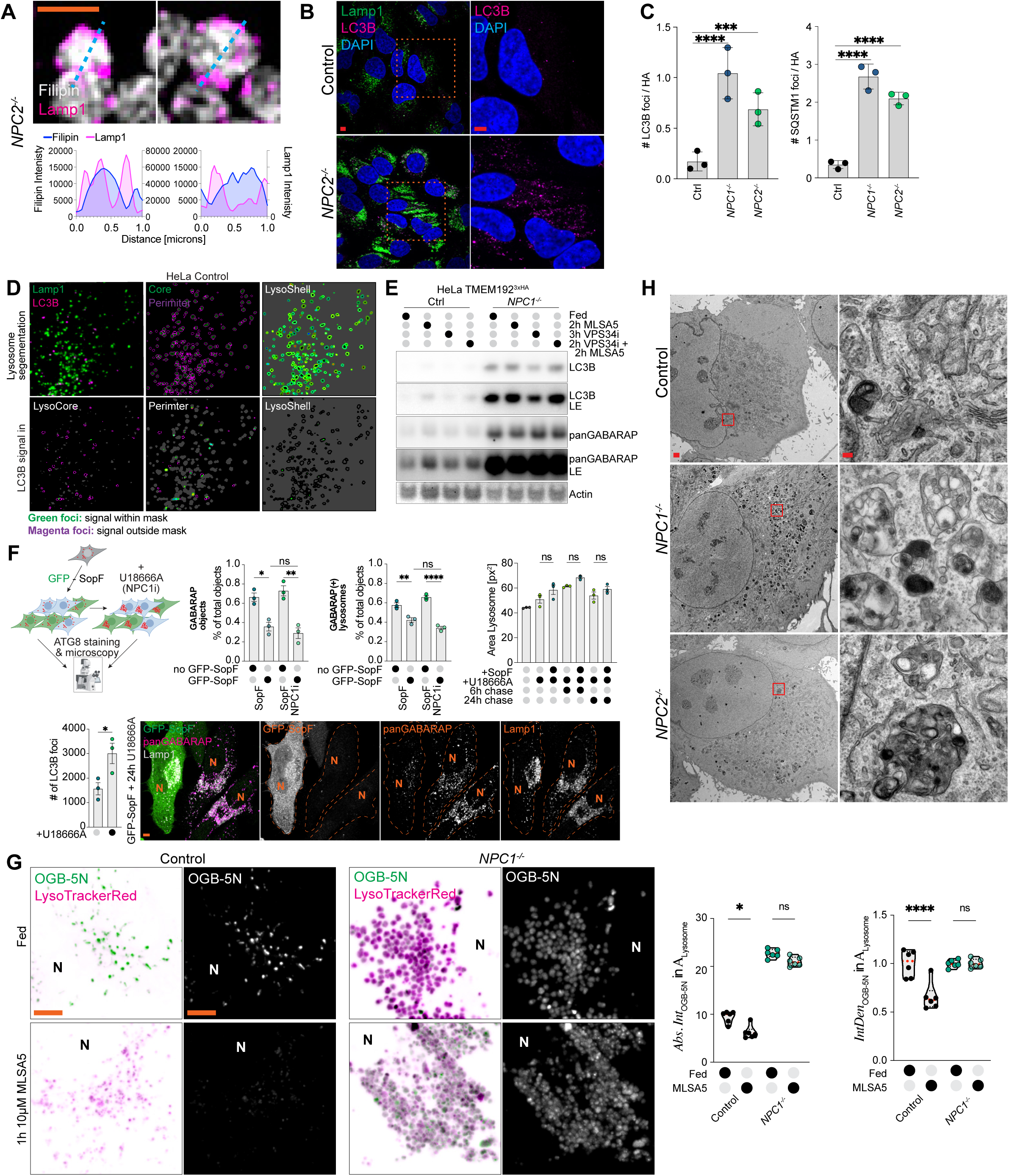
Profiling of lysosomal function in NPC1 and NPC2 mutant cells. (A) Example of Filipin-positive lysosomes and α-LAMP1 signal derived from 3D-SIM imaging. Scale bar = 1 µm. Lineplot of filipin and LAMP1 intensities for the dashed line are plotted beneath. (B) Immunostaining of Control and *NPC2*^-/-^ cells with α-LAMP1 and α-LC3B. Nuclei were stained with DAPI. Scale bars = 5 µm. **(C)** Evaluation based on confocal imaging of Control, *NPC1*^-/-^ and *NPC2*^-/-^ cells immunostained with α-LC3, α-SQSTM1, and α-HA to detect TMEM192^-HA^. Quantification was performed on three biological replicates with 5 stacks in each replicate. MAPLC3B: p(****) <0.0001; p(***) = 0.0002; p(*)=0.0129 & 0.0157. p62/SQSTM1: p(****) <0.0001. Data from quadruplicate replicates, ordinary one-way ANOVA with multiple comparisons, alpha = 0.05. Error bars depict S.D. (**D**) Example for segmentation and analysis strategy for quantifying LC3B localization relative to LAMP1. Input image is segmented and filtered to create a lysosomal core & perimeter mask, and the resulting lysosomal shell (Perimeter \ Core). Underneath, example results for LC3B signal in the different localization in a HeLa Control cell is shown. Green foci depict LC3B signal that reside within the specific mask, magenta-coloured foci represent foci that are outside the specific mask. **(E)** Western Blot for select autophagy proteins of whole cell lysates from HeLa Control and *NPC1^-/-^* mutants treated with MLSA5 and/or VPS34inhibitor. **(F)** Schematic of experimental approach to study role of GFP-SopF and lysosomal ATG8lyation in relationship to NPC1 inhibition. Example confocal images of GFP-SopF expressing Hela cells immunostained for α-LAMP1 and α-panGABARAP. Scale bar = 5 µm. GABARAP per cell comparisons: A-B: p(*) = 0.0103. C-D: p(**) = 0.0011. GABARAP per lysosome comparisons: A-B: p(**) = 0.0033. C-D: p(****) = <0.0001. LC3B p(*) = 0.0180. Data from 3 replicates with 15 stacks each. One-way ANOVA with multiple comparisons, alpha = 0.05. Error bars depict S.E.M. **(G)** Images from live-cell microscopy of Control and *NPC1^-/-^*in fed and MLSA5-treated conditions. Lysosomes are stained with LysoTrackerRed and lysosomal Ca^2+^ using OGB-5N. Scale bar = 5 µm. Violin plots of quantification of lysosomal Ca^2+^ intensity ± MLSA5 treatment in absolute measures (left) and relative to lysosomal area (right). Quantification was performed on five replicates with 3 image stacks in each replicate. Ctrl: p(****) <0.0001; p(*)=0.0143. Two-way ANOVA with multiple comparisons, alpha = 0.05. **(H)** Control, *NPC1*^-/-^ or *NPC2*^-/-^ cells were examined by electron microscopy. Scale bars = 1 µm (left & middle panel) and 100 nm (right panel).

**Figure S4.**
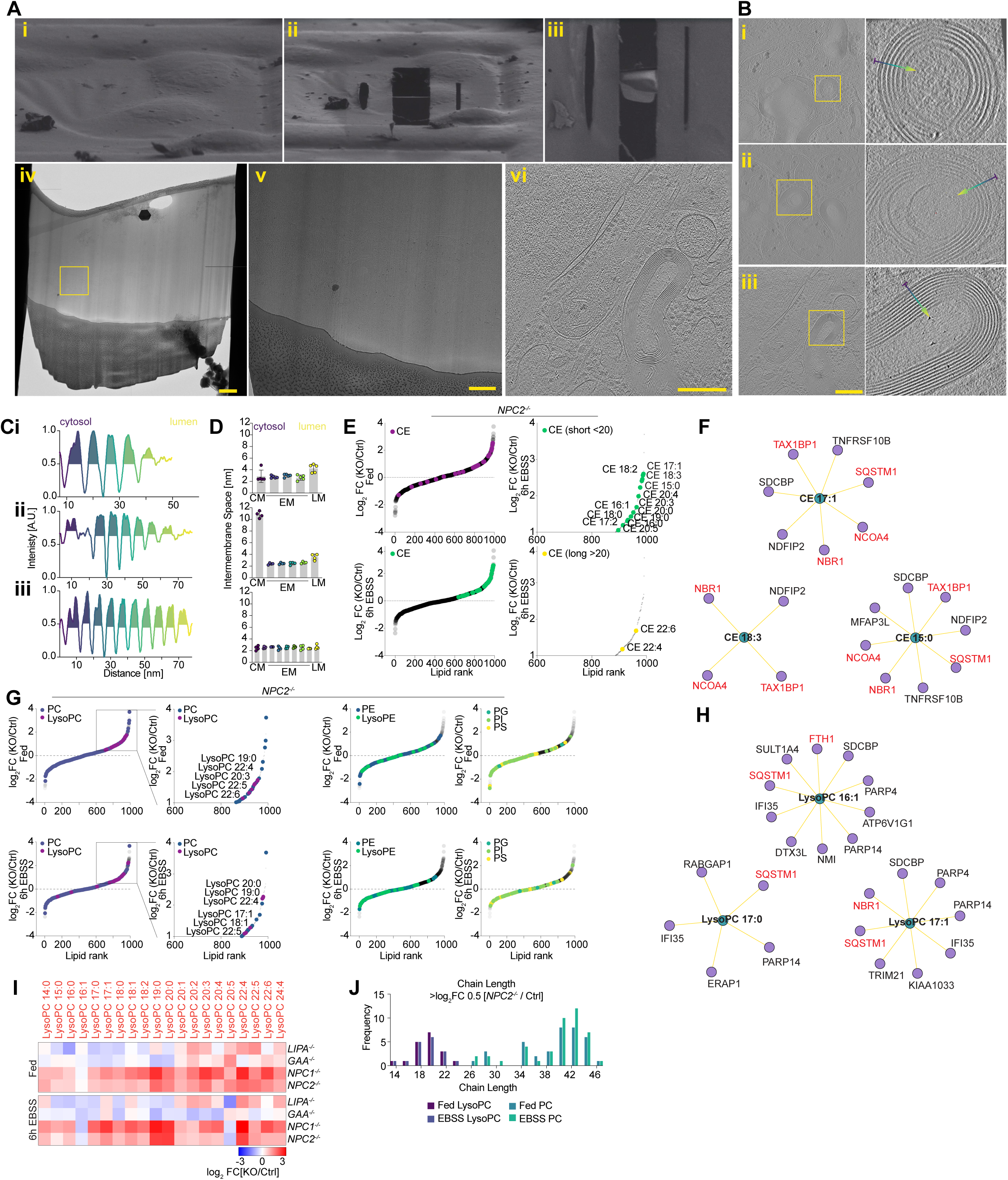
Visualization of multi-lamellar membranes in *NPC2*^-/-^ lysosomes by cryo-ET. **(A)** Example images of the cryo-PFIB and cryo-ET workflow. Vitrified cells before (i) and after (ii) lamella preparation by cryo-PFIB milling. (iii) final SEM view of milled and polished lamella. (iv-v) Lamella overview with zoom-in on a MLV containing area. (vi) Reconstructed tomogram showing an MLV. Scale: (i-ii) 156 µm horizontal field width (HFW), (iii) 124 µm HFW. Scale bar = (iv) 1 µm (v) 500 nm (vi) 250 nm. (**B**) Overview images and zoom-ins of three tomograms depicting MLV membrane stacks. Scale bar = 250 nm. **(C)** Averaged, inverted intensity along the arrows from B to determine membrane thickness. The gradient indicates the measurement direction from cytosol (purple) to lumen (yellow). Membrane peaks are coloured to indicate their full width at half maximum. **(D)** Intermembrane Space of selected MLVs between adjacent membrane pairs. The gradient indicates the measurement direction from cytosol (purple) to lumen (yellow). **(E)** Ranked lipid log_2_FC abundance of *NPC2*^-/-^ lipidome for fed and 6 h EBSS nutrient starvation conditions. Cholesterol esters (CE) are highlighted in colour on top of the overall lipidome spread. The lower row depicts ranking of short or longer CE in 6 h EBSS nutrient starvation conditions against the whole lipidome rank. Data based on quadruplicate biological replicate nMOST measurements. **(F)** Lipid-protein networks of select CE species based on cross-ome correlations of the LSD-nMOST dataset. **(G)** Ranked lipid log_2_FC abundance of phospholipids in *NPC2*^-/-^ lipidome for fed and 6 h EBSS nutrient starvation conditions. Lysosomal specific phospholipids (LysoPC/LysoPE) are highlighted in color on top of the overall lipidome and parent-lipid class spread. Lower row depicts the lipidome of annotated lipids in 6 h EBSS nutrient starvation conditions against the whole lipidome rank. Data based on quadruplicate biological replicate nMOST measurements. **(H)** Lipid-protein networks of select LysoPC species based on cross-ome correlations of the LSD-nMOST dataset. **(I)** Heatmap depicting log_2_FC of LysoPC species in either Fed and EBSS-treated Control and 4KO cells. Data based on quadruplicate biological replicate nMOST measurements. **(J)** Histogram depicting frequency of (Lyso-)PCs enriched ≥ 0.5 log_2_FC in [*NPC2^-/-^*/Control] against their chain length. Data based on quadruplicate biological replicate nMOST measurements.

**Figure S5.**
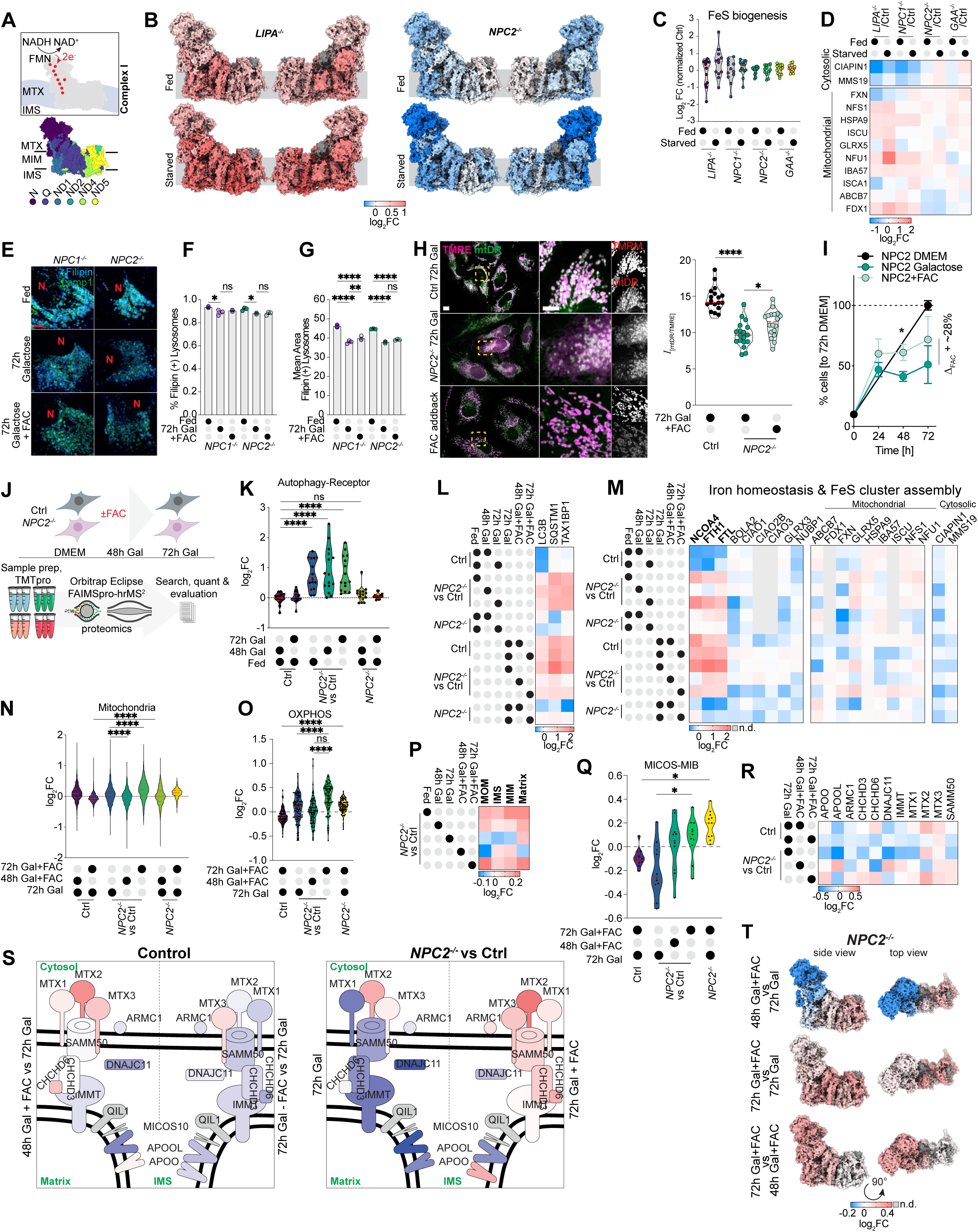
Profiling of mitochondrial proteome in 4KOs and alleviation of mitochondrial defects in *NPC2*^-/-^ cells by extracellular iron**. (A)** Schematic of CI of the OXPHOS system with individual sub-modules (PDB: 5XTH). **(B)** Log_2_FC of CI sub-module abundance in Fed and EBSS-treated conditions measured in *LIPA*^-/-^, and *NPC2*^-/-^ cells by nMOST [normalized to control]. Based on quadruplicate replicate nMOST data. Legend shows colour panel for log_2_FC values. **(C,D)** Log_2_FC Violinplot and heatmap for components of the mitochondrial and cytosolic FeS cluster biogenesis system for 4KO cells in Fed and EBSS-treated cells based on quadruplicate replicate nMOST data. **(E)** Confocal images of HeLa *NPC1^-/-^* and *NPC2^-/-^* in indicated growth media conditions immunostained with α-LAMP1 and cholesterol-rich lysosomes were stained with Filipin. Scale bar = 5 µm. **(F)** Quantification of % Filipin(+) lysosomes in different growth media conditions. Data based on three biological replicates with 6 image stack per repeat. *NPC1^-/-^*: p(*) = 0.03, *NPC2^-/-^*: p(*) = 0.0124. Unpaired t-test. Error bars depict S.E.M.. **(G)** Quantification of mean lysosomal size in different growth media conditions. Data based on three biological replicates with 6 image stack per repeat. *NPC1^-/-^*: A-B: p(****) = <0.0001, A-C: p(****) = <0.0001, B-C: : p(**) = 0.0021. *NPC2^- /-^*: A-B: p(****) = <0.0001, A-C: p(****) = <0.0001. Two-way ANOVA with multiple comparisons, alpha = 0.05. Error bars depict S.E.M.. **(H)** Stills of spinning-disk live-cell microscopy of Control and *NPC2*^-/-^ cells cultured in Galactose for 72 h or in Galactose for 72 h with FAC. Mitochondria are stained with TMRE (ΔΨm) and MitoTrackerDeepRed. Scale bar = 10 µm and 5 µm (insets). Right panel shows quantification of *I*_[mtDR-TMRE]_ for indicated genotypes and treatments. Data from four biological replicates (18 stacks per replicates); p(****) < 0.0001, p(*) = 0.0150; unpaired t.test. **(I)** % of cell count of *NPC2^-/-^*[relative to growth in 72 h DMEM] in Galactose ± FAC over a 72 h growth period. **(J)** Schematic of TMT proteomics workflow for analysis of the effect of FAC addition to Control or *NPC2*^-/-^ cells. **(K,L)** Violin plot of all autophagy receptors (panel K) and heatmap of LC3B, SQSTM1 and TAX1BP1 (panel L) for log_2_FC [*NPC2*^-/-^/Control] in cells cultured in Galactose in the presence or absence of FAC. p(****) < 0.0001; data based on biological triplicate TMTpro measurements. **(M)** Heatmap of log_2_FC [*NPC2*^-/-^/Control] for components of the cytosolic and mitochondrial FeS cluster assembly system as well as the Ferritin system with or without FAC with cells grown in Galactose. Data based on biological triplicate TMTpro measurements. **(N)** Violin plots for total mitochondrial proteins in Control versus *NPC2*^-/-^ cells grown in Galactose ± FAC. p(****) < 0.0001; data based on biological triplicate TMTpro measurements. **(O)** Violin plot of OXPHOS subunit log_2_FC values in *NPC2*^-/-^ versus Control cells grown in Galactose with or without FAC. p(****) < 0.0001, ordinary one-way ANOVA with multiple comparisons, alpha = 0.05; data based on triplicate biological replicate TMTpro measurements. **(P)** Heatmap depicting log_2_ FC of components of different mitochondrial compartments in cells cultured in Glucose and Galactose in the presence or absence of FAC. Data based on biological triplicate TMTpro measurements. **(Q)** Violin plots of log_2_FC [*NPC2*^-/-^/Control] of MICOS-MIB subunits in response to FAC. Data based on biological triplicate replicate TMTpro measurements; p(*) = 0.0162. **(R)** Heatmap of log_2_FC [*NPC2*^-/-^ /Control] for individual MICOS-MIB subunits in response to FAC. Data based on triplicate biological replicate TMTpro measurements. **(S)** Schematic showing alterations in various MICOS-MIB subunits in response to FAC. Colour coding is based on log_2_FC scale in panel R. **(T)** Log_2_FC of CI abundance in Galactose with or without FAC addback for *NPC2*^-/-^ cells. Legend shows color panel for log_2_FC values. Data based on triplicate biological replicate TMTpro measurements.

**Figure S6.**
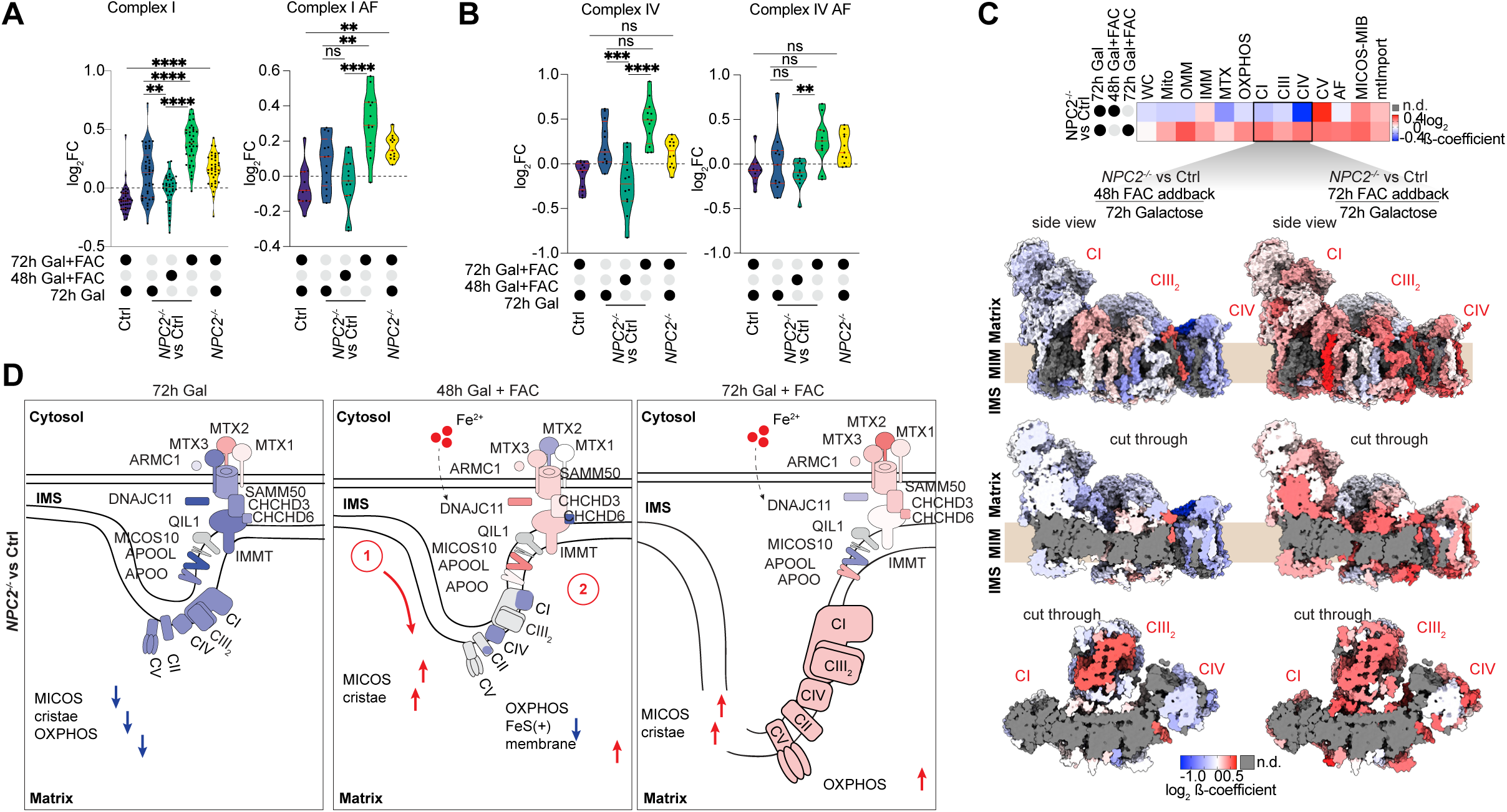
**Rescue of OXPHOS complex abundance in *NPC2*^-/-^ cells by extracellular iron**. **(A,B)** Violin plots of log_2_FC values for CI (panel D) and CIV (panel E) subunits (left panels) and associated assembly factors (right panels) in *NPC2*^-/-^ versus Control cells grown in Galactose with or without FAC. Complex I: p(****) < 0.0001, p(**) = 0.0036; Complex I AF: p(****) < 0.0001, p(**) = 0.0018; Complex IV: p(****) < 0.0001, p(***) = 0.0002; Complex IV AF: p(**) = 0.0056; ordinary one-way ANOVA with multiple comparisons, alpha = 0.05; data based on biological triplicate TMTpro measurements. **(C)** Log_2_FC of β-coefficient of mitochondrial components (see middle heatmap) in Galactose and either 48 or 72 h FAC addback for *NPC2*^-/-^ versus Control cells. Abundance of supercomplex subunits is mapped onto the structure (PDB: 5XTH). Vertical and horizontal cut throughs of the structure are depicted in the lower panels. Legend shows color panel for log_2_FC values. Data based on biological triplicate TMTpro measurements. **(D)** Schematic model for the rescue of mitochondrial cristae and OXPHOS complexes upon FAC addback in *NPC2*^-/-^ cells. See text for details.

**Figure S7.**
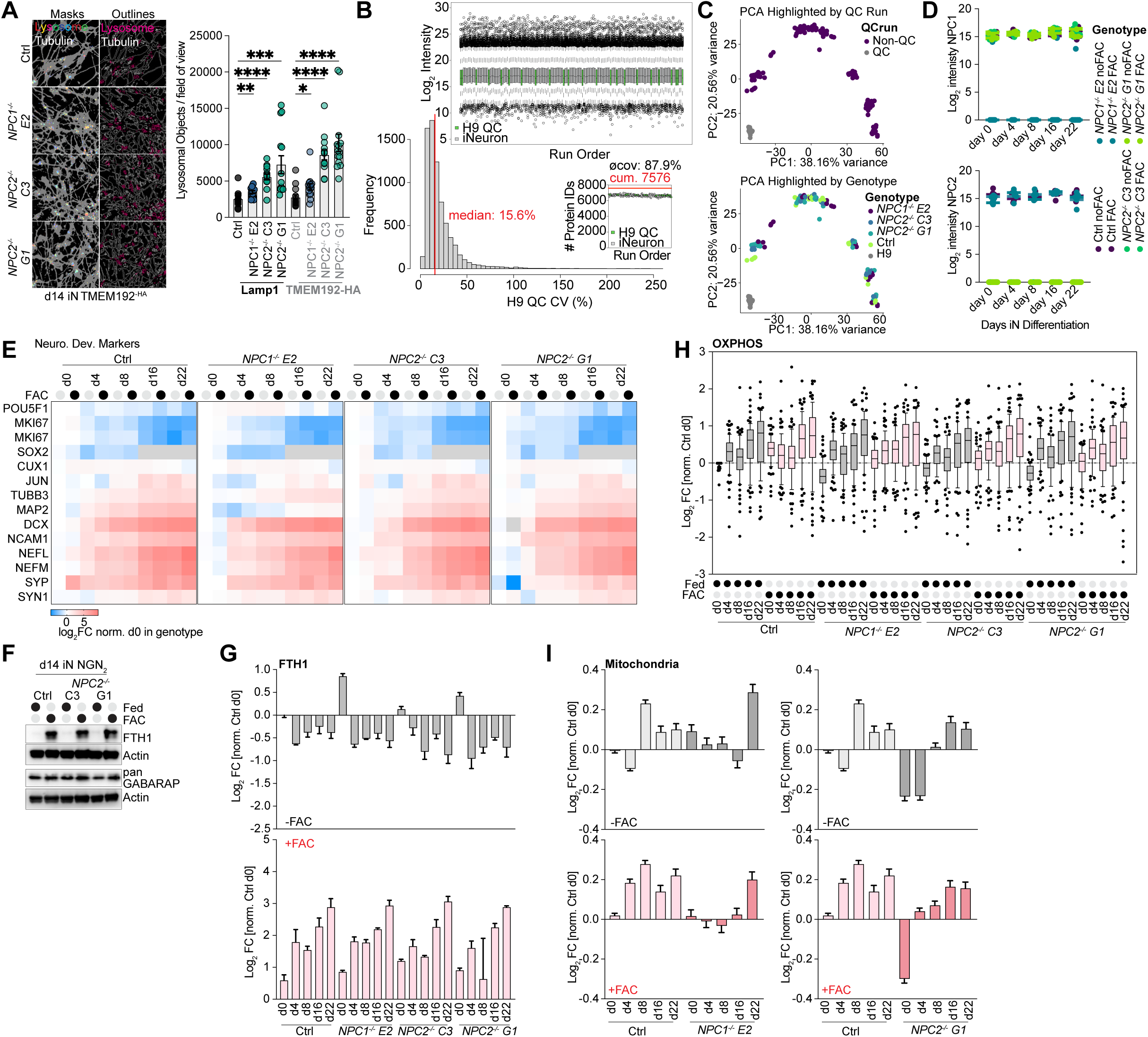
Neuronal proteomics of NPC mutants in presence of FAC. **(A)** Example object segmentation overlays (lysosome & tubulin) of day 14 iNeurons of the indicated genotypes. Quantification of lysosomal objects per stack for both α-LAMP1 and α-HA. Unpaired t.test Lamp1: *NPC1^-/-^*E2: p(**) = 0.0079, *NPC2^-/-^* C3: p(****) <0.0001, *NPC2^-/-^* G1: p(***) = 0.0009. HA: *NPC1^-/-^* E2: p(*) = 0.0167, *NPC2^-/-^* C3: p(****) <0.0001, *NPC2^-/-^* G1: p(****) < 0.0001. Data based 14 replicates. Error bars show S.E.M.. **(B)** QC-assessment of LFQ proteome data from iNeuron ± FAC. Boxplot of log_2_ intensity across the 142 LC-MS runs. Green boxplots depict H9 ESC QC samples, grey boxplots depict time-course samples (d0 – d22). Frequency distribution of H9 QC sample coefficient of variation (CV) across the 22 H9 ESC QC samples, covering the whole acquisition time-window. Average unique protein group coverage rate per run. On average 6659 protein IDs were detected. **(C)** PCA plot of LFQ data, color-coded according to run or genotype. **(D)** Log_2_ intensity for NPC1 (top) or NPC2 (bottom) across all time-points for the indicated genotypes. **(E)** Heatmap of neuronal development markers (log_2_FC norm. within genotype) across all time-points for the indicated genotypes. **(F)** Western blot of FTH1 and panGABARAP from iNeurons at day 14 of differentiation of Control and two *NPC2^-/-^* clones ± FAC. **(G)** Bargraph of mean log_2_FC (normalized within genotype day 0) of FTH1 ± FAC across all time-points and genotypes. **(H)** Boxplot of log_2_FC (normalized to Control at day 0) mitochondrial OXPHOS-components across the indicated genotypes, differentiation times and ± FAC treatment. **(I)** Bargraph of mean log_2_FC (normalized within genotype day 0) of the mitochondrial proteome ± FAC across all time-points.

## SUPPLEMENTARY TABLES

Table S1. Generation of CRISPR-edited cell lines for interrogation of lysosomal storage disease gene function analysis. This file contains gRNA sequences as well as allele sequencing results for all edits examined. Relevant to Figure S1E, S1F.

Table S2. nMOST proteomic and lipidomic analysis of 33 LSD cell lines (total proteome). Related to Figure 2, Figure 3, S1G-I.

Table S3. nMOST Cross-Ome analysis of 33 LSD cell lines (whole cell). Related to Figure 2, 3A-C.

Table S4. nMOST proteomic and lipidomic analysis of Control, LIPA-/-, GAA-/-, NPC1-/- and NPC2-/- HeLaTMEM192-HA cells under fed and starved (EBSS) conditions. Related to Figure 4A, S2D-I, S4E-H, 5A-E, S5A-D.

Table S5. TMTpro-based proteomic analysis of Control and NPC2-/- HeLaTMEM192-HA with and without FAC addition. Relevant to Figure 5L, M; S5J-T, S6A-D.

Table S6. nDIA label-free whole cell proteomics of neuronal differentiation timecourse (d0, d4, d8, d16, d22) of H9TMEM192-HA NGN2 Control, NPC1-/- E2, NPC2-/- C3, NPC2-/- G1. Relevant to Figure 7, S7B-E, H-I.

Table S7. Source data file containing data tabulated data used in figures.

