## Supplementary material for "Global cellular proteo-lipidomic profiling of diverse lysosomal storage disease mutants using nMOST": Key Resources Table

| **REAGENT or RESOURCE** | **SOURCE** | **IDENTIFIER** |
| --- | --- | --- |
| Deposited data | | |
| BioPlex 3.0 | https://bioplex.hms.harvard.edu/ | doi: 10.1016/j.cell.2021.04.011. |
| CORUM protein complexes database (28.11.2022 Corum 4.1 release) | https://mips.helmholtz-muenchen.de/corum/ |  |
| STRINGDB (version 12.0) | https://string-db.org/cgi/download?sessionId=bxEPGb8BbQKu |  |
| UniProt (release 2021-11; 2024-01) | https://www.uniprot.org/ | RRID:SCR_002380 |
| Mitocarta 3.0 | https://personal.broadinstitute.org/scalvo/MitoCarta3.0/human.mitocarta3.0.html | RRID:SCR_018165 |
| nMOST Dataset (LSD and 4KO) | MassIVE | MSV000094201 |
| LSD-nMOST data online viewer | Coon Lab | <https://coonlabdatadev.com/>  Username: most_lsd and Password: viewer |
| raw files of TMTpro protemic data of HeLa Control, NPC1-/-, NPC2-/- in Glucose or Galactose conditions ± FAC | PRIDE | PXD049336 |
| Tabulated Data of TMTpro protemic data of HeLa Control, NPC1-/-, NPC2-/- in Glucose or Galactose conditions ± FAC | Supplemental Table, Zenodo folder | 10.5281/zenodo.13905094 |
| raw files of nDIA-LFQ of H9 Control, NPC1-/- and NPC2-/- during neurogenesis ±FAC | MassIVE | MSV000095985 |
| Tabulated Data of nDIA-LFQ of H9 Control, NPC1-/- and NPC2-/- during neurogenesis ±FAC | Supplemental Table, Zenodo folder | 10.5281/zenodo.13905094 |
| Scripts used for image evaluation | Zenodo folder | 10.5281/zenodo.13905094 |
| CryoET raw tomograms of HeLa TMEM192-3xHA Control and NPC2-/- | EMDB | EMD-51701, EMD-51702 |
| nMOST-LSD Protocol Collection | Protocols.io | dx.doi.org/10.17504/protocols.io.5qpvokmmzl4o/v1 |
| Software and algorithms | | |
| CHOPCHOP | http://chopchop.cbu.uib.no/ | RRID:SCR_015723 |
| Snapgene (v.5.1.7) | Dotmatics | https://snapgene.com (RRID:SCR_015052) |
| Illustrator (v28) | Adobe Inc. | RRID:SCR_010279 |
| Excel (v.16.81) | Microsoft Inc. | RRID:SCR_016137 |
| Prism (v10.3.1) | GraphPad | GraphPad Prism, (RRID:SCR_002798) |
| R (4.2.2) / RStudio (2022.07.01 Build 554) | R Project for Statistical Computing | https://www.rstudio.com/, (RRID:SCR_000432) |
| library(tibble) | v3.1.8 | https://cran.r-project.org/web/packages/tibble/tibble.pdf |
| library(ggVennDiagram) | v1.2.0 | https://cran.r-project.org/web/packages/ggVennDiagram/index.html |
| library(stringr) | v1.4.0 | RRID:SCR_022813 |
| library(dplyr) | v1.0.9 | RRID:SCR_016708 |
| library(ggsci) | v2.9 | https://github.com/nanxstats/ggsci |
| library(ggrepel) | v0.9.1 | RRID:SCR_017393 |
| library(ggplot2) | v3.3.6 | RRID:SCR_014601 |
| library(tidyverse) | v1.3.2 | RRID:SCR_019186 |
| library(timecourse) | v1.68.0 | RRID:SCR_000077 |
| library(msstats) | v4.10.0 | https://www.bioconductor.org/packages/release/bioc/html/MSstats.html, (RRID:SCR_014353) |
| library(purr) | v1.0.1 | https://www.rdocumentation.org/packages/purrr/versions/1.0.2, (RRID:SCR_021267) |
| library(tibble) | v3.1.8 | https://www.rdocumentation.org/packages/tibble/versions/3.2.1 |
| library(RColorBrewer) | v1.1-3 | https://www.rdocumentation.org/packages/RColorBrewer/versions/1.1-3, (RRID:SCR_016697) |
| library(pheatmap) | v.1.0.12 | https://www.rdocumentation.org/packages/pheatmap/versions/1.0.12/topics/pheatmap, (RRID:SCR_016418) |
| library(cowplot) | v.1.1.3 | https://www.rdocumentation.org/packages/cowplot/versions/1.1.3, (RRID:SCR_018081) |
| library(reshape2) | v1.4.4 | https://cran.r-project.org/web/packages/reshape2/index.html; reshape2 (RRID:SCR_022679) |
| library(viridis) | v0.6.5 | https://cran.r-project.org/web/packages/viridis/vignettes/intro-to-viridis.html ; viridis (RRID:SCR_016696) |
| library(car) | v3.1-2 | https://cran.r-project.org/package=car; https://cran.r-project.org/package=car |
| library(ggpubr) | v0.6.0 | https://cran.r-project.org/web/packages/ggpubr/index.html ; ggpubr (RRID:SCR_021139) |
| library(igraph) | v1.3.5 | https://cran.r-project.org/web/packages/igraph/index.html |
| ShinyGO | Ge et al., 2020 | http://bioinformatics.sdstate.edu/go/ |
| MassPike, In-house mass spectrometry data analysis software | Huttlin et al Cell. (2010) 143:1174-89. | https://gygi.hms.harvard.edu/software.html |
| MaxQuant (Version 2.0.3.0) | Cox et al, 2008 | MaxQuant (RRID:SCR_014485) |
| Monocle | Rad et al, 2021 | RRID:SCR_018685 |
| SEQUEST | Eng et al., 1994 | https://proteomicsresource.washington.edu/protocols06/sequest.php |
| Comet 2019.01 | Eng et al 2013 | http://comet-ms.sourceforge.net/; CoMet (RRID:SCR_011925) |
| Perseus (v1.6.5) | Tyanova et al., Nat Methods. (2016) 13:731-40. | http://www.perseus-framework.org, (RRID:SCR_015753) |
| LipiDex v1.1 | Hutchins et al., 2018 | https://github.com/coongroup/LipiDex |
| LipiDex Spectrum Annotator | Park et al., 2022 | https://github.com/coongroup/LipiDexSpectrumAnnotator |
| Compound Discoverer 3.1 | Thermo Fisher Scientific | RRID:SCR_014477 |
| MSConvert | Adusumilli et al., 2017 | http://www.proteowizard.org/download.html |
| DIA-NN | https://github.com/vdemichev/DiaNN | https://github.com/vdemichev/DiaNN |
| Fragpipe | https://github.com/Nesvilab/FragPipe | RRID:SCR_022864 |
| AmiGO | http://amigo.geneontology.org/ | AmiGO (RRID:SCR_002143) |
| Morpheus | https://morpheus.gitlab.io/ | Morpheus (RRID:SCR_014975) |
| CUDA | http://sourceforge.net/projects/cudasphere/ | CUDA-SPHERE-FWD-MEEG (RRID:SCR_013225) |
| Arctis WebUI | Spurný et al., 2023 | https://academic.oup.com/mam/article/29/Supplement_1/2081/7228167 |
| FiJi | ImageJ 1.53t30 | https://imagej.net/Fiji, (RRID:SCR_002285) |
| cudasirecon | Github | https://github.com/scopetools/cudasirecon |
| otfsearch | Github | https://github.com/tlambert03/otfsearch |
| CellProfiler (v.4.2.1) | Stirling DR et al., 2021 | https://cellprofiler.org, (RRID:SCR_007358) |
| NIS Elements | 5.21.3 (Build 1489) | https://www.microscope.healthcare.nikon.com/products/software/nis-elements, (RRID:SCR_014329) |
| TomoMAN Code | n/a | https://doi.org/10.5281/ZENODO.4110737 |
| IMOD (v.4.10.49) | n/a | https://bio3d.colorado.edu/imod/ (RRID:SCR_003297) |
| cryoCARE | n/a | https://github.com/juglab/cryoCARE_T2T |
| pandas (v.1.3.0) | n/a | https://pandas.pydata.org/ (RRID:SCR_018214) |
| matplotlib (v.3.3.0) | n/a | https://matplotlib.org/ (RRID:SCR_008624) |
| seaborn (v.0.11.0) | n/a | https://seaborn.pydata.org/ (RRID:SCR_018132) |
| Membrain-Seg (v9) | n/a | https://github.com/teamtomo/membrain-seg |
| Huygens | Huygens Software | Huygens Software (RRID:SCR_014237) |
| Tomo5 (v.5.12.0) | TomoPy | TomoPy (RRID:SCR_021359) |
| Motioncorr2 | MotionCor2 | MotionCor2 (RRID:SCR_016499) |
| Python (v.3.9.7) | Python Programming Language | Python Programming Language (RRID:SCR_008394) |
| BigWarp (v.9.0.0) | n/a | https://imagej.net/plugins/bigwarp |
| Arctis WebUI (1.0) | Thermo Fisher Scientific | 10.1093/micmic/ozad067.1077 |
| Amira | Thermo Fisher Scientific | https://www.thermofisher.com/us/en/home/electron-microscopy/products/software-em-3d-vis/amira-software.html, (RRID:SCR_007353) |
| AreTomo (v.1.3.3) | Zheng et al., 2022 | https://github.com/czimaginginstitute/AreTomo2 |
| EMAN (v. 2.99.47) | https://blake.bcm.edu/emanwiki/EMAN2 | EMAN (RRID:SCR_016867) |
| Gwyddion (v.2.63) | http://gwyddion.net/ | Gwyddion (RRID:SCR_015583) |
| Dragonfly (v.2022.2) | Comet Technologies Canada Inc. | https://www.theobjects.com/dragonfly/index.html, Dragonfly (RRID:SCR_025150) |
| OriginLab 2023 | OriginLab Corp. | https://www.originlab.com/2023, Origin (RRID:SCR_014212) |
| UCSF ChimeraX | version 1.4.dev202201050842 (2022-01-05) | https://www.cgl.ucsf.edu/chimerax/ (RRID:SCR_015872) |
| Antibodies | | |
| Anti-HSP60 | Abcam | ab128567; (RRID:AB_11145464) |
| Anti-HA High Affinity (from rat IgG1) | Roche | 11867423001; (RRID:AB_390918) |
| Anti-Tomm20 | Abcam | ab186735; (RRID:AB_2889972) |
| Anti-SQSTM | Abnova | H00008878-M01; (RRID:AB _437085) |
| Anti-LC3B (D11) | Cell Signaling Technology | 3868; (RRID:AB_915950) |
| Anti-GABARAP+GABARAPL1+GABARAPL2 | Abcam | ab109364; (RRID:AB_10861928) |
| Anti-Ferritin antibody FTH1 [EPR3004Y] | Abcam | ab75973; (RRID:AB_1310222) |
| Anti-LAMP1 (D2D11) XP Rabbit | Cell Signaling Technology | #9091S; (RRID:AB_2687579) |
| Anti-LAMP1 (D4O1S) Mouse | Cell Signaling Technology | #15665; (RRID:AB_2798750) |
| Anti-MTCO2 | Abcam | ab110258; (RRID:AB_10887758) |
| Anti-GFP | Thermo Fisher | a10262; (RRID:AB_2534023) |
| Anti-Tubulin (beta III ) antibody (chicken) | Abcam | ab41489; (RRID:AB_727049) |
| Anti-Actin | Sigma | A2228; (RRID:AB_476697) |
| Anti-NCOA4 | Cell Signaling | #66849; (RRID:AB_3064842) |
| Goat anti-Rabbit IgG (H+L) Highly Cross-Adsorbed Secondary Antibody, Alexa Fluor Plus 405 | Thermo Fisher Scientific | A48254; (RRID:AB_2890548) |
| Donkey anti-Rat IgG (H+L) Highly Cross-Adsorbed Secondary Antibody, Alexa Fluor Plus 405 | Thermo Fisher Scientific | A48268; (RRID:AB_2890549) |
| Goat anti-Mouse IgG (H+L) Cross-Adsorbed Secondary Antibody, Alexa Fluor 488 | Thermo Fisher Scientific | A-11001; (RRID:AB_2534069) |
| Goat anti-Rabbit IgG (H+L) Highly Cross-Adsorbed Secondary Antibody, Alexa Fluor™ 488 | Thermo Fisher Scientific | A-11034; (RRID:AB_2576217) |
| Goat anti-Chicken IgY (H+L) Secondary Antibody, Alexa Fluor™ 488 | Thermo Fisher Scientific | A-11039; (RRID:AB_2534096) |
| Goat anti-Rat IgG (H+L) Cross-Adsorbed Secondary Antibody, Alexa Fluor 555 | Thermo Fisher Scientific | A-21434; (RRID:AB_2535855) |
| Goat anti-Rabbit IgG (H+L) Cross-Adsorbed Secondary Antibody, Alexa Fluor 568 | Thermo Fisher Scientific | A-11011; (RRID:AB_143157) |
| Goat anti-Rat IgG (H+L) Cross-Adsorbed Secondary Antibody, Alexa Fluor™ 647 | Thermo Fisher Scientific | A-21247; (RRID:AB_141778) |
| Goat anti-Rabbit IgG (H+L) Cross-Adsorbed Secondary Antibody, Alexa Fluor 647 | Thermo Fisher Scientific | A27040; (RRID:AB_2536101) |
| Goat anti-Chicken IgY (H+L) Secondary Antibody, Alexa Fluor™ 647 | Thermo Fisher Scientific | A-21449; (RRID:AB_2535866) |
| Goat anti-Mouse IgG (H+L) Cross-Adsorbed Secondary Antibody, Alexa Fluor 647 | Thermo Fisher Scientific | A-21235, (RRID:AB_2535804) |
| anti-rabbit IgG horse radish peroxidase (HRP) conjugate | Bio-Rad | Cat# 170-6515, RRID:AB_11125142 |
| anti-mouse IgG HRP conjugate | Bio-Rad | Cat# 170-6516, RRID:AB_11125547 |
| Bacterial and virus strains | | |
| pAAVS1-TRE3G-NGN2 | Ordureau et al., 2020 | 10.1016/j.molcel.2019.11.013 |
| pET-NLS-Cas9-6xHis | Addgene | RRID:Addgene_62934 |
| px459 | Addgene | RRID:Addgene_62988 |
| hCAS9 plasmid | Addgene | RRID:Addgene_41815 |
| pEGFP-C1-SopF | Addgene | RRID:Addgene_137734 |
| Critical commercial assays | | |
| Recombinant SpCas9 | Zuris et al., 2015; This paper | N/A |
| *GeneArt Precision gRNA Synthesis Kit* | Thermo Fisher Scientific | A29377 |
| FastBreak Buffer | Promega Inc | V8571 |
| Pierce™ BCA Protein Assay Kit | Thermo Fisher Scientific | 23227 |
| FuGENE HD Transfection Reagent | Promega | E2311 |
| QIAquick PCR Purification Kit | Qiagen | 28104 |
| Rneasy Mini Kit | Qiagen | 74104 |
| Chemicals, peptides, and recombinant proteins | | |
| Urea | Sigma | U5378 |
| Dithiothreitol (DTT) | Gold Biotechnology | DTT25 |
| NuPAGE 8%, Bis-Tris, 1.0 mm, Midi Protein Gel, 12+2-well | Thermo Fisher Scientific | WG1001BOX |
| NuPAGE Novex® 4-12% Bis-Tris Midi Protein Gels, 20 well | Thermo Fisher Scientific | WG1402BOX |
| NuPAGE Novex® 4-12% Bis-Tris Midi Protein Gels, 26 well | Thermo Fisher Scientific | WG1403BOX |
| NuPAGE™ LDS Sample Buffer (4X) | Thermo Fisher Scientific | NP0008 |
| Revert 700 Total Protein Stain Kits for Western Blot Normalization | LI-COR | 926-11016 |
| Precision Plus Protein™ Kaleidoscope™ Prestained Protein Standards | BioRad | 1610395 |
| Immobilon-P Membrane, PVDF, 0.45µm, 26.5cm x3.75m roll | EMD Millipore | IPVH00010 |
| EPPS | Sigma-Aldrich | E9502 |
| Formic Acid | Sigma-Aldrich | 94318 |
| Isopropanol, Optima™LC/MS Grade | Thermo Fisher Scientific | A461-4 |
| cOmplete, EDTA-free Protease Inhibitor Cocktail | Milipore Sigma | 11873580001 |
| Acetonitrile, Optima™ LC/MS Grade | Thermo Fisher Scientific | A955-4 |
| Adenosine 5’ triphosphate, disodium, trihydrate (ATP) | Thermo-Fisher Scientific | 10326943 |
| Ammonium formate CHROMASOLV™ LC-MS Ultra | Honeywell | 14266 Fluka |
| 2-Chloroacetamide | Sigma-Aldrich | C0267 |
| n-Butanol | Thermo Fisher Scientific | A383SK-4 |
| Sep-Pak tC18 96-well Plate, 25 mg Sorbent per Well | Waters | 186002319 |
| Lys-C | Wako Chemicals | 129-02541 |
| PhosSTOP phosphatase inhibitor cocktail | Roche | 4906845001 |
| Water, Optima™ LC/MS Grade | Thermo Fisher Scientific | W64 |
| TCEP | Gold Biotechnology | TCEP2 |
| Trypsin | Promega | V511C |
| TMTpro 18plex Label Reagent | Thermo Fisher Scientific | A52045 |
| Tris (1 M), pH 8.0, RNase-free | Thermo Fisher Scientific | AM9855G |
| Silicon Dioxide R1/4 film | Quantifoil, Fisher Scientific | 50-192-7684 |
| SeraSil-Mag™ silica coated superparamagnetic beads 700nm | Cytiva | 29357374 |
| Nunc Cell-Culture Treated 12-well | Thermo Fisher Scientific | 150628 |
| Nunc Cell-Culture Treated 6-well | Thermo Fisher Scientific | 140685 |
| µ-Slide 8 Well, Glass Bottom - #1.5H Glass Bottom Coverslip | ibidi | 80807 |
| Marienfeld Precision cover glasses thickness No. 1.5H (tol. ± 5 μm) 18x18mm | VWR | 107032 |
| 35 mm Dish \| High Precision 1.5 Coverslip \| 14 mm Glass Diameter | MatTek | P35G-0.170-14-C |
| 6 Well glass bottom plate with high performance #1.5 cover glass | Cellvis | P06-1.5H-N |
| 12 Well glass bottom plate with high performance #1.5 cover glass | Cellvis | P12-1.5H-N |
| 24 Well glass bottom plate with high performance #1.5 cover glass | Cellvis | P24-1.5H-N |
| 96 Well glass bottom plate with high performance #1.5 cover glass | Cellvis | P96-1.5H-N |
| 100x21mm Dish, Nunclon Delta | Thermo Fisher Scientific | 172931 |
| 150mm plates (15cm) | MidSci | TP93150 |
| Filipin complex from Streptomyces filipinensis | Sigma | F9765 |
| SiR-Lysosome Kit | Cytosleleton / Spirochrome | CY-SC012 |
| Hoechst33342 | Thermo Fisher Scientific | H1399 |
| SPY555-DNA | Cytosleleton / Spirochrome | CY-SC201 |
| PKmitoRed | Cytosleleton / Spirochrome | CY-SC052 |
| MitoTrackerDeepRed FM | Thermo Fisher Scientific | M22426 |
| Tetramethylrhodamine, Ethyl Ester, Perchlorate (TMRE) | Thermo Fisher Scientific | T669 |
| Vectashield | Vector Laboratories | H-1000-10 |
| Lysosomal pH Detection Reagent Red (3 tubes) | Dojindo | L265-12 |
| Oregon Green 488 BAPTA-5N, Hexapotassium Salt, cell impermeant | Thermo Fisher Scientific | O6812 |
| Oregon Green 488 BAPTA-1, AM, cell permeant | Thermo Fisher Scientific | O6807 |
| LysoTrackerRed DND-99 | Thermo Fisher Scientific | L7528 |
| Paraformaldehyde 3% Glutaraldehyde 0.35% in 0.1M Sodium Cacodylate, pH 7.4 | Electron Microscopy Science | 15949-50 |
| 16% Paraformaldehyde, Elecron-Mircoscopy Grade | Electron Microscopy Science | 15710 |
| Cover Glasses | VWR | 16004-308 |
| Dextran, Alexa Fluor™ 647; 10,000 MW, Anionic, Fixable | Thermo Fisher Scientific | D22914 |
| DAPI (4',6-Diamidino-2-Phenylindole, Dihydrochloride) | Thermo Fisher Scientific | D1306 |
| DMEM, High Glucose, Pyruvate | Thermo Fisher Scientific | 11995-073 |
| DMEM, no glucose | Thermo Fisher Scientific | 11966025 |
| DMEM/F12 | Thermo Fisher Scientific | 11330057 |
| Doxycycline | Sigma-Aldrich | D9891 |
| EBSS | Thermo Fisher Scientific | 14155063 |
| Monensin | Cayman Chemicals | 16488 |
| Uridine | Sigma | U3003 |
| Puromycin | Gold Biotechnology | P-600-500 |
| Trypsin-EDTA | Sigma | T4049-100ML |
| Sodium Pyruvate | Invitrogen | 11360070 |
| PBS | Corning | 21-031-CV |
| Penicillin-Streptomycin (10,000 U/mL) | Thermo Fisher Scientific | 15140163 |
| OptiMEM I reduced serum media | Thermo Fisher Scientific | 31985062 |
| Lipofectaime LTX | Thermo Fisher Scientific | 15338100 |
| GlutaMAX | Thermo Fisher Scientific | 35050061 |
| SAR405 Selective ATP-competitive inhibitor of Vps34 | APExBio | A8883 |
| ML-SA5 | MedChemExpress | HY-152182 |
| Torin 1 | Sellekchem | S2827 |
| U18666A | MedChemExpress | HY-107433 |
| Accutase | StemCell | 7920 |
| Insulin Human | Sigma-Aldrich | I9278-5ML |
| holo-Transferrin human | Sigma-Aldrich | T0665 |
| Hygromycin B | Thermo Fisher Scientific | 10687010 |
| MEM NEAA | Thermo Fisher Scientific | 11140050 |
| N-2 Supplement (100X) | Thermo Fisher Scientific | 17502048 |
| B27 | Thermo Fisher Scientific | 17504001 |
| Geltrex LDEV-Free Reduced Growth Factor Basement Membrane Matrix | Thermo Fisher Scientific | A1413202 |
| NEAA | Life Technologies | 11140050 |
| Neurobasal | Thermo Fisher Scientific | 21103049 |
| Neurotrophin-3 | Peprotech | 450-03 |
| Sodium selenite | Sigma-Aldrich | S5261-10G |
| Sodium Bicarbonate | Sigma-Aldrich | S5761-500G |
| TGF-beta | PeproTech | 100-21C |
| Y-27632 Dihydrochloride (ROCK inhibitor) | PeproTech | 1293823 |
| Brain-derived neurotrophic factor (BDNF) | Peprotech | 450-02 |
| UltraPure 0.5M EDTA, pH 8.0 | Thermo Fisher Scientific | 15575020 |
| Cultrex 3D Culture Matrix Laminin I | R&D Systems | 3446-005-01 |
| FGF3 | in-house | N/A |
| FluoroBrite DMEM | Thermo Fisher Scientific | A1896701 |
| Biological samples | | |
| HEK293T | ATCC | RRID:CVCL_0045 |
| HeLa TMEM192-3xHA | Boland et al. 2022 | RRID:CVCL_D1KR |
| HeLa TMEM192-3xHA ARSB-/- | This study | RRID:CVCL_D5EJ |
| HeLa TMEM192-3xHA ASAH1-/- | This study | RRID:CVCL_D5EK |
| HeLa TMEM192-3xHA CLN3-/- | This study | RRID:CVCL_D5EL |
| HeLa TMEM192-3xHA CTSD-/- | This study | RRID:CVCL_D5EM |
| HeLa TMEM192-3xHA GAA-/- | This study | RRID:CVCL_D5EN |
| HeLa TMEM192-3xHA GALNS-/- | This study | RRID:CVCL_D5EP |
| HeLa TMEM192-3xHA GLB1-/- | This study | RRID:CVCL_D5EQ |
| HeLa TMEM192-3xHA GNS-/- | This study | RRID:CVCL_D5ER |
| HeLa TMEM192-3xHA GRN-/- | Boland et al. 2022 | RRID:CVCL_D5ES |
| HeLa TMEM192-3xHA GUSB-/- | This study | RRID:CVCL_D5ET |
| HeLa TMEM192-3xHA HEXA-/- | Boland et al. 2022 | RRID:CVCL_D5EU |
| HeLa TMEM192-3xHA HEXB-/- | This study | RRID:CVCL_D5EV |
| HeLa TMEM192-3xHA IDUA-/- | This study | RRID:CVCL_D5EW |
| HeLa TMEM192-3xHA KCTD7-/- | This study | RRID:CVCL_D5EX |
| HeLa TMEM192-3xHA LAMP2-/- | This study | RRID:CVCL_D5EY |
| HeLa TMEM192-3xHA LIPA-/- | This study | RRID:CVCL_D5EZ |
| HeLa TMEM192-3xHA MAN2B1-/- | This study | RRID:CVCL_D5F0 |
| HeLa TMEM192-3xHA MFSD8-/- | This study | RRID:CVCL_D5F1 |
| HeLa TMEM192-3xHA NAGLU-/- | This study | RRID:CVCL_D5F2 |
| HeLa TMEM192-3xHA NPC1-/- | This study | RRID:CVCL_D5F3 |
| HeLa TMEM192-3xHA NPC2-/- | This study | RRID:CVCL_D5F4 |
| HeLa TMEM192-3xHA PPT1-/- | This study | RRID:CVCL_D5F5 |
| HeLa TMEM192-3xHA PSAP-/- | This study | RRID:CVCL_D5F6 |
| HeLa TMEM192-3xHA CLN6-/- | This study | RRID:CVCL_D5FC |
| HeLa TMEM192-3xHA CTSA-/- | This study | RRID:CVCL_D5FD |
| HeLa TMEM192-3xHA GBA-/- | This study | RRID:CVCL_D5FE |
| HeLa TMEM192-3xHA GLA-/- | This study | RRID:CVCL_D5FF |
| HeLa TMEM192-3xHA GM2A-/- | This study | RRID:CVCL_D5FG |
| HeLa TMEM192-3xHA GNPTG-/- | This study | RRID:CVCL_D5FH |
| HeLa TMEM192-3xHA SCARB2-/- | This study | RRID:CVCL_D5FI |
| HeLa TMEM192-3xHA SGSH-/- | This study | RRID:CVCL_D5FJ |
| HeLa TMEM192-3xHA SMPD1-/- | This study | RRID:CVCL_D5FK |
| HeLa TMEM192-3xHA TPP1-/- | This study | RRID:CVCL_D5FL |
| H9 ES cells + AAVS-NGN2 + TMEM192-3xHA | Hickey et al., 2023 | RRID:CVCL_D1KT |
| H9 ES NPC1^-/-^ + AAVS-NGN2 + TMEM192-3xHA | This paper | RRID:CVCL_E3BP |
| H9 ES NPC2^-/-^ + AAVS-NGN2 + TMEM192-3xHA | This paper | RRID:CVCL_E3BQ |
| Oligonucleotides | | |
| Primers for 5’ AAVS junction PCR | 5’-ctctaacgctgccgtctctc-3’ and 5’-tgggcttgtactcggtcatc-3’ | N/A |
| Primers for 3’ AAVS junction PCR | 5’-cacacaacatacgagccgga-3’ and 5’-accccgaagagtgagtttgc-3’ | N/A |
| Primes for MiSeq KO validation and gRNAs can be found in Table S1 |  | N/A |
| Other | | |
| Orbitrap Fusion Lumos Tribid Mass Spectrometer | ThermoFisher Scientific | RRID:SCR_020562 |
| Orbitrap Eclipse Tribrid Mass Spectrometer | ThermoFisher Scientific | RRID:SCR_020559 |
| Orbitrap Exploris 480 Mass Spectrometer | ThermoFisher Scientific | RRID:SCR_022215 |
| Easy-nLC 1200 | ThermoFisher Scientific | https://www.thermofisher.com/order/catalog/product/LC120, (RRID:SCR_014993) |
| UltiMate 3000 UHPLC | ThermoFisher Scientific | https://www.thermofisher.com/us/en/home/industrial/chromatography/liquid-chromatography-lc/hplc-uhplc-systems/ultimate-3000-hplc-uhplc-systems.html |
| Vanquish™ Neo UHPLC System | ThermoFisher Scientific | https://www.thermofisher.com/order/catalog/product/VN-S10-A-01 |
| ACQUITY UPLC BEH C18 Column, 130Å, 1.7 µm, 1 mm X 150 mm, 1/pk | Waters | 186002347 |
| Aeris™ 2.6 μm PEPTIDE XB-C18 100 Å 250 x 4.6 mm | Phenomenex | 00G-4505-E0 |
| Sep-Pak C18 1cc Vac Cartridge, 50 mg | Waters | WAT054960 |
| StrataX 10 mg 96-well Plate | Phenomenex | 8E-S100-AGB |
| Illumina MiSeq | Illumina | RRID:SCR_016379 |
| SOLA HRP SPE Cartridge, 10 mg | Thermo Fisher Scientific | 60109-001 |
| Empore™ SPE Disks C18 | 3M Bioanalytical Technologies | 2215 |
| Nikon Ti w/ Yokagawa CSU-W1 spinning disk confocal | Nikon Corporation | https://www.microscope.healthcare.nikon.com/products/confocal-microscopes/csu-series/csu-w1, (RRID:SCR_021242) |
| Nikon Ti w/ Yokagawa CSU-X1 spinning disk confocal | Nikon Corporation | https://www.microscope.healthcare.nikon.com/products/confocal-microscopes/csu-series/csu-x1, (RRID:SCR_021242) |
| DeltaVision OMX V4 | Applied Precision (GE Healthcare) | https://sim.hms.harvard.edu/tech-specs/, (RRID:SCR_019956) |
| SoftWoRx | Applied Precision (GE Healthcare) | http://incelldownload.gehealthcare.com/bin/download_data/SoftWoRx/7.0.0/SoftWoRx.htm, (RRID:SCR_019157) |
| Krios G4 Cryo-TEM | Thermo Fisher Scientific | https://www.thermofisher.com/us/en/home/electron-microscopy/products/transmission-electron-microscopes/krios-cryo-tem.html, (RRID:SCR_019887) |
| Arctis Cryo-Plasma-FIB | Thermo Fisher Scientific | https://www.thermofisher.com/us/en/home/electron-microscopy/products/dualbeam-fib-sem-microscopes/arctis-cryo-pfib.html |
| Arctis WebUI | Spurny et al 2023, 10.1093/micmic/ozad067.1077 | https://academic.oup.com/mam/article/29/Supplement_1/2081/7228167 |
| Vitrobot Mark IV | Thermo Fisher Scientific | https://www.thermofisher.com/us/en/home/electron-microscopy/products/sample-preparation-equipment-em/vitrobot/instruments/vitrobot-mark-iv.html |
